# A specialized extracellular matrix patterns the conserved cortical protein EFA-6 at neuron-epidermal interfaces

**DOI:** 10.64898/2026.11.20.768229

**Authors:** Anjali Sandhu, Xiaohui Lyu, Jianing Shi, Xinghaoyun Wan, Jing Xie, Kancheng Yin, Ngang Heok Tang, Andrew D Chisholm, Yishi Jin

## Abstract

Sensory neurons establish intimate connections with the surrounding epidermis, where specialized extracellular matrix (ECM) help maintain neuron-epidermal interactions. However, whether and how this specialized ECM regulates the spatial organization of cortical proteins at these interfaces remains poorly understood. Here, we show a specialized ECM patterns the cortical microtubule regulator EFA-6 at neuron-epidermal interfaces in *Caenorhabditis elegans*. We show that EFA-6 in the epidermal cortex is enriched adjacent to the neurites of touch receptor neurons, characterized by discrete foci at the distal neurite and a diffuse distribution in the proximal neurite. We identify the specialized ECM components MEC-1 and HIM-4/hemicentin as regulators of EFA-6 spatial patterning at the interface between epidermis and neuronal processes. EFA-6 proteins contain an intrinsically disordered region (IDR) that is necessary and sufficient for EFA-6 targeting to and patterning at the neuron-epidermal interface. Within the IDR, we further identify an 8 residue motif that regulates enrichment and patterning of EFA-6 IDR at this interface. Finally, we find that EFA-6 contributes to age-dependent neurite defects in *him-4* mutants. Together, our findings reveal how a specialized extracellular matrix and an intrinsically disordered region coordinate the spatial organization of a cortical MT regulator at neuron-epidermal interface.

**Highlights:**

- The cortical MT regulator EFA-6 exhibits distinct spatial patterning along ALM neurites
- Epidermal EFA-6 is selectively enriched at neuron-epidermal interfaces through its intrinsically disordered region (IDR)
- A short sequence motif within the EFA-6 IDR is essential for interface-specific enrichment
- EFA-6 patterning at neuron-epidermal interface is regulated by extracellular matrix protein hemicentin/HIM-4

## INTRODUCTION

Sensory neurons establish intimate associations with surrounding epidermal cells, forming specialized interfaces that support neuronal development and function. These associations range from neurite extending along the epidermal surface to those becoming enclosed within epidermal invaginations. In Drosophila, integrin-mediated adhesion regulates positioning of sensory neurites along the basal epidermal surface while the membrane-associated protein Coracle promotes their enclosure by epidermal cells and restricts excessive dendritic branching (Kim et al., 2012; Tenenbaum et al., 2017). Similar epidermal ensheathment of sensory neurites has been observed in zebrafish, indicating that these associations occur across species (Kottmeier et al., 2020; O’Brien et al., 2012). In *Caenorhabditis elegans*, epidermal adhesion proteins also play important roles in sensory dendrite patterning. SAX-7/L1CAM and MNR-1/Menorin interact with the neuronal receptor DMA-1 to direct the stereotyped branching of PVD sensory dendrites (Dong et al., 2013). The spatial distribution of epidermal SAX-7 is further regulated by the basement membrane protein UNC-52/Perlecan and associated hemidesmosomal structures (Liang et al., 2015). These studies highlight how extracellular interactions regulate membrane proteins spatial distribution and dendritic positioning. In *C. elegans*, touch receptor neurons (TRN) extend long processes along the body wall that become progressively embedded within epidermis during development (Chalfie & Sulston, 1981). These neurites are associated with a specialized extracellular matrix (ECM), referred to as the mechanosensory ECM (mECM) or mantle, which consists of unique molecular composition and plays structural roles distinct from basement membranes present elsewhere (Du et al., 1996; Emtage et al., 2004). mECM protects neurite from mechanical stress induced by movement of the animal (Bonacossa-Pereira et al., 2022; Das et al., 2024). Specialized mECM components include EGF and Kunitz domains containing proteins MEC-1 and MEC-9, and collagen MEC-5, which are essential for organization of mechanosensory complexes and neuronal attachment (Das et al., 2024; Emtage et al., 2004). mECM also include HIM-4/hemicentin, a large conserved extracellular matrix protein involved in tissue attachment and extracellular organization (Dong et al., 2006; Gianakas et al., 2023; Lin et al., 2020; Morrissey et al., 2014; Vogel & Hedgecock, 2001; Welcker et al., 2021). More recently, laminin and nidogen were shown to associate with MEC-4/degenerin mechanosensory channels at these interfaces, demonstrating that the specialized ECM also contributes to the assembly of membrane-associated protein complexes (Das et al., 2024). Together, these studies establish that specialized ECM maintain neuron-epidermal interactions and organize neuronal membrane-associated proteins. However, whether this ECM also regulates the spatial distribution of proteins at the adjacent epidermal cortex remains unclear.

Mechanisms linking ECM engagement to cytoskeletal organization have been extensively investigated, particularly in the context of integrin-mediated focal adhesions. At these sites, adaptor proteins such as talin connect integrins to the actin cytoskeleton while vinculin reinforces these connections in response to mechanical forces (Calderwood et al., 1999; Grashoff et al., 2010). Focal adhesions also influence cortical microtubules (MT) organization through scaffolds such as KANK1, which associates with talin and promotes the recruitment of cortical MT-associated complexes containing CLASPs, LL5β, and liprins (Bouchet et, 2016; Lansbergen et al., 2006). Recent studies have identified phase separation as an additional mechanism for organizing proteins at adhesion sites. For example, LIMD1 and tensin3 undergoes phase separation to promote the assembly of adhesion-associated complexes (Li et al., 2026; Liang et al., 2024). Similarly, KANK1 forms condensates that facilitate the recruitment and organization of cortical MT-associated proteins (Aureille et al., 2024; Guo et al., 2023; Wang et al., 2021). These studies highlight how ECM and associated adhesion structures influence recruitment and spatial organization of cortical proteins. However, whether specialized ECM similarly regulates the spatial distribution of cortical proteins, especially MT regulators, at neuron-epidermal interfaces in vivo remain poorly studied.

EFA6/EFA-6 proteins are conserved regulators of cortical MT dynamics that restrict MT growth at the cell periphery (Bu et al., 2021; Chen et al., 2011; Chen et al., 2015; Eva et al., 2017; O’Rourke et al., 2010; Qu et al., 2019). In *C. elegans*, EFA-6 regulates axon growth during development and regeneration. EFA-6 contains a microtubule elimination domain (MTED), an intrinsically disordered region (IDR), and several conserved structured domains, including Sec7, pleckstrin homology (PH), and coiled-coil (CC) domains. These domains provide multiple potential mechanisms for coupling MT regulation to membrane localization and protein interactions. Our previous work found that EFA-6 is expressed in multiple tissues, including epithelial cells, neurons, germ cells. We demonstrated that EFA-6 undergoes IDR-dependent phase separation in vitro and forms discrete foci at the epithelial cortex. EFA-6 cortical organization is further influenced by interactions with MTs (Sandhu et al., 2024). These findings established a connection between the biophysical organization of EFA-6 and its cortical MT-regulatory function. However, how EFA-6 is spatially organized at neuron-epidermal interfaces and if extracellular matrix influences its spatial distribution and molecular dynamics, remains unknown.

Intrinsically disordered regions are increasingly recognized as important determinants of protein localization and organization within cells. Unlike structured protein domains, IDRs lack a stable three-dimensional fold and can mediate dynamic, multivalent interactions through short sequence motifs and distributed interaction sites (Banani et al., 2016; Cruz et al., 2024; Langstein-Skora et al., 2026; Li et al., 2012). Such properties enable IDRs to participate in molecular recognition, regulate protein association with cellular compartments, and promote the formation of biomolecular condensates (Banjade & Rosen, 2014; Lacy et al., 2004; Pombo-García et al., 2024; Shelby et al., 2023; Wang et al., 2023). Despite the recent advances, the contribution of IDRs to selective recognition of specialized cell-cell interfaces is less well understood. In particular, whether sequences embedded within an IDR can encode spatial specificity and how surrounding disordered sequences influence subcellular targeting efficiency under physiological conditions, remain important questions.

Here, we investigate how *C. elegans* EFA-6 is spatially organized at the interface between mechanosensory neurons and epidermis. We find that epidermal EFA-6 exhibits a striking proximal-distal distribution along ALM neurites, characterized by discrete distal foci and a diffuse proximal distribution. This patterning is developmentally regulated and correlates with neurite ensheathment by epidermal epithelia. Through an unbiased genetic screen, we identify ECM molecules MEC-1 and HIM-4 as regulators of EFA-6 patterning at the neuron-epidermal interface. EFA-6 IDR is essential and sufficient for selective enrichment at neuron-epidermal interfaces. Systematic dissection of the IDR identifies an eight-amino-acid motif that can recapitulate the spatial pattern of EFA-6, while surrounding disordered sequences promote efficient interface targeting. We further demonstrate that EFA-6 contributes to age-dependent neurite defects in *him-4* mutants. Together, our findings reveal how ECM organization influences the subcellular distribution of a cytoskeletal regulator.

## RESULTS

### EFA-6 exhibits distinct spatial patterning and protein dynamics along ALM anterior neurites

To investigate how EFA-6 is patterned at neuron-epidermal interfaces, we examined expression of endogenously GFP-tagged EFA-6 (GFP::EFA-6) (Sandhu et al., 2024) in TRNs. In day 1 adults, GFP::EFA-6 fluorescence was most prominently associated with the ALM and PLM neurons, located along the lateral body wall (Fig. S1A). GFP::EFA-6 displayed a predominantly uniform distribution with occasionally small visible foci along approximately 300 µm of the ALM neurite extending anteriorly from the cell soma, hereafter referred to as the proximal region (Fig. 1A, B; Fig. S1B). Toward the distal neurite, this diffuse distribution of GFP::EFA-6 progressively transitioned into discrete foci that became increasingly prominent and frequent. Within the distal most ∼30 µm of the neurite, or distal neurite tip, GFP::EFA-6 was observed almost exclusively as discrete foci (Fig. 1B, C). GFP::EFA-6 fluorescence intensity was also lower in the proximal region than at the distal tip (Fig. 1D). GFP::EFA-6 also exhibited a predominantly diffuse distribution along the PLM neurite and around the cell soma, transitioning into discrete foci within the distal most ∼50 µm of the neurite (Fig. S1C). Together, these observations reveal that EFA-6 is patterned along the proximo-distal axes of ALM and PLM neurites. Because this pattern was more clear-cut in ALM neurons, we focused analyses on the ALM anterior neurite.

**Figure 1:**
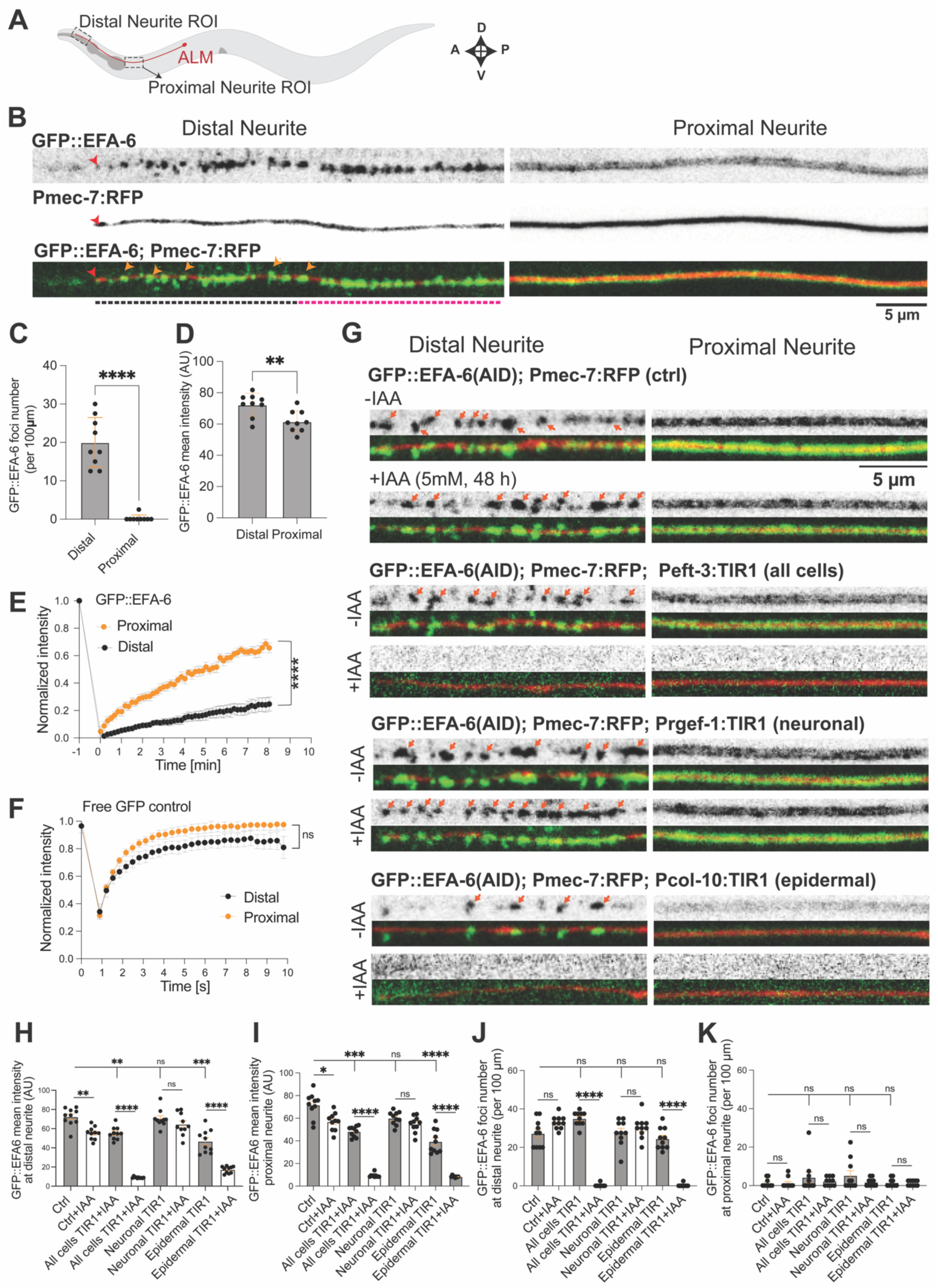
Epithelial EFA-6 is spatially patterned along ALM neurites. **(A)** Cartoon showing the distal and proximal neurite regions of interest (ROIs; indicated by black dotted rectangles) along the ALM neurite (marked in red) imaged in this study. **(B)** Representative single-plane confocal images of endogenous GFP::EFA-6 (KI) at the distal and proximal ROIs of the ALM neurite in adult animals. Within the distal ROI (∼40 µm), EFA-6 forms discrete foci over the distal-most ∼20 µm (marked by black dotted line) and transitions to a diffuse distribution proximally (marked by pink dotted line). In the proximal ROI, EFA-6 exhibits a predominantly uniform distribution. Pmec-7:RFP labels the ALM neurite. Red arrowheads mark the tip of neurite and orange arrowhead marks GFP::EFA-6 foci. Orange arrows mark EFA-6 foci. n = 10 animals per ROI. Scale bar, 5 µm. **(C, D)** Quantification of GFP::EFA-6 (C) foci density and (D) mean fluorescence intensity at the distal and proximal ROIs of ALM neurite in adult animals. **(E, F)** Fluorescence recovery after photobleaching (FRAP) analysis of (E) GFP::EFA-6 and (F) free GFP (Pmec-4:GFP) at the distal and proximal ROIs of ALM neurites (same as in panel (A)). **(G)** Representative single-plane confocal images of endogenous AID-tagged GFP::EFA-6 (KI) at the distal and proximal ROIs of ALM neurites (same in panel (A)) in L4 animals before and after indole-3-acetic acid (IAA; auxin) treatment for 48 h. The upper panels show the GFP channel alone, whereas the lower panels show merged GFP and RFP channels. Auxin treatment reduced GFP::EFA-6 levels at both distal and proximal ALM regions in animals expressing TIR1 in all cells (Peft-3:TIR1). Animals expressing neuronal TIR1 (Prgef-1:TIR1) showed little to no change in GFP::EFA-6 levels following auxin treatment. In contrast, epithelial TIR1 expression (Pcol-10:TIR1) resulted in a marked reduction in GFP::EFA-6 at both regions. Orange arrows mark EFA-6 foci. n = 30 animals per genotype. Scale bar, 5 µm. **(H-I)** Quantification of GFP::EFA-6 mean fluorescence intensity at the distal (H) and proximal (I) ROIs of ALM neurite (same as panel (A)) in L4 animals before and after auxin treatment. **(J-K)** Quantification of GFP::EFA-6 foci density at the distal (J) and proximal (K) at the distal and proximal ROIs of ALM neurites (same in panel (A)) in L4 animals before and after auxin treatment. Graphs show number foci per 100 µm length of neurite. **(C-F, H-K)** n=10-15 animals per genotype; Data are presented as mean ± SEM. Statistics: Unpaired t-test in (C, D), RM two-way ANOVA with the Geisser Greenhouse correction for (E, F), and Brown-Forsythe and Welch ANOVA tests for (H-K); ns, not significant (P > 0.05); *P < 0.05; **P < 0.01; ***P < 0.001; ****P < 0.0001.

We next asked whether EFA-6 patterning was associated with differences in its mobility. To address this, we performed fluorescence recovery after photobleaching (FRAP) analysis of GFP::EFA-6 along the ALM neurite (Fig. 1E). GFP::EFA-6 exhibited substantial fluorescence recovery in the proximal region, where it was predominantly diffusely distributed. In contrast, EFA-6 foci at the distal neurite tip showed significantly reduced recovery. In comparison, freely diffusible GFP expressed in neurons exhibited comparable fluorescence recovery in both regions (Fig. 1F). Together, these findings demonstrate that EFA-6 exhibits distinct spatial patterning along TRN neurites, accompanied by region-specific differences in protein mobility between the distal and proximal regions.

### EFA-6 spatial patterning along ALM neurites is developmentally regulated and correlates with neurite ensheathment

We next asked how GFP::EFA-6 spatial patterning occurred during development. In early larval stages, ALM neurites lie adjacent to both the epidermis and body wall muscle (BWM). During larval growth, ALM neurites become progressively ensheathed within the epidermal epithelium and displaced away from the BWM (Chalfie M, 1981; Du et al., 1996; Emtage et al., 2004). To investigate the relationship between neurite ensheathment and EFA-6 spatial organization, we examined GFP::EFA-6 localization at successive developmental stages while labeling BWM and ALM neurites with mCherry and RFP, respectively. We used the spatial separation between ALM neurites and BWM as an indicator of neuronal ensheathment (Fig. S2). In early L2 larvae, when the ALM neurite is adjacent to BWM, GFP::EFA-6 formed discrete foci along the entire length (Fig. S2A). By the L3 stage, the proximal ALM neurite began separating from BWM, and GFP::EFA-6 exhibited a more continuous distribution in these regions. In contrast, GFP::EFA-6 retained punctate distribution in the distal neurite, which remained adjacent to BWM (Fig. S2B). By the L4 stage, the proximal ALM neurite was fully ensheathed within the epidermis, as indicated by clear separation between the neurite and BWM. At this stage, GFP::EFA-6 exhibited a uniform distribution along the ensheathed proximal region while remaining punctate at the distal neurite tip (Fig. S2C), similar to adult animals.

Together, these observations reveal that EFA-6 spatial patterning undergoes a progressive developmental transition from discrete foci to a diffuse distribution in proximal neurite, correlating with ALM neurite ensheathment.

### EFA-6 localization along ALM neurites predominantly originates from the epidermis

Because ALM neurites are closely associated with the surrounding epidermis and EFA-6 is expressed in both neurons and epidermal cells (Chen et al., 2015; Sandhu et al., 2024), we next determined the relative contributions of these tissues to EFA-6 spatial patterning along ALM neurites. We first employed the auxin-inducible degron (AID) system for tissue-specific protein depletion. We generated an endogenous GFP::EFA-6(AID) knock-in strain by introducing an AID tag into the existing GFP::EFA-6 allele (Fig. S3A, B; Methods; Table S1, 2). To test the efficacy of AID-mediated EFA-6 depletion, we first used a transgene expressing E3 ligase TIR1 in all cells under the *eft-3* promoter (Zhang et al., 2015). Following auxin treatment for 48 h, GFP::EFA-6 fluorescence was abolished along the entire ALM neurites (Fig. 1G-K), demonstrating efficient depletion of the tagged protein. We next depleted EFA-6 specifically in neurons using pan-neuronal TIR1 transgene (Ashley et al., 2021). We verified the effectiveness of neuronal GFP::EFA-6 depletion by examining GFP::EFA-6 expression in the nerve ring and head ganglia. Auxin treatment markedly reduced GFP::EFA-6 fluorescence in these neuronal structures (Fig. S4A, B). Moreover, neuronal EFA-6 depletion significantly increased PLM neurite overgrowth (Fig. S4C), consistent with the previously reported cell-autonomous requirement for EFA-6 in TRN neurite growth (Chen et al., 2011; Chen et al., 2015). However, neuronal depletion did not detectably alter GFP::EFA-6 fluorescence intensity or spatial patterning along ALM neurites (Fig. 1G-K). In contrast, epidermal EFA-6 depletion significantly reduced GFP::EFA-6 fluorescence along ALM neurites, including the diffuse proximal signal and distal foci (Fig. 1G-K). Epidermal EFA-6 depletion did not reduce GFP::EFA-6 fluorescence in the nerve ring or head ganglia (Fig. S4A, B), supporting the tissue specificity of GFP::EFA-6 depletion. Epidermal EFA-6 depletion also did not affect PLM neurite overgrowth (Fig. S4C). These data establish that epidermal and neuronal EFA-6 have distinct localization and function.

To further test whether epidermal EFA-6 is sufficient to reproduce the observed spatial pattern, we generated a single-copy transgene expressing mNeonGreen (mNG)-tagged EFA-6 specifically in the epidermis in *efa-6* null animals (methods, Table S1, 2). Epidermally expressed mNG::EFA-6 exhibited discrete foci at the distal neurite and a predominantly diffuse pattern along the proximal region, recapitulating the characteristic spatial distribution along ALM neurites (Fig. S5A, B). Also, consistent with the cell-autonomous requirement for neuronal EFA-6 in neurite growth regulation, epidermal EFA-6 expression did not rescue the ALM or PLM neurite overgrowth phenotypes associated with loss of *efa-6* (Fig. S5C-E).

We additionally employed a split-GFP reconstitution strategy to amplify and visualize the endogenous GFP::EFA-6 signals in neurons (He et al., 2019; Kamiyama et al., 2016). We tagged endogenous EFA-6 with seven tandem copies of GFP11, while the complementary GFP1-10 fragment was expressed specifically in either neurons or the epidermis. Reconstituted GFP::EFA-6 fluorescence in either tissue recapitulated the characteristic spatial distribution observed with endogenously GFP-tagged EFA-6 (Fig. S6A, B). This suggests that, while neuronal EFA-6 is expressed at very low levels, it overall mirrors epidermal EFA-6 spatial patterning.

Together, these findings suggest that GFP::EFA-6 associated with ALM neurites is primarily in surrounding epidermis, while neuronal and epidermal EFA-6 exhibit similar proximal-distal spatial distributions. Hereafter, we refer to the region of the epidermal cortex immediately adjacent to the ALM neurite as the neuron-epidermal interface, distinguishing it from the surrounding epidermal cortex.

### Specialized extracellular matrix regulates EFA-6 spatial patterning at neuron-epidermal interface

To identify regulators of EFA-6 spatial patterning we screened for aberrant localization of ALM-associated GFP::EFA-6 (Methods). Among the isolated mutants, we identified a class of mutations that disrupted the diffuse distribution of EFA-6 along the proximal ALM neurite, inducing ectopic foci without substantially altering the characteristic punctate distribution at the distal neurite tip. Combining whole-genome sequencing with genetic analysis, we identified one alleles of *mec-1* and three alleles of *him-4* (Fig. 2A).

**Figure 2:**
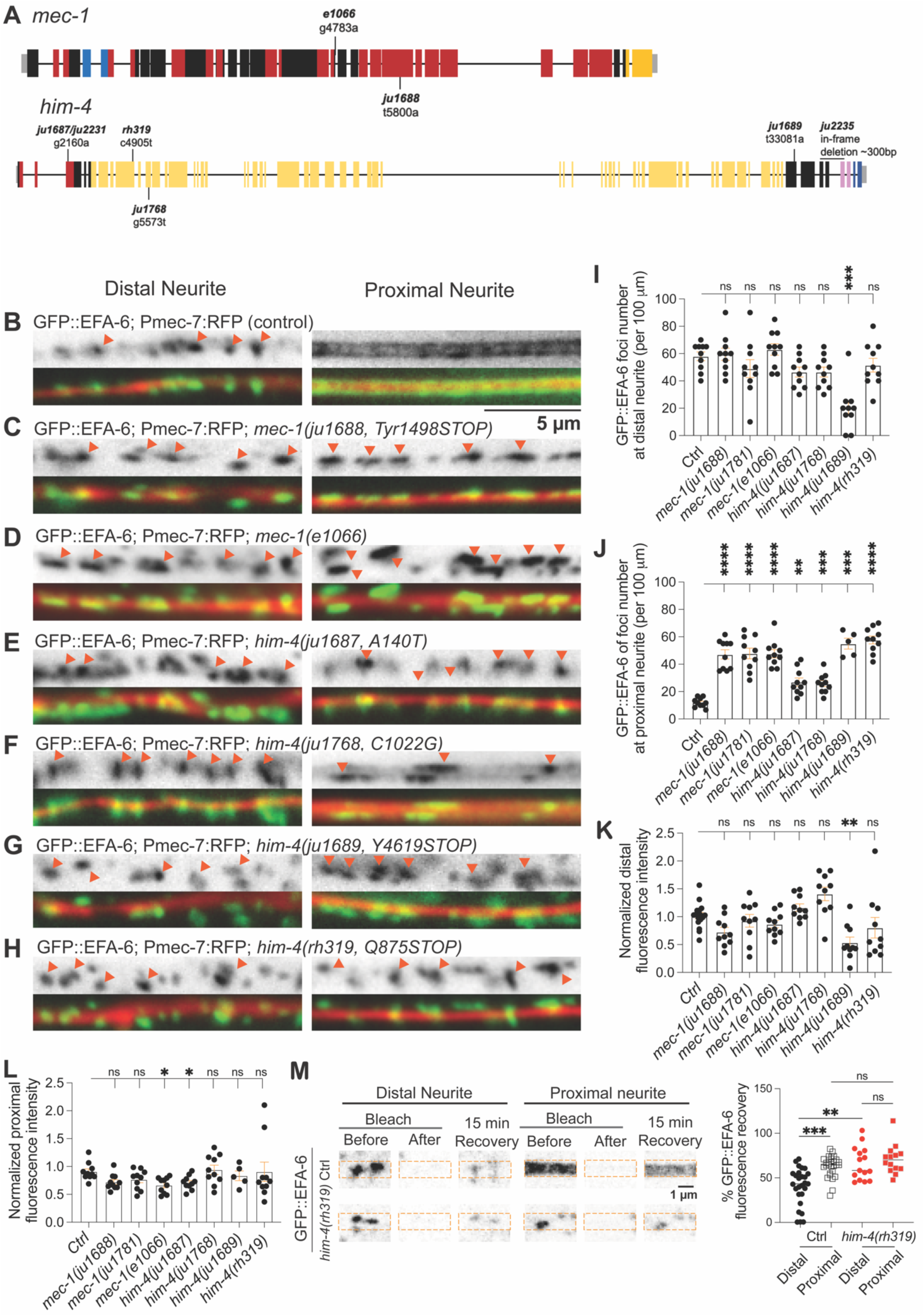
An unbiased genetic screen identifies ECM components as regulators of EFA-6 patterning. **(A)** Illustration showing *mec-1* and *him-4* gene organization with mutations isolated from the forward genetic screen and known loss-of-function mutations. Gray boxes represent untranslated regions (UTRs); colored boxes represent exons; solid black lines represent introns. Different colors indicate different domains. For *mec-1*: Red: Kunitz domain; blue: EGF domains; yellow: cysteine-rich C-terminal domain. For *him-4*: Red, VWA domain; yellow, Ig repeats; pink, EGF domains; blue, FC domain. **(B-H)** Representative compound fluorescence images of GFP::EFA-6 in (B) wildtype control (ctrl) and extracellular matrix components mutants: (C) *mec-1(ju1688, Tyr1498STOP),* (D) *mec-1(e1066),* (E) *him-4(ju1687, A140T),* (F) *him-4(ju1768, C1022G),* (G) *him-4(ju1689, Y4619STOP), and* (H) *him-4(rh319, Q875STOP)* mutant backgrounds. Images show distal and proximal ROIs of ALM neurites (same as Fig. 1A) in L4 stage animals. The upper panels show the GFP channel alone, whereas the lower panels show merged GFP and RFP channels. In control animals, GFP::EFA-6 forms discrete foci at the distal ROI and exhibits diffuse expression at the proximal ROI. In ECM mutants, EFA-6 forms discrete foci at the distal and proximal ROIs. Orange arrows mark EFA-6 foci. n=10∼33 per genotype. Scale: 5μm. **(I-J)** Quantification of GFP::EFA-6 foci density at the distal (I) and proximal (J) regions of ALM neurite in control and ECM component mutants shown in panel (B-H). **(K-L)** Quantification of GFP::EFA-6 fluorescence intensity at the distal (K) and proximal (L) regions of ALM neurite in control and ECM component mutants shown in panel (B-H). Raw values were normalized to control GFP::EFA-6 fluorescence intensity. **(M)** Quantification of fractional fluorescent recovery after photobleaching analysis of GFP::EFA-6 in the distal and proximal regions of the ALM anterior neurite in wildtype control and *him-4(rh319)* loss of function background. Representative airyscan images before and after fluorescence bleaching and 15 mins of recovery are shown on the left side. Scale bar, 1 µm. **(I-M)** n=10∼21 for (I-L) and n≥10 for (M) per genotype. Data are presented as mean ± SEM. Statistics: Brown-Forsythe and Welch ANOVA tests for (I-L) and Kruskal-Wallis test followed by Dunn’s multiple-comparisons test for (M); ns, not significant (P > 0.05); *P < 0.05; **P < 0.01; ***P < 0.001; ****P < 0.0001.

*mec-1* encodes a secreted protein containing two epidermal growth factor (EGF)-like domains, 15 Kunitz domains, and a cysteine-rich C-terminal region (Fig. 2A; Fig. S7A). MEC-1 is essential for mECM formation and organization (Du et al., 1996; Emtage et al., 2004). We found that *mec-1(ju1688,* Tyr1498STOP) has a premature stop codon at Tyr1498, which is predicted to truncate MEC-1 and remove seven Kunitz domains and its C-terminal region (Fig. 2A; Fig. S7A). *mec-1*(Tyr1498STOP) animals exhibited lethargic behavior and failed to respond to plate tapping, similar to *mec-1* loss-of-function mutants. In *mec-1*(Tyr1498STOP) mutants, GFP::EFA-6 formed discrete foci along the entire ALM neurite (Fig. 2B, C), resulting in a significant increase in proximal foci density in these mutants (Fig. 2J). Despite this redistribution, the overall GFP::EFA-6 fluorescence intensity in the proximal region remained comparable to controls (Fig. 2L). Neither foci density nor fluorescence intensity was significantly altered at the distal neurite tip (Fig. 2I, K). To further validate that loss of function in *mec-1* disrupts EFA-6 patterning, we examined an independently characterized *mec-1(e1066)* allele containing a splice site mutation (Du et al., 1996; Emtage et al., 2004). *mec-1(e1066)* mutants exhibited a similar redistribution of GFP::EFA-6, with increased proximal foci formation and no change in distal neurite foci expression (Fig. 2D, I, J). However, *mec-1(e1066)* also caused a reduction in GFP::EFA-6 fluorescence intensity in the proximal region, while distal fluorescence intensity remained unaffected (Fig. 2K, L). The consistent disruption of proximal EFA-6 patterning across *mec-1* alleles supports a requirement for MEC-1 in maintaining the spatial organization of EFA-6 along ALM neurites.

We identified three independent mutations in *him-4*, encoding hemicentin, a ∼600 kDa evolutionarily conserved extracellular matrix protein. HIM-4 contains an N-terminal signal peptide, a von Willebrand factor type A (VWA) domain, 48 tandem immunoglobulin-like (Ig) domains, a hydrophobic spacer, three EGF-like domains, and a C-terminal fibulin-like module (Fig. 2A; Fig. S7B; Dong et al., 2006). HIM-4 is implicated in mechanosensory ECM organization, neuronal attachment to the epidermis, and mechanical coupling between tissues (Gianakas et al., 2023; Morrissey et al., 2014; Vogel & Hedgecock, 2001). HIM-4 is dispensable for mechanotransduction and mECM secretion (Nekimken et al., 2020). Previously characterized *him-4* null mutants exhibit defective TRN ensheathment, gonadal extrusion through the vulva, and an increased incidence of males among self-progeny.

The three isolated alleles affect distinct HIM-4 domains: *him-4(ju1687, A140T)* introduces an Ala140Thr substitution in the conserved VWA domain, *him-4(ju1768, C1022G)* introduces a Cys1022Gly substitution in the seventh Ig domain, and *him-4(ju1689, Y4619STOP)* introduces a premature stop codon predicted to remove the final two Ig domains, EGF-like domains, and fibulin-like module (Fig. 2A; Fig. S7B, C). All three alleles showed ectopic foci of GFP::EFA-6 along the ALM proximal neurite (Fig. 2E-G). However, their effects differed in severity and spatial distribution. Both *him-4(A140T)* and *him-4(C1022G)* significantly increased proximal foci density without affecting distal foci density, whereas *him-4(Y4619STOP)* increased proximal foci density but reduced distal foci density (Fig. 2I, J). Ectopic foci in *him-4(A140T)* exhibited heterogeneous morphologies, including comet-like structures displaced from the neurite and foci aligned with the neurite axis (Fig. S8A). GFP::EFA-6 fluorescence intensity was reduced proximally in *him-4(A140T)* and distally in *him-4(Y4619STOP)*, but remained comparable to controls in both regions of *him-4(C1022G)* mutants (Fig. 2K, L). All three alleles caused ALM ensheathment defects, ranging from relatively mild in *him-4(C1022G)* to more severe in *him-4(Y4619STOP)*, and increased the incidence of males among self-progeny. However, unlike previously characterized *him-4* null mutants, none exhibited gonadal extrusion defects (Fig. S8A-D).

To determine whether complete loss of HIM-4 similarly affects EFA-6 patterning, we examined an independent *him-4(rh319, Q875STOP)* allele (Fig. 2A; Vogel & Hedgecock, 2001). We found *him-4(rh319)* disrupted the diffuse proximal GFP::EFA-6 distribution, producing ectopic foci with heterogeneous morphologies, including comet-like and elongated foci along the neurite axis (Fig. 2H; Fig. S8). Proximal foci density increased significantly, whereas distal foci density and GFP::EFA-6 fluorescence intensity in both regions remained unchanged (Fig. 2I-L). Thus, complete loss of HIM-4 disrupts EFA-6 spatial patterning, causing ectopic foci in the normally diffuse proximal region of ALM neurites.

To examine whether the ectopic EFA-6 foci observed in *him-4* mutants originate from the surrounding epidermis, we performed tissue-specific depletion of GFP::EFA-6(AID) in *him-4(rh319)* background. Consistent with our earlier observations that epidermal EFA-6 constitutes the predominant population associated with ALM neurites, both ubiquitous and epidermis-specific depletion abolished ectopic EFA-6 foci formation and fluorescence intensity in *him-4* mutants (Fig. S9A-E). In contrast, neuronal EFA-6 depletion had no significant effect on ectopic GFP::EFA-6 foci formation or fluorescence intensity (Fig. S9A-E). These findings suggest that the ectopic EFA-6 foci induced by loss of HIM-4 originate predominantly from epidermal cells.

We next asked whether loss of HIM-4 altered the extent of fluorescence recovery of EFA-6 along ALM neurites. We performed FRAP analysis of GFP::EFA-6 in the proximal and distal regions of ALM neurites in *him-4(rh319)* loss-of-function mutants (Fig. 2M). We compared fluorescence recovery after 15 minutes and found that in *him-4* mutants distal GFP::EFA-6 foci showed significantly greater recovery compared to control, reaching levels that were not significantly different from those of diffuse proximal EFA-6 in wild-type animals. In *him-4* mutants, ectopic GFP::EFA-6 foci in the proximal region of ALM exhibited fluorescence recovery comparable to that of diffuse proximal EFA-6 in wildtype animals.

Collectively, these results support a role for specialized ECM in regulating both the spatial distribution and restricting epithelial EFA-6 exchange at the neuron-epidermal interface.

### HIM-4 organization along ALM neurites is associated with EFA-6 spatial patterning

Because HIM-4 is an ECM protein implicated in neuronal attachment and tissue organization, we next asked whether these mutations alter the expression or distribution of HIM-4 itself along ALM neurites. To address this, we genome-edited corresponding mutations in endogenously mNG tagged HIM-4 (HIM-4::mNG) (Keeley et al., 2020) and compared their expression pattern with the control. This approach allowed us to examine how distinct molecular alterations in HIM-4 affect its extracellular organization and how these changes relate to EFA-6 patterning at the adjacent epidermal cortex.

In control animals, HIM-4::mNG formed a continuous track along the ALM neurite, with reduced fluorescence intensity in the distal neurite compared to the proximal neurite (Fig. 3A, E). HIM-4::mNG track was adjacent to mScarlet-tagged EFA-6 (mSc::EFA-6), which exhibited discrete distal foci and a predominantly diffuse proximal distribution, resembling GFP::EFA-6. Although neither A140T nor C1022G substitutions caused overall morphology defects, both disrupted EFA-6 spatial patterning, consistent with their corresponding untagged *him-4* alleles. Both substitutions also disrupted the continuous HIM-4::mNG track, but to different extents (Fig. 3B, C). HIM-4(A140T)::mNG formed discrete accumulations along the neurite, frequently juxtaposed with ectopic mSc::EFA-6 foci. In contrast, HIM-4(C1022G)::mNG exhibited a less pronounced disruption, with mSc::EFA-6 enriched in regions containing HIM-4(C1022G)::mNG fluorescence and absent where HIM-4 fluorescence was undetectable. Both substitutions further reduced distal HIM-4::mNG fluorescence intensity without significantly affecting proximal intensity (Fig. 3F, G).

**Figure 3:**
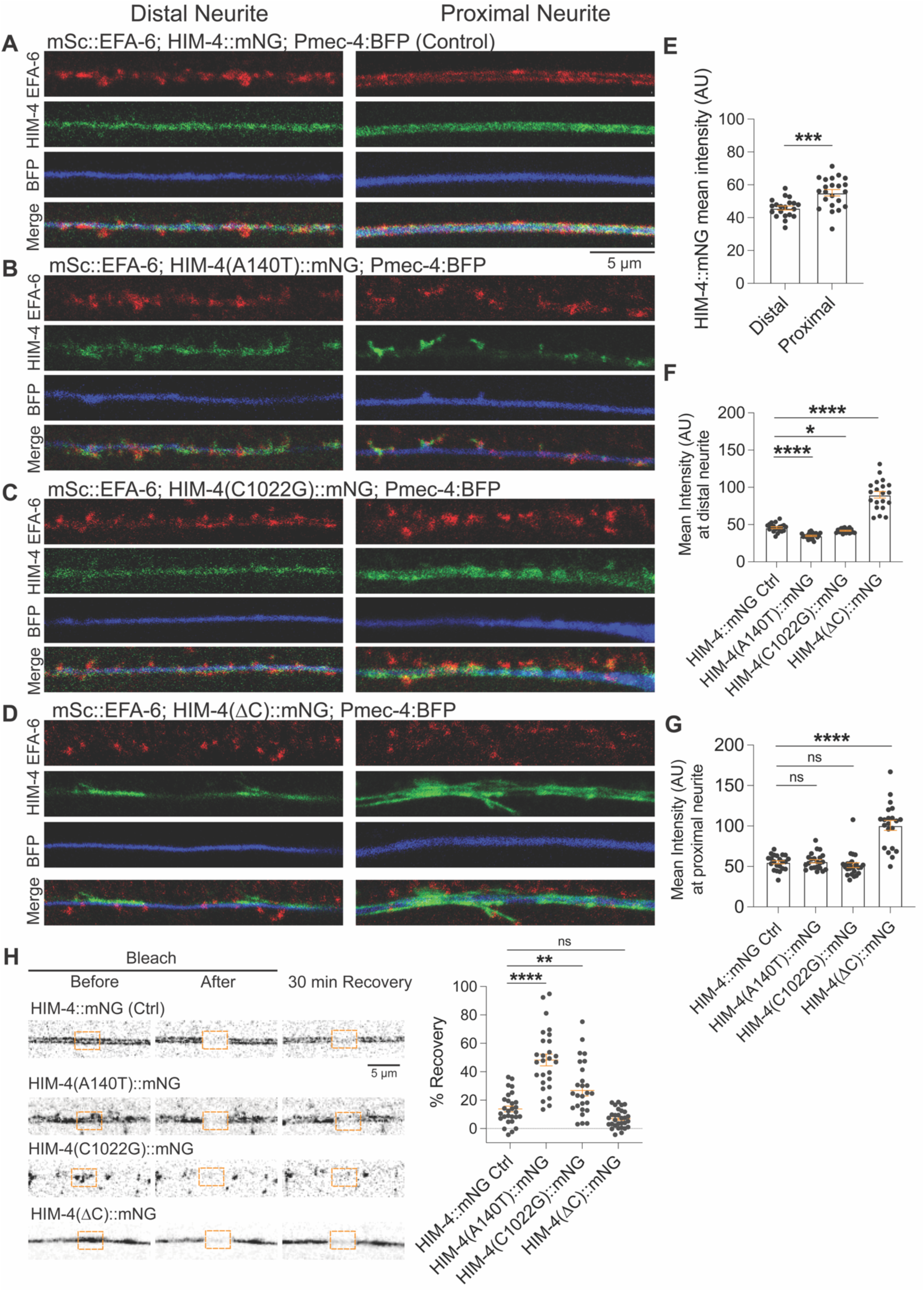
HIM-4 localization along ALM neurite controls EFA-6 spatial patterning. **(A-D)** Representative single-plane airyscan images of L4-stage mScarlet::EFA-6 co-expressed with mNG::HIM-4 in (A) wild-type HIM-4::mNG (control/ctrl), (B) HIM-4(A140T)::mNG (*ju2232*), (C) HIM-4(C1022G)::mNG (*ju2233*), or (D) HIM-4(ΔC)::mNG (*ju2235*), containing an in-frame C-terminal deletion. Images show distal and proximal ROIs of ALM neurites (same as Fig. 1A) in L4 stage animals. Pmec-7:BFP label ALM neurites. For each genotype, the panels show mScarlet channel (EFA-6), mNG channel (HIM-4), and BFP channel (ALM neurite) from top to bottom, followed by a merged image of all three channels. In control animals, HIM-4::mNG forms a continuous, relatively uniform track along the ALM neurite at both distal and proximal regions. HIM-4(A140T)::mNG localizes along the ALM neurite but exhibits a discontinuous pattern, with regions of HIM-4 enrichment interrupted by gaps lacking detectable fluorescence. HIM-4(C1022G)::mNG exhibits a similar but less pronounced disruption of the continuous HIM-4 pattern. HIM-4(ΔC)::mNG displays increased fluorescence along the ALM neurite and a highly irregular pattern, characterized by regions lacking detectable fluorescence interspersed with patches of strong HIM-4 enrichment and occasional short branch-like structures. n=10 per genotype. Scale bar: 5 µm. **(E)** Quantification of mean HIM-4::mNG fluorescence intensity at the distal and proximal ROIs of the ALM neurite in control animals (same as Fig. 1A). **(F, G)** Quantification of mean HIM-4::mNG fluorescence intensity at the (F) distal and (G) proximal ROIs of the ALM neurite (same as Fig. 1A) for HIM-4 control and variants shown in panel (A-D). **(H)** Quantification of fractional fluorescence recovery after photobleaching of HIM-4::mNG at the proximal regions of the ALM neurite (same as Fig. 1A) in animals expressing wild-type HIM-4::mNG or the indicated HIM-4 variants: A140T (*ju2232*), C1022G (*ju2233*), and ΔC (*ju2235*). Representative Airyscan images acquired before photobleaching, immediately after photobleaching, and after 30 min of recovery are shown on the left. Scale bar, 5 µm. **(E-H)** n=10-12 animals per genotype for (E-H); Data are presented as mean ± SEM. Statistics: Unpaired t-test in (E), Brown-Forsythe and Welch ANOVA tests for (F, G) and ordinary one-way ANOVA for (H); ns, not significant (P > 0.05); *P < 0.05; ***P < 0.001; ****P < 0.0001.

To investigate the contribution of the HIM-4 C-terminal region implicated by the *him-4(Y4619STOP)* mutation, we generated an in-frame deletion of approximately 300 bp between the last Ig-like domain and the EGF-like domains in HIM-4::mNG animals, (HIM-4(ΔC)::mNG; Fig. 2A). HIM-4(ΔC)::mNG mutants exhibited no overt morphological defects but mSc::EFA-6 formed ectopic foci along the ALM neurite, resembling the phenotype observed in *him-4*(Y4619STOP) mutants (Fig. 3D). HIM-4(ΔC)::mNG also exhibited a discontinuous distribution along of ALM neurites, similar to A140T and C1022G substitutions (Fig. 3D). However, HIM-4(ΔC)::mNG fluorescence intensity was significantly increased compared with control HIM-4::mNG animals and the two HIM-4 substitution mutants (Fig. 3D, F, G). Abnormal branch-like HIM-4 structures were also observed along the neurite. These findings indicate that the local distribution of HIM-4 is associated with EFA-6 patterning at the adjacent epidermal cortex.

To determine whether the altered spatial distributions of HIM-4 variants were accompanied by changes in their fluorescence recovery properties, we performed FRAP analysis in the proximal neurite (Fig. 3H). Both HIM-4(A140T)::mNG and HIM-4(C1022G)::mNG exhibited increased fluorescence recovery compared with control HIM-4::mNG, whereas HIM-4(ΔC)::mNG showed no significant difference in recovery at the 30-minute endpoint. Thus, although all three HIM-4 variants exhibited altered spatial distributions along the ALM neurites, only A140T and C1022G substitutions significantly increased fluorescence recovery. These findings indicate that changes in HIM-4 spatial distribution are not necessarily accompanied by changes in its fluorescence recovery.

As animals with these *him-4* mutations are morphologically normal, we examined how these mutations affect HIM-4 expression in other tissues. In control animals. HIM-4 forms extracellular structures in the head, between body wall muscles and the pharyngeal basement membrane (Fig. S10A; Table S7; Vogel & Hedgecock, 2001). These structures were undetectable in HIM-4(A140T)::mNG animals, which also exhibited reduced HIM-4 fluorescence intensity in the vulva and germline (Fig. S10B, E-G). In contrast, HIM-4(C1022G)::mNG distribution and fluorescence intensity in these regions were comparable to controls (Fig. S10C, E-G). HIM-4(ΔC)::mNG exhibited abnormal extracellular structures and increased fluorescence intensity in the head, whereas fluorescence intensity in the vulva and germline remained comparable to controls (Fig. S10D-G). Together, these observations indicate that the A140T substitution and C-terminus deletion affects HIM-4 distribution across multiple tissues, whereas the effects of C1022G are more restricted to the ALM-associated HIM-4 distribution.

Altogether, these findings demonstrate that distinct structural regions of HIM-4 contribute differentially to its extracellular organization. Mutations in the N-terminal VWA and Ig-like domains disrupt the continuity of ALM associated HIM-4 tracks and increase fluorescence recovery, whereas deletion of the C-terminal region causes abnormal HIM-4 accumulation and disorganization along ALM neurites. Despite these distinct effects on HIM-4 organization, all three variants result in ectopic EFA-6 foci along ALM neurites. Conversely, *efa-*6 mutants had normal HIM-4::mNG localization (Fig. S11A-J). These data support a model in which the extracellular organization of HIM-4 contributes to the spatial regulation of EFA-6 at the adjacent epidermal cortex.

### EFA-6 domains differentially regulate its spatial patterning and dynamics

EFA-6 is a multidomain protein containing several functionally distinct regions. We next determined how each domain of EFA-6 contributes to its enrichment and spatial patterning at neuron-epidermal interface. We generated a series of EFA-6 constructs lacking individual domains (Fig. S12) and expressed them as epidermis-specific, single-copy transgenes in the *efa-6(ju1883)* loss-of-function background. Full-length mNG::EFA-6 recapitulated the endogenous GFP::EFA-6 distribution, exhibiting discrete foci in the distal region and a predominantly diffuse pattern along the proximal ALM neurite. In contrast, free mNG was uniformly distributed throughout the epidermis, without detectable enrichment along ALM neurites (Fig. 4A). The pleckstrin homology (PH) domain of EFA-6 is implicated in its recruitment to the plasma membrane. Deletion of the PH domain, mNG::EFA-6(ΔPH), substantially reduced EFA-6 fluorescence intensity along ALM neurites but did not completely abolish its enrichment at the interface (Fig. 4A-C). mNG::EFA-6(ΔPH) exhibited a predominantly punctate distribution in the proximal region, resulting in increased foci density (Fig. 4A, D, E).

**Figure 4.**
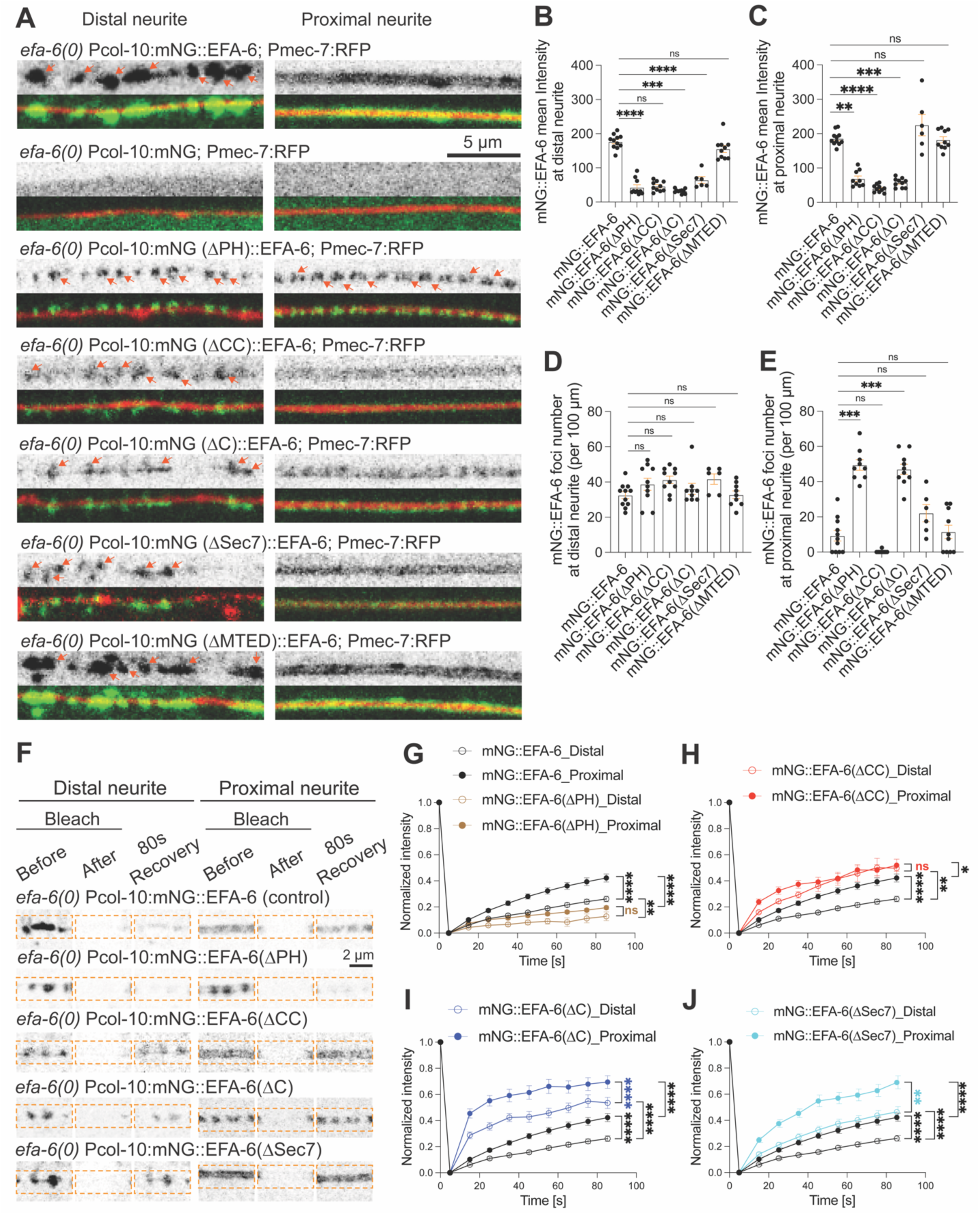
Distinct EFA-6 domains regulate spatial patterning and protein dynamics. **(A)** Representative single-plane confocal images of L4-stage animals expressing epithelial-specific (Pcol-10) full-length mNG::EFA-6, free mNG, or mNG::EFA-6 variants lacking the PH, coiled-coil (CC), C-terminal (ΔC), Sec7, or microtubule elimination domain (MTED) in *efa-6(ju1883)* null background as single-copy insertion. See Fig.S12 for construct details. Images show the distal and proximal ROIs of the ALM neurite (same as Fig. 1A), labelled by Pmec-7:RFP. The upper panels show the mNG channel alone, whereas the lower panels show merged mNG and RFP channels. Full-length mNG::EFA-6 is enriched along the ALM neurite and exhibits distinct spatial patterning, characterized by discrete foci in the distal region and a uniform distribution in the proximal region. In contrast, free mNG is diffusely distributed throughout the epidermis without specific enrichment along the ALM neurite. Deletion of the PH domain (ΔPH) reduces EFA-6 fluorescence intensity along the ALM neurite and disrupts its spatial patterning, resulting in a predominantly punctate distribution in both distal and proximal regions. Deletion of the CC domain (ΔCC) similarly reduces EFA-6 fluorescence intensity but largely preserves its spatial patterning. Deletion of the entire C-terminus (ΔC), encompassing both the CC and PH domains, also reduces EFA-6 fluorescence intensity and disrupts its spatial organization. The ΔC variant exhibits a more punctate distribution than ΔCC, but with less distinct foci than ΔPH, resulting in an intermediate pattern between the two individual domain deletions. Deletion of the Sec7 domain reduces EFA-6 fluorescence intensity along the ALM neurite without noticeably altering its spatial patterning. In contrast, deletion of the MTED has no obvious effect on EFA-6 fluorescence intensity or spatial organization. Orange arrows mark EFA-6 foci. n=10 animals per genotype. Scale bar, 5 µm. **(B-C)** Quantification of mean mNG::EFA-6 fluorescence intensity at the (B) distal and (C) proximal ROIs of ALM neurite (same as Fig. 1A) in the L4-stage animals expressing epithelial specific mNG::EFA-6 control and its variants shown in panel (A). **(D-E)** Quantification of mNG::EFA-6 foci density at the (D) distal and (E) proximal ROIs of ALM neurite (same as Fig. 1A) in the L4-stage animals expressing epithelial specific mNG::EFA-6 control and its variants shown in panel (A). **(F)** Representative airyscan images before photobleaching, after photobleaching, and 80 seconds of recovery in the distal and proximal ROIs of ALM neurite (same as Fig. 1A) of L4-stage animals expressing epithelial-specific full-length mNG::EFA-6 control or its variants lacking the PH, coiled-coil (CC), C-terminal (ΔC), or Sec7 in *efa-6* null background as single-copy insertion. Scale bar, 2 µm. **(G-J)** FRAP analysis of mNG::EFA-6 in the distal and proximal ROIs of ALM neurite (same as Fig. 1A) of L4-stage animals expressing epithelial-specific full-length mNG::EFA-6 control or mNG::EFA-6 variants shown in panel (F) in *efa-6* null background as single-copy insertion. **(B-E, G-J)** n = 10-12 animals per genotype. Data are presented as mean ± SEM. Statistics: Kruskal-Wallis test followed by Dunn’s multiple-comparisons test for (B-E) and RM two-way ANOVA with the Geisser Greenhouse correction for (G-J). ns, not significant (P > 0.05); *P < 0.05; **P < 0.01; ***P < 0.001; ****P < 0.0001.

Deletion of the coiled-coil (CC) domain similarly reduced EFA-6 fluorescence intensity along ALM neurites (Fig. 4A-C), but largely preserved its characteristic proximal-distal spatial patterning (Fig. 4A, D, E). Deletion of the entire C-terminal region containing both the PH and CC domains (mNG::EFA-6(ΔC)) produced an intermediate patterning phenotype such that mNG::EFA-6(ΔC) formed ectopic foci in the proximal region, but these foci were less pronounced than those observed in mNG::EFA-6(ΔPH) (Fig. 4A, D, E). Overall fluorescence intensity along ALM neurites remained reduced, comparable to that observed with the individual PH and CC domain deletions (Fig. 4A-C). Deletion of the Sec7 domain also reduced EFA-6 fluorescence intensity at the neuron-epidermal interface (Fig. 4A-C) without appreciably altering its characteristic proximal-distal distribution (Fig. 4A, D, E). In contrast, deletion of the microtubule elimination domain (MTED) had no significant effect on either EFA-6 fluorescence intensity or spatial patterning along ALM neurites (Fig. 4A-E). Together, these observations indicate that distinct EFA-6 domains contribute differently to its enrichment and spatial organization at the neuron-epidermal interface.

As described above, GFP::EFA-6 along ALM neurite displayed different mobility in the distal and proximal regions (Fig. 1E). We next examined how individual EFA-6 domains contribute to its fluorescence recovery properties at the neuron-epidermal interface. We found that deletion of the PH domain significantly reduced EFA-6 fluorescence recovery in both the distal and proximal regions compared to full-length mNG::EFA-6 control (Fig. 4F, G). In contrast, deletion of the CC domain increased fluorescence recovery in the distal and proximal region (Fig. 4F, H). Notably, despite their opposing effects on fluorescence recovery, deletion of either the PH or CC domain abolished the characteristic distal-proximal difference in EFA-6 recovery observed in control animals. Interestingly, deletion of the entire C-terminal region containing both the PH and CC domains increased EFA-6 fluorescence recovery in both distal and proximal regions (Fig. 4F, I). However, unlike the individual domain deletions, removal of the entire C-terminal region preserved the spatial difference in fluorescence recovery between these regions. Similarly, deletion of the Sec7 domain increased EFA-6 fluorescence recovery in both regions while maintaining the characteristic distal-proximal difference (Fig. 4F, J).

Together, these findings indicate that distinct EFA-6 domains differentially regulate its fluorescence recovery at the neuron-epidermal interface. In particular, the PH and CC domains contribute to the spatial differences in EFA-6 recovery between distal and proximal regions, despite exerting opposing effects on overall recovery. The preservation of these spatial differences following deletion of the entire C-terminal region further suggests that the contributions of the PH and CC domains are not simply additive and that additional regions of EFA-6 participate in regulating its spatially distinct fluorescence recovery properties.

### EFA-6 IDR is required and sufficient for its patterning at neuron-epidermal interface

Because deletion of individual structured domains did not completely abolish EFA-6 enrichment at the neuron-epidermal interface, we next investigated whether its intrinsically disordered region (IDR) contributes to interface targeting and spatial patterning. To address this, we generated an epidermis-specific, single-copy transgene expressing mNG::EFA-6 lacking the IDR (mNG::EFA-6(ΔIDR)) in the *efa-6(ju1883)* loss-of-function background (Methods; Table S1, 2). Deletion of the IDR dramatically reduced EFA-6 enrichment along the entire neuron-epidermal interface and abolished the characteristic foci at the distal neurite tip (Fig. 5A, F-I). To validate these findings at endogenous expression levels, we examined a GFP::EFA-6(ΔIDR) knock-in allele (Sandhu et al., 2024). Consistent with the epidermis-specific transgene, endogenous deletion of the IDR substantially reduced EFA-6 enrichment along ALM neurites and disrupted distal foci formation (Fig. S13A-C). Notably, IDR deletion selectively impaired EFA-6 enrichment at the neuron-epidermal interface without disrupting its localization at other epithelial cell-cell boundaries, including seam cell-epidermal junctions (Fig. S13D). Furthermore, in *him-4(rh319)*, deletion of the IDR markedly reduced EFA-6 fluorescence along ALM neurites and abolished the ectopic foci induced by loss of HIM-4 (Fig. 5B, F-I). These findings establish that the EFA-6 IDR is essential for its enrichment at the neuron-epidermal interface and for both the normal distal foci and ectopic foci associated with disrupted HIM-4 organization.

**Figure 5.**
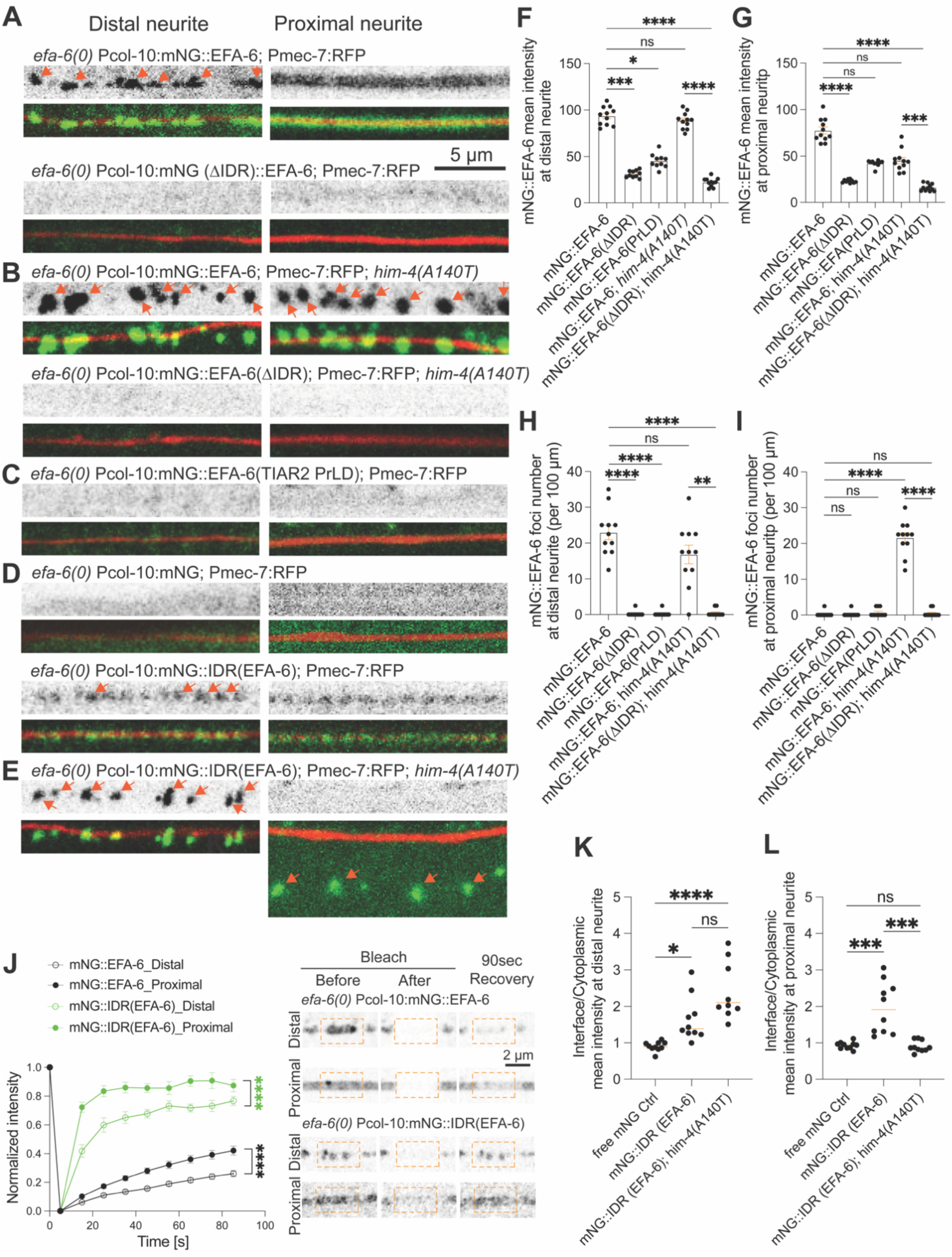
EFA-6 IDR is necessary and sufficient for enrichment at the neuron-epidermal interface. **(A-E)** Representative single-plane confocal images of L4-stage animals expressing epithelial-specific transgenes either with wild-type or *him-4(A140T)* mutation in *efa-6* null background as single-copy insertions. Constructs include full-length mNG::EFA-6, mNG::EFA-6(ΔIDR), mNG::EFA-6(PrLD) where EFA-6 IDR is replaced with the prion-like domain (PrLD) of TIAR-2, free mNG, and the isolated EFA-6 IDR (mNG::IDR). See Fig S12 for construct details. Images show the distal and proximal ROIs of the ALM neurite (same as Fig. 1A), labeled with Pmec-7:RFP. The upper panels show the mNG channel alone, whereas the lower panels show merged mNG and RFP channels. In animals with wild-type, full-length mNG::EFA-6 exhibits distinct spatial enrichment along the ALM neurite, whereas deletion of the IDR markedly reduces EFA-6 fluorescence intensity along the neurite and abolishes detectable foci formation (A). Similarly, replacement of the native IDR with the TIAR-2 PrLD abolishes EFA-6 enrichment along the neurite (C). mNG::EFA-6(ΔIDR) exhibit similar localization defects in *him-4(A140T)* mutant background (B). In contrast, the isolated mNG::IDR is sufficient for enrichment along both distal and proximal ALM regions in animals with wild-type, whereas free mNG remains diffusely distributed throughout the epidermis without specific neurite-associated enrichment (D). In *him-4(A140T)* mutants, mNG::IDR forms distinct foci along the distal ALM neurite, while its enrichment along the proximal region is reduced, accompanied by the formation of ectopic foci predominantly away from the neurite (E). Orange arrows mark EFA-6 foci. n=10 per genotype. Scale bar, 5 µm. **(F-G)** Quantification of mean mNG fluorescence intensity at the (F) distal and (G) proximal ROIs of the ALM neurite (same as Fig. 1A) in L4-stage animals expressing epithelial-specific full-length mNG::EFA-6 or EFA-6 variants lacking the native IDR, either through deletion (mNG::EFA-6(ΔIDR)) or replacement with the TIAR-2 PrLD. All transgenes were expressed in the *efa-6* null background as single-copy insertions, either with wild-type or *him-4(A140T)* mutation. **(H-I)** Quantification of mNG::EFA-6 foci density at the (H) distal and (I) proximal ROIs of the ALM neurite (same as Fig. 1A) in L4-stage animals for the same full-length, ΔIDR, and TIAR-2 PrLD replacement constructs and genetic backgrounds described in (B-C). **(J)** FRAP analysis comparing full-length mNG::EFA-6 and the isolated mNG::IDR epithelial-specific transgenes at the distal and proximal ROIs of the ALM neurite (same as Fig. 1A) in L4-stage animals. Representative Airyscan images acquired before photobleaching, immediately after photobleaching, and after 80 s of recovery are shown on the right. Scale bar, 2 µm. **(K-L)** Quantification of fluorescence enrichment at the (K) distal and (L) proximal ROIs of the ALM neurite (same as Fig. 1A) in L4-stage animals expressing epithelial-specific free mNG (negative control) or the isolated EFA-6 IDR tagged with mNG (mNG::IDR) in wild-type or *him-4(A140T)* mutation. All constructs were expressed in the *efa-6* null background as single-copy transgenes. **(F-L)** n=10-12 animals per genotype. Data are presented as mean ± SEM. Statistical significance was assessed using the Kruskal-Wallis test followed by Dunn’s multiple-comparisons test for (F-I, K, L), and two-way repeated-measures ANOVA with Geisser-Greenhouse correction for (J). ns, not significant (P > 0.05); *P < 0.05; **P < 0.01; ***P < 0.001; ****P < 0.0001.

We next asked whether this requirement reflects a specific property of the EFA-6 IDR or whether it could be fulfilled by a heterologous, phase-separation associated disordered region. To test this, we replaced the native EFA-6 IDR with the prion-like low-complexity domain (PrLD) of TIAR-2, previously shown to undergo phase separation in neurons and reported to be expressed in epidermal cells (Andrusiak et al., 2019; Rousakis et al., 2014). The resulting chimeric protein, mNG::EFA-6(PrLD) was not enriched at neuron-epidermal interfaces (Fig. 5C, F-I), although it formed ectopic foci within the surrounding epidermis and seam cells (Fig. S13E). Thus, although the heterologous PrLD formed ectopic foci, it did not recapitulate EFA-6 IDR enrichment or patterning along ALM neurites. These observations suggest that interface targeting of EFA-6 depends on sequence-specific properties of the native IDR rather than simply the presence of a disordered region associated with phase separation.

Having established that EFA-6 IDR is necessary for its enrichment, we next asked whether it is sufficient to direct protein enrichment at the neuron-epidermal interface independently of the remaining EFA-6 domains. To address this, we expressed mNG::IDR as an epidermis-specific, single-copy transgene. Compared with the free mNG control, mNG::IDR exhibited significant enrichment along the neuron-epidermal interface (Fig. 5D, K, L) but did not accumulate at seam cell-epidermal boundaries (Fig. S13D). This indicates that the IDR is sufficient to selectively target EFA-6 to neuron-epidermal interface.

We next examined whether HIM-4 regulates the spatial organization of mNG::IDR. In *him-4* loss-of-function mutants, mNG::IDR enrichment at the distal neurite remained largely unchanged, whereas its accumulation along the proximal region was significantly reduced (Fig. 5E, K, L). Notably, mNG:: IDR formed discrete ectopic comet-like foci that were predominantly displaced away from the ALM neurite into the surrounding epidermal cortex, similar to but more penetrant than the GFP::EFA-6 pattern disruption in *him-4* mutants (Fig. 5E; Fig. S14A-F). These observations suggest that the EFA-6 IDR is sufficient to support focal accumulation, whereas HIM-4 organization contributes to the spatial restriction of IDR enrichment to the neuron-epidermal interface.

Finally, we examined the fluorescence recovery properties of mNG::IDR. Compared with full-length mNG::EFA-6 control, mNG::IDR exhibited substantially greater fluorescence recovery (Fig. 5J). Despite this increased recovery rate, mNG::IDR retained the characteristic difference in fluorescence recovery between the distal and proximal regions of ALM neurites. These findings suggest that EFA-6 IDR contains determinants sufficient to establish spatially distinct fluorescence recovery properties, whereas additional EFA-6 domains contribute to restricting molecular exchange at the interface.

Together, these findings suggest that EFA-6 enrichment at neuron-epidermal interfaces depends on two distinct but complementary mechanisms: sequence-dependent targeting mediated by its IDR and spatial regulation by the extracellular protein HIM-4. The ability of the isolated IDR to retain regional differences in fluorescence recovery further indicates that determinants of spatially distinct EFA-6 organization reside within the IDR, while other EFA-6 domains modulate its molecular exchange at the interface.

### Identification of Epidermal-Neuronal Interface Recognition Motif within EFA-6 IDR

We next sought to identify IDR regions involved in selective targeting and enrichment at the neuron-epidermal interface. To address this, we systematically truncated the 284 amino-acid EFA-6 IDR and examined the ability of individual fragments to enrich at the interface (Fig. S15). We over-expressed mNG fused EFA-6 IDR and shorter fragments of IDR using a promoter specific to the adult epidermis (*col-19*; see Methods). Consistent with our single-copy expression analysis, the full-length IDR exhibited pronounced enrichment at the neuron-epidermal interface (Fig. S16A, N). Notably, despite the elevated expression levels, the EFA-6 IDR did not form detectable condensates in the surrounding epidermal cytoplasm, indicating that increased IDR abundance alone was insufficient to induce cytoplasmic condensation. In contrast, free mNG expressed under the same promoter remained diffusely distributed throughout the epidermis without detectable enrichment at the interface (Fig. S16B, N). By systematically analyzing mNG-tagged IDR fragments, we narrowed the interface-targeting region to a 14 amino-acid sequence (A123-E136) that retained significant enrichment at the neuron-epidermal interface compared with the free mNG control (Fig. S16C-I, N). By subdividing this 14 amino-acid fragment into overlapping peptides of 6-8 amino acids, with 2-3 amino acids of overlap between adjacent fragments (Fig. S15), we identified an eight amino-acid sequence, AIRMLKTA (A123-A130), that when overexpressed in epidermis of *efa-6* null animals, was sufficient to confer enrichment at the neuron-epidermal interface and to recapitulate the characteristic spatial patterning of full-length EFA-6 IDR (Fig. S16I-N). We therefore designated this sequence the Epidermal-Neuronal Interface Recognition Motif (ENIRM). ENIRM peptide also phenocopied the full-length IDR expression pattern. To determine whether ENIRM-mediated interface targeting depends on its specific amino acid sequence, we generated two scrambled variants of the 14 amino acid fragment containing ENIRM. Neither scrambled variant exhibited detectable enrichment at the neuron-epidermal interface, in contrast to the full-length IDR and the 14- and 8-amino-acid targeting fragments (Fig. S17A-F, H). To further establish whether ENIRM is required for interface recognition in the context of the full-length IDR, we expressed mNG::IDR lacking the 14 amino acid region containing ENIRM (A123-E136) and observed a complete abolishment of IDR enrichment at the neuron-epidermal interface (Fig. S17G, H). Together, these findings identify an 8 amino acid motif that selectively targets EFA-6 IDR to the neuron-epidermal interface.

### ENIRM and surrounding IDR sequences cooperatively achieve neuron-epidermal interface targeting at near physiological levels

Next, we asked whether ENIRM could mediate interface targeting at near-physiological expression levels. To address this, we generated epidermis-specific, single-copy transgenic lines expressing the corresponding constructs approaching physiological expression levels (Fig. S12; Fig. S17I, J). We observed that expression of IDR lacking the 14 amino-acid containing ENIRM (A123-E136) did not show enrichment at the neuron-epidermal interface (Fig. 6A-C, G, H), further establishing that the ENIRM-containing region is essential for IDR-mediated interface targeting. Conversely, we asked whether ENIRM alone was sufficient to confer interface enrichment at lower expression levels by expressing mNG tagged 14-amino-acid fragment containing ENIRM (A123-E136) as an epidermis-specific, single-copy transgene. We found that mNG-tagged fragment did not exhibit detectable enrichment at the neuron-epidermal interface (Fig. 6D, G, H). These findings indicate that, although ENIRM is sufficient for interface targeting under overexpression conditions, ENIRM alone is insufficient to support detectable enrichment at near-physiological expression levels. We therefore investigated whether the intrinsically disordered sequences flanking ENIRM contribute to its enrichment at the neuron-epidermal interface. To address this, we expressed constructs containing the ENIRM-bearing fragment together with either its N-terminal (G59-E136) or C-terminal (A123-L342) flanking sequence as epidermis-specific, single-copy transgenes. Interestingly, addition of either flanking region resulted in significant detectable enrichment at the neuron-epidermal interface (Fig. 6E-H). These observations suggest that ENIRM provides a sequence-specific determinant for interface recognition, whereas surrounding disordered regions enhance its ability to accumulate at the interface under near-physiological expression conditions.

**Figure 6.**
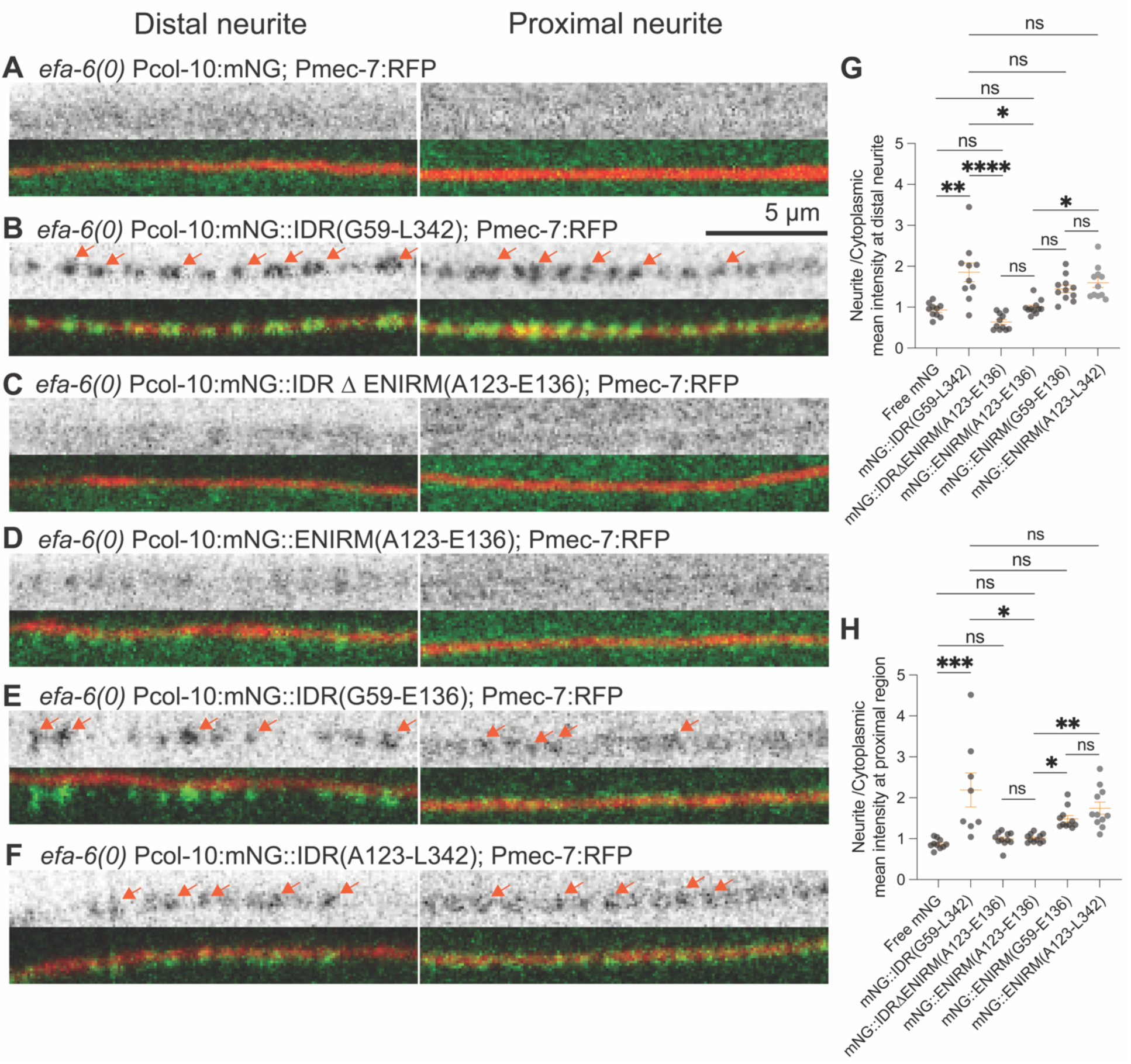
ENIRM and surrounding intrinsically disordered sequences cooperate to mediate neuron-epidermal interface targeting at near-physiological expression levels. **(A-F)** Representative single-plane confocal images of L4-stage animals expressing epithelial-specific transgenes in the *efa-6* null background. Constructs include (A) free mNG, (B) the full-length EFA-6 IDR tagged with mNG (mNG::IDR), (C) the full-length IDR lacking the 14-amino-acid ENIRM-containing sequence (mNG::IDRΔENIRM(A123-E136)), (D) the isolated 14-amino-acid ENIRM-containing fragment (mNG::ENIRM(A123-E136)), (E) the 14-amino-acid fragment combined with its N-terminal flanking disordered sequence (mNG::IDR(G59-E136)), and (F) the 14-amino-acid fragment combined with its C-terminal flanking disordered sequence (mNG::IDR(A123-L342). See Fig. S15 for transgene details. Images show the distal and proximal ROIs of the ALM neurite (same as Fig. 1A), labeled with Pmec-7:RFP. The upper panels show the mNG channel alone, whereas the lower panels show merged mNG and RFP channels. The full-length IDR is enriched along the ALM neurite, whereas deletion of the 14-amino-acid ENIRM-containing sequence abolishes this enrichment. Unlike under overexpression conditions, the isolated 14-amino-acid fragment is insufficient for detectable neuron-epidermal interface targeting at near-physiological expression levels. However, inclusion of either the N-terminal or C-terminal flanking disordered sequence enhances recruitment of the ENIRM-containing fragment to the ALM neurite, indicating that sequences surrounding ENIRM contribute to efficient interface targeting. Orange arrows mark EFA-6 foci. n=10 animals per genotype. Scale bar, 5 µm. **(G, H)** Quantification of fluorescence enrichment in the (G) distal and (H) proximal ROIs of the ALM neurite (same as Fig. 1A) in L4-stage animals expressing the epithelial-specific constructs shown in (A-F). n=10-15 animals per genotype. Data are presented as mean ± SEM. Statistical significance was assessed using the Kruskal-Wallis test followed by Dunn’s multiple-comparisons test. ns, not significant (P > 0.05); *P < 0.05; **P < 0.01; ***P < 0.001; ****P < 0.0001.

To further investigate how flanking disordered sequences influence ENIRM-dependent interface enrichment, we examined extrachromosomal arrays. In such overexpression conditions, either N (G59-Thr122)- and C (S137-L342)-terminus ENIRM flanking sequences did not show any significant enrichment along the neurites whereas their enrichment was significantly enhanced when the14 amino acid region containing ENIRM (A123-E136) was included (Fig. S18A-H). Next, we examined the relationship between absolute fluorescence intensity at the neuron-epidermal interface and cytoplasmic fluorescence intensity across animals expressing different levels of each construct. Addition of the N-terminal flanking sequence to ENIRM (G59-E136) shifted this relationship toward lower cytoplasmic fluorescence intensities, suggesting enhanced interface accumulation relative to cytoplasmic abundance. In contrast, addition of the C-terminal flanking sequence to the ENIRM (A123-L342) shifted the relationship toward higher cytoplasmic fluorescence intensities, indicating reduced interface accumulation at comparable cytoplasmic expression levels (Fig. S18I). Consistent with these observations, addition of the N-terminal sequence modestly increased mean interface fluorescence intensity, whereas the C-terminal sequence significantly reduced it under overexpression conditions (Fig. S18H). Together, these findings identify ENIRM as an essential for EFA-6 targeting to the neuron-epidermal interface, while revealing that its flanking disordered sequences enhance enrichment at near-physiological expression levels and differentially regulate accumulation in a concentration-dependent manner.

### Disrupted EFA-6 patterning contributes to age-dependent neurite defects in *him-4* mutants

We next asked whether proper spatial organization of EFA-6 is required to maintain ALM neuronal morphology. Previous studies have reported age-dependent ectopic neurite branching in *him-4* null mutants (Pan et al., 2011). *him-*4 mutants overall displayed normal ALM development up to L4 stage. However, in 5-day old adults, *him-4* mutant ALMs exhibited prominent morphological defects, including ectopic neurite branching and localized neurite swellings (Fig. 7A-C). Neither phenotype was observed in control animals. *efa-6* null mutants did not exhibit these defects, although they displayed the previously reported neurite overshooting phenotype associated with the cell-autonomous function of neuronal EFA-6. In *him-4(A140T)* mutants, EFA-6 spatial patterning at the neuron-epidermal interface is disrupted (Fig 2). To determine whether altered EFA-6 contributes to the neurite defects observed in *him-4(A140T)* mutants, we examined ALM morphology in *efa-6; him-4* double mutants. Loss of *efa-6* significantly reduced ectopic neurite branching but did not appreciably affect the neurite swelling phenotype. Expression of full-length EFA-6 specifically in the epidermis modestly increased ectopic branching compared with *efa-6; him-4* double mutants, although branching remained significantly lower than in *him-4* single mutants. However, it had no effect on neurite swelling. In contrast, epidermal expression of EFA-6 lacking its intrinsically disordered region (EFA-6(ΔIDR)) increase ectopic branching similar to *him-4* single mutants. Interestingly, epidermal expression of EFA-6(ΔIDR) markedly reduced neurite swellings compared with *efa-6; him-4* double mutants. Together, these findings suggest that disrupted epidermal EFA-6 organization contributes to age-dependent ALM neurite defects in *him-4* mutants. Furthermore, the distinct effects of full-length EFA-6 and EFA-6(ΔIDR) suggest that the IDR influences how epidermal EFA-6 affects neurite morphology in the context of disrupted extracellular matrix organization.

**Figure 7.**
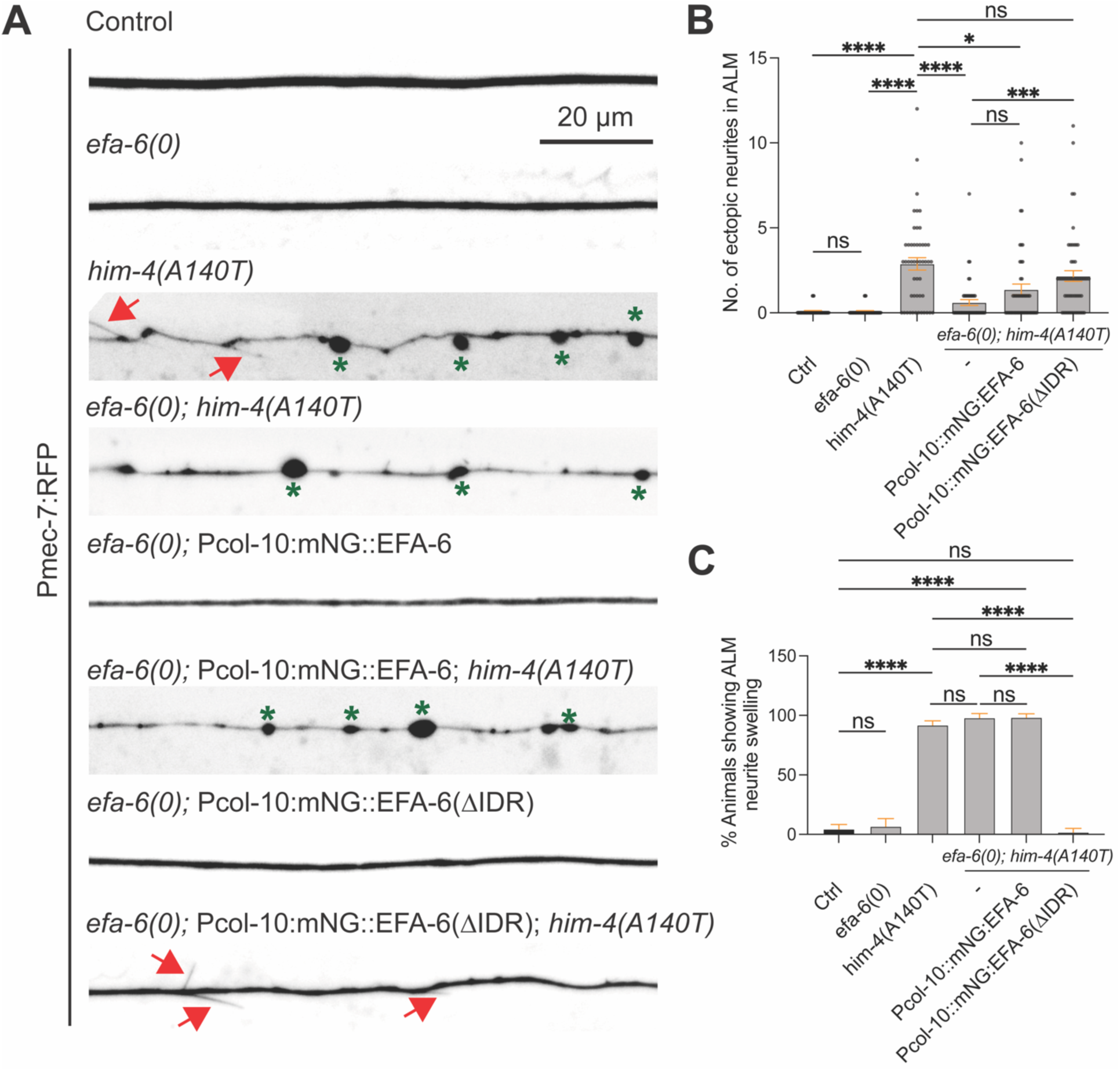
Disrupted EFA-6 patterning contributes to neurite defects in ECM mutants. **(A)** Representative confocal images of ALM neurites labeled with *Pmec-7*:RFP in wildtype, *efa-6(0)*, *him-4(A140T)*, and *efa-6(0); him-4(A140T)* animals, as well as *efa-6(0)* and *efa-6(0); him-4(A140T)* animals expressing epidermis-specific mNG::EFA-6 or mNG::EFA-6(ΔIDR) transgenes, as indicated. Images were acquired from animals five days after the L4 stage (D5). Red arrows mark ectopic branches and green asterisk indicate neurite swellings. *him-4(A140T)* mutants exhibit ectopic neurite branching and localized neurite swellings, whereas control and *efa-6(0)* animals show largely normal neurite morphology. Scale bar, 20 μm. **(B)** Quantification of the number of ectopic branches along ALM neurites in the indicated genotypes. Loss of *efa-6* significantly reduces ectopic branching in *him-4(A140T)* mutants. Epidermal expression of full-length mNG::EFA-6 partially restores ectopic branching, whereas expression of mNG::EFA-6(ΔIDR) does not significantly increase branching compared with *efa-6(0); him-4(A140T)* double mutants. Each dot represents an individual ALM neurite. **(C)** Quantification of the percentage of animals exhibiting ALM neurite swelling in the indicated genotypes. Loss of *efa-6* does not significantly suppress neurite swelling in *him-4(A140T)* mutants. Similarly, epidermal expression of full-length mNG::EFA-6 does not significantly alter the frequency of swelling. In contrast, epidermal expression of mNG::EFA-6(ΔIDR) markedly reduces the percentage of animals exhibiting neurite swelling. **(B, C)** n=70-90 animals per genotype; Data are presented as mean ± SEM. Statistics: Brown-Forsythe and Welch ANOVA tests for (B, C); ns, not significant (P > 0.05); *P < 0.05; ***P < 0.0001; ****P < 0.0001.

## DISCUSSION

In this study, we demonstrate that EFA-6 is selectively enriched at the epidermal cortex adjacent to ALM mechanosensory neurites and that the spatial distribution of EFA-6 undergo distinct changes during development. We identify a short motif within the IDR of EFA-6 (ENIRM) that recognizes this specialized epidermal cortex and promotes EFA-6 enrichment at neuron-epidermal interfaces. Other structured domains of EFA-6 also differentially regulate its enrichment, patterning, and molecular exchange at the interface. Developmental re-distribution of EFA-6 from foci to a diffused state at the interface requires proper localization of the conserved extracellular matrix protein HIM-4 along the neurite. We show that in *him-4* mutants ALM neurites exhibit age-dependent neurite defects that are suppressed by loss of function in *efa-6*. Together, these findings reveal a mechanism by which extracellular matrix organization influences the spatial distribution and dynamics of a cortical microtubule regulator at specialized neuron-epidermal interfaces.

Previous studies have shown that epithelial cells organize their spectrin cytoskeleton into nanoscale structures along neuronal processes (MPS: membrane-associated periodic scaffold), with neuron-dependent assembly of these structures contributing to neuronal protection from mechanical damage (Coakley et al., 2026). Our findings suggest that the epidermal cortex may also organize larger macro-scale, spatially restricted domains defined by EFA-6 condensates at neuron-epidermal interfaces. However, unlike spectrin nano-domains, these macro-domains appear to undergo developmental remodeling that correlates with neurite ensheathment within the epidermis. Such macro-domains could potentially accommodate the changing requirements of neurite growth, epithelial wrapping, and mechanical stabilization as the interface develops. The persistence of EFA-6 foci in the distal region of ALM and diffused distribution along the proximal neurite suggests that these epidermal macro-domains are organized into spatially distinct domains along a single neuronal process. These distal and proximal domains may reflect different structural or functional requirements at these interfaces.

Presence of EFA-6 foci before neurite ensheathment suggests that the formation of these cortical domains is likely regulated by intrinsic developmental programs or early ensheathment cues. Whereas their association with neurite and re-distribution in adults require proper ECM formation and neuron-epidermal attachment. Supporting this idea, in *mec-1* and *him-4* mutants where ensheathment is impaired, EFA-6 forms ectopic foci along the neurite. In *him-4* mutants, some of these foci remain closely associated with ALM neurites, whereas others are displaced into the surrounding epidermal cortex. This suggests that ECM is not strictly required for EFA-6 foci formation but instead contribute to EFA-6 foci spatial positioning and subsequent redistribution to diffuse pattern along the neurite. This raises the question of what initially drives EFA-6 accumulation into discrete foci and how ECM organization influences their localization and maturation. Our FRAP revealed that EFA-6 foci at both distal and proximal neurite of ALM in *him-4* mutants exhibit faster recovery than EFA-6 foci at the distal region of ALM in control animals, suggesting that the formation of morphologically similar foci does not necessarily reflect equivalent molecular properties. These differences may arise from changes in EFA-6 interactions with cortical components that regulate its retention or exchange at the neuron-epidermal interface. Thus, ECM organization may influence not only the spatial distribution of EFA-6 but also its molecular interactions and dynamic properties within the epidermal cortex.

An additional question is how EFA-6 achieves selective enrichment at the neuron-epidermal interface without undergoing widespread condensation in the surrounding cytoplasm. Although the EFA-6 IDR can undergo phase separation *in vitro* (Sandhu et al., 2024), we did not observe condensates even when EFA-6 IDR was expressed at much higher levels than endogenously tagged EFA-6. This suggests that increasing IDR concentration alone is insufficient to drive condensation in the epidermal cytoplasm. One possibility is that the intracellular environment restricts nonspecific self-association of IDR, whereas interactions with membrane-associated components locally favor EFA-6 foci formation at specialized cortical regions. Such spatial regulation could allow the same disordered sequences to remain predominantly diffuse in the cytoplasm while promoting selective enrichment of EFA-6 IDR at the neuron-epidermal interface.

Our experiments further suggest that EFA-6 selective enrichment at neuron-epidermal interfaces is regulated by distinct regions within the EFA-6 IDR. We identified a short peptide sequence, ENIRM, that is sufficient to confer interface recognition and enrichment when overexpressed. However, when expressed at the level comparable to endogenous EFA-6, ENIRM requires the flanking disordered sequences for enrichment at neuron-epidermal interfaces. These flanking regions differ in their sequence composition and influence interface enrichment differently depending on expression levels. The N-terminal flanking region is enriched in positively charged residues (net charge: +5), whereas the C-terminal region is predominantly enriched in negatively charged residues (net charge: -20) (Fig. S15B). Addition of the N-terminal region enhanced ENIRM-dependent interface enrichment, whereas addition of the C-terminal region reduced enrichment under overexpression conditions. Nevertheless, addition of either flanking region to the ENIRM supported interface enrichment when expressed near physiological levels, suggesting that their effects on ENIRM-mediated enrichment depends on protein abundance. These findings suggest that ENIRM provides sequence-specific recognition of the neuron-epidermal interface, while its flanking disordered sequences modulate the efficiency of enrichment across different expression levels. At lower concentrations, these flanking regions may stabilize otherwise weak interactions with interface-associated components, allowing EFA-6 to selectively enrich when protein abundance is limited. At higher concentrations, additional intra- or intermolecular interactions involving these sequences could alter ENIRM accessibility or compete with interface recruitment, thereby limiting further enrichment. Such expression-dependent modulation could potentially buffer interface-associated EFA-6 accumulation against changes in cellular protein abundance, thereby maintaining an optimal abundance at neuron-epidermal interfaces. The molecular interactions underlying these effects remain to be determined, as does whether changes in endogenous EFA-6 abundance contribute to its spatial patterning during development.

The plasma membrane cortex is organized into spatially distinct domains through interactions among membrane lipids, cytoskeletal networks, and membrane-associated proteins (Honigmann & Pralle, 2016; Mangiarotti & Dimova, 2026). Local differences in phosphoinositide composition can regulate the recruitment of peripheral membrane proteins, while cortical actin and anchored transmembrane proteins can restrict their lateral mobility and contribute to membrane compartmentalization (Fujiwara et al., 2002; Hammond et al., 2012; Bogaart et al., 2011). Multivalent protein interactions and membrane-associated condensates provide additional mechanisms for concentrating proteins within specialized cortical regions (Banjade & Rosen, 2014; Su et al., 2016). These mechanisms offer potential explanations for the distinct distributions and fluorescence recovery properties of EFA-6 along ALM neurites. In mammalian EFA6 proteins, the pleckstrin homology (PH) domain interacts with phosphatidylinositol 4,5-bisphosphate [PI(4,5)P₂], contributing to their association with the plasma membrane (Kanamarlapudi, 2014; Macia et al., 2008). The coiled-coil (CC) domain has also been implicated in cortical stabilization through interactions with the actin cytoskeleton; together the PH- and CC-containing regions have been associated with ECM-linked signaling (Derrien et al., 2002; Kanamarlapudi, 2014; Macia et al., 2019). Furthermore, in mammalian cells, the transmembrane aminopeptidase CD13 recruits EFA6 to cortical signaling complexes containing IQGAP1, ARF6, and β-integrins, promoting integrin recycling and focal adhesion turnover (Ghosh et al., 2019). These studies demonstrate that EFA6 localization can be influenced by membrane lipids, cytoskeletal interactions, and transmembrane protein complexes. Consistent with distinct contributions of these interactions to cortical organization, we found that deletion of either the PH or CC domain abolished the normal differences in EFA-6 fluorescence recovery between the distal and proximal ALM regions. However, deletion of the PH domain reduced fluorescence recovery, whereas deletion of the CC domain increased recovery, suggesting that these domains influence EFA-6 molecular exchange through different mechanisms. The Sec7 domain may provide an additional connection between EFA-6 organization and membrane geometry. In mammalian EFA6 proteins, the Sec7 domain interacts with endophilin, whose association with membrane is sensitive to curvature (Boulakirba et al., 2014). Because ALM neurites become progressively ensheathed within the epidermis during development, changes in local membrane geometry could potentially influence the recruitment or stabilization of EFA-6 or associated complexes. The reduced EFA-6 enrichment observed following Sec7 deletion is consistent with a contribution of this domain to cortical localization, although whether membrane curvature plays a role remains unknown.

One possible explanation for our findings is that the IDR mediates selective recruitment of EFA-6 to components associated with the neuron-epidermal interface, whereas interactions involving the PH, CC, and Sec7 domains influence its association with locally distinct cortical environments. Differences in membrane lipid composition, cytoskeletal scaffolding, or membrane geometry between proximal and distal regions could therefore contribute to the observed differences in EFA-6 distribution and fluorescence recovery. This model predicts that perturbing these cortical properties would alter EFA-6 organization even when IDR-mediated interface targeting remains intact.

What is the functional significance of these specialized cortical domains? The enrichment and spatial patterning of EFA-6 along ALM neurites raise the possibility that these domains regulate local microtubule organization at the epidermal cortex. The developmental transition from punctate to predominantly diffuse EFA-6 distribution along the proximal neurite may reflect changes in cortical microtubule organization as the neurite becomes progressively ensheathed within the epidermis. More broadly, our findings raise the possibility that ECM-derived cues spatially organize, and remodel specialized cortical domains, potentially allowing epithelial cells to adapt their cytoskeletal architecture to the changing developmental and functional requirements of neighboring neurons. How these EFA-6 enriched domains contribute to local microtubule regulation remains to be determined, MEC-5, a collagen required for the organization of mechanosensory complexes along touch receptor neurites, exhibits spatial patterning similar to EFA-6 along the ALM neurite.

MEC-5 forms discrete foci in the distal region but displays a more uniform distribution along the proximal region (Emtage et al., 2004). In *him-4* mutants, MEC-5 spatial patterning is completely disrupted where MEC-5 forms ectopic foci in the proximal neurite. Some of these foci lie close to the neurite whereas other the displaced far from the neurite. The striking similarity between MEC-5 and EFA-6 patterning raises the possibility that the spatial organization of the epidermal cortex is coordinated with the patterning of underlying ECM. This could help epithelial cells remodel their membrane and cortical cytoskeleton to accommodate both structural and functional changes associated with neurite ensheathment. However, whether MEC-5 patterning in the ECM correlates to EFA-6 patterning at epithelial cortex during ensheathment and how the two patterns are structurally and functionally linked remains to be examined.

Interestingly, LET-805/Myotactin, a transmembrane protein involved in epidermal attachment to the ECM, also exhibits a similar developmental and spatial distribution along PLM neurites (Coakley et al., 2020; Hresko et al., 1999). LET-805 initially forms foci that become more uniformly distributed along PLM proximal neurite in adults, while remaining punctate in the distal region. Although its distribution along ALM neurites remains to be examined, the similarity in its spatial organization and its established role in epidermal-ECM attachment raise the possibility that EFA-6 enriched cortical domains could be associated with LET-805-containing attachment structures. However, unlike EFA-6, which exhibits IDR-dependent enrichment at specific neuron-epidermal interfaces, LET-805 is broadly distributed throughout the epidermis, where it forms punctate structures associated with ECM attachment. This distinction suggests that EFA-6 enrichment is unlikely to arise solely from the presence of epidermal attachment structures. Instead, these observations raise the possibility that the ECM establishes spatially distinct cortical environments that independently recruit EFA-6 and organize epidermal attachment complexes, thereby coordinating cortical organization with neuron-epidermal interactions.

Collectively, these studies suggest a model in which epidermal cortex is organized in relation to ECM organization at the neuronal-epidermal interfaces to coordinate epithelial membrane cytoskeleton with neurite ensheathment. Future studies examining localized microtubule cytoskeleton changes at the interfaces during ensheathment and the impact of this organization on ECM dependent modeling, mechanical and functional coupling, of these interfaces would be crucial to test this model.

## METHODS

### C. elegans genetics

All experiments were conducted using *Caenorhabditis elegans*. Strains were maintained on standard nematode growth media (NGM) plates seeded with E. coli OP50 as described (Brenner, 1974). All strains were kept at 20°C, except where noted. Compound mutants with desired fluorescent reporters (KI or transgenes) were generated by genetic crosses; genotypes were confirmed using allele-specific polymorphisms or visualization of fluorescence. Strains and alleles are listed in Table S1-2, and primers for genotyping is in Table S3.

### Screen for altered GFP::EFA-6 patterning

To identify regulators of endogenous EFA-6, we performed a forward genetic screen using CZ26544 GFP::efa-6(ju1658); P*mec-7*-mKate2(*jsIs973*). L4 animals were treated with 47 mM Ethyl Methane Sulphonate (EMS) diluted in M9 buffer for 4 h following standard procedures (Brenner, 1974). P0 animals were washed with M9 buffer and recovered for 1 h at room temperature. 3-5 P0 animals were transferred to freshly seeded plates to produce F_1_ progeny. About 1200 F_1_s were subsequently transferred to 360 fresh seeded plates (about 3 F_1_/plate) to produce F_2_ progeny. About 20 F_2_s per F_1_ plate were screened at L4 stage by mounting on a 5% agarose pad with 2 mM levamisole and observed under Zeiss Axioplan 2 microscope equipped with Chroma HQ filters and a 63x objective. 12 independent isolates showing significant changes of GFP::EFA-6 pattern compared to non-mutagenized animals were found from the screens. Mutant isolates were outcrossed with either N2 or CZ26498 GFP::*efa-6*(ju1658).

### Whole-genome sequencing (WGS) analysis and identification of mutants

Genomic DNA for original isolates and outcrossed strains was prepared using a Gentra Puregene Kit (Qiagen). 20x coverage WGS data was obtained as 90 bp paired-end reads (BGI Americas). We mapped the raw reads to the C. elegans reference genome (WBcel235/ce11) using the Galaxy platform (http://usegalaxy.org). MiModd 0.1.9 (https://mimodd.readthedocs.io/ed/latest/#) was used to subtract mutations present in the parental strain (CZ26544), facilitating the identification of causative mutations. We verified linked SNPs in *mec-1(ju1688), him-4(ju1689), him-4(ju1687), him-4(ju1768)* by Sanger sequencing on multiple outcrossed strains.

### CRISPR/Cas9-mediated genome editing

#### AID insertion into GFP::EFA-6(ju1658)

CRISPR/Cas9-mediated genome editing was performed using the melting method as previously described (Ghanta & Mello, 2020). Target-specific crRNA (TCAGAAGTTGCATGGAGTGA) was designed using the Integrated DNA Technologies (IDT) design tool. To assemble the guide RNA complex, 0.5 µL of 100 µM target-specific crRNA was mixed with 1.5 µL of 100 µM tracrRNA (IDT) in a total volume of 5 µL and incubated at 95°C for 5 min. The mixture was cooled to room temperature for 5-10 min, followed by the addition of Cas9 protein (0.5 µl of 40 µM; MacroLab, UC Berkeley) and incubation at 25°C for 1-2 h. The repair template containing the AID sequence flanked by *efa-6* homology arms was amplified from plasmid XL21 using primers SD21595 and SD21596 and Phusion DNA polymerase (annealing at 60°C for 30 s). The PCR product was gel-purified using a Zymo Research kit (11-301). Subsequently, 5 µL of the purified PCR product (∼200 ng/µL) was subjected to the melting procedure by heating at 95°C for 5 min, followed by stepwise cooling in 5°C decrements every 30 s until reaching 30°C. Immediately before injection, 1 µL of the melted repair template was added to the crRNA-tracrRNA-Cas9 mixture. The injection mix also contained 1 µL of pCFJ90 (*Pmyo-2:mCherry*; final concentration, 3 ng/µL) as a co-injection marker. The injection mixture was introduced into the gonads of approximately 12 P0 animals carrying GFP::EFA-6 (CZ27027). Injected animals were transferred to six seeded nematode growth medium (NGM) plates (two animals per plate) and maintained at room temperature for 3-4 days. F_1_ progeny from plates exhibiting the co-injection marker were individually isolated and screened for successful genome editing by PCR using primers SD23144 and SD23145. Correct AID insertion was identified by an increase in the PCR amplicon size from 365 bp to 509 bp, corresponding to the 144 bp AID insertion corresponding to *ju2193*. Addition of the AID tag did not alter the overall fluorescence intensity or spatial distribution of GFP::EFA-6 compared with the parental GFP::EFA-6 strain (Fig. S3C, D). Furthermore, GFP::EFA-6::AID animals did not exhibit the PLM neurite overgrowth phenotype associated with loss of *efa-6* function (Fig. S3E; (Chen et al., 2011; Chen et al., 2015)), indicating that the AID-tagged protein retains its function in regulating neurite growth.

#### HIM-4::mNG editing

Genome editing was performed using the CRISPR/Cas9 co-conversion strategy with *dpy-10(cn64)* as a phenotypic marker, as previously described (Arribere et al., 2014). sgRNAs were selected using IDT database are listed in Table S6. To generate deletions, 0.5 µL of each target-specific crRNA (200 µM) was mixed with 1.5 µL of tracrRNA (100 µM, IDT) and 0.5 µL of *dpy-10* crRNA (100 µM). The mixture was incubated at 95°C for 5 min and cooled to RT for 5-10 min. 0.5 µL of the annealed crRNA/tracrRNA mixture was combined with 2-3 µL of purified Cas9 protein (Macrolabs, UC Berkeley) and incubated the mixture at RT for 1-2 h to form ribonucleoprotein (RNP) complexes. Immediately before injection, the homology-directed repair (HDR) template for *dpy-10(cn64)* was added to the RNP mixture. For single-nucleotide editing, additional HDR templates of ∼200 bp were designed to introduce the desired base substitution along with silent mutations disrupting the crRNA recognition sequence, PAM site172 and introducing or abolishing a restriction enzyme site for genotyping. The injection mix was prepared as described above, with the addition of 4 µM of the mutation-specific HDR template to the final crRNA-tracrRNA-Cas9 mix. About 15 day 1 adult P_0_ animals were injected in each experiment and transferred to six seeded NGM plates (2-3 animals/plate) to propagate for 3-4 days at 20°C. From plates where Dpy or Rol phenotypes were observed, multiple F_1_ progeny were isolated and screened for correct editing using PCR coupled with enzyme digestion. All edited alleles were verified by Sanger sequencing and outcrossed before further analyses. Sequence changes and genotyping information are in Table S2, 6.

### Auxin mediated EFA-6 depletion

A 5 mM indole-3-acetic acid (IAA) working solution was freshly prepared before each experiment by diluting a 400 mM IAA stock solution (prepared in ethanol and stored at -20°C in the dark for up to one month) with water. For IAA treatment throughout development, 300 µL of the 5 mM IAA solution was evenly distributed over the surface of seeded nematode growth medium (NGM) plates. Plates were kept in the dark at room temperature overnight to allow the solution to dry. Control plates were prepared similarly using an equivalent volume of water instead of IAA solution. On day 1, four to five P_0_ animals at the L4 or young adult stage were transferred onto each plate and allowed to lay eggs overnight. On day 2, the P_0_ animals were removed, and the progeny were allowed to develop on treatment or control plates at room temperature for an additional two days (48-52 h). Animals were imaged at L4 stage.

To perform AID dependent depletion, tissues specific TIR1 transgenes were crossed with GFP::EFA-6(AID). TIR1 expression in all cells or in neurons did not affect GFP::EFA-6(AID) expression or overall morphology of animals in the absence of IAA. However, epidermal TIR1 expression mildly reduced GFP::EFA-6 fluorescence in the absence of IAA, accompanied by developmental delay and reduced body size, both of which became more pronounced following IAA treatment. Although these animals exhibited elevated baseline PLM overgrowth, the absence of further enhancement upon IAA treatment suggests that this phenotype may reflect indirect effects associated with reduced body size or genetic background rather than epidermal EFA-6 depletion itself.

### Fluorescence microscopy and image acquisition

#### Confocal microscopy

Animals were anesthetized in 5 mM levamisole prepared in M9 buffer and mounted on 4% agar pads for imaging. Unless otherwise specified, confocal images were acquired using a Zeiss LSM 800 microscope (Axio Observer.Z1/7) equipped with an α Plan-Apochromat 63×/1.40 NA oil-immersion objective. The pinhole was set to 100 µm, with detector gains of 680 V and 650 V for GFP or mNG and RFP, respectively, and a digital gain of 1.0 for both channels. Images were acquired as z-stacks of 7-8 optical sections at 0.5-µm intervals. Images were processed and analyzed using ImageJ (Fiji) and are presented as single optical sections unless otherwise indicated in the figure legends.

For imaging GFP::EFA-6(ju1658) knock-in and mNG-tagged EFA-6 single-copy insertion strains, 488 nm and 561 nm laser powers were set to 0.5% and 1%, respectively. For extrachromosomal transgenic strains, the 488 nm laser power was reduced to 0.2%, and z-stacks comprising 13 optical sections spanning 6 µm were acquired. For the quantitative analyses presented in Supplementary Fig. 18H,I, laser power was further reduced to 0.1%, and z-stacks comprising 15 optical sections spanning 14 µm were acquired.

For imaging ALM neurite morphology in animals aged five days beyond the L4 stage (D5A), images were acquired using an EC Plan-Neofluar 40×/1.30 NA oil-immersion objective on the Zeiss LSM 800. The pinhole was set to 100 µm, with detector gains of 680 V and 650 V for GFP and RFP, respectively, and a digital gain of 1.0 for both channels. Z-stacks comprising 20 optical sections spanning 4.75 µm were acquired and displayed as sum-intensity projections generated using ImageJ (Fiji).

For HIM-4::mNG expression analysis in *efa-6(0) mutants (for Fig. S10)*, HIM-4::mNG and Pmec-7:RFP were imaged using 488 nm and 561 nm laser excitation at 3% and 1% power, respectively. Z-stacks comprising 13 optical sections spanning 3 µm were acquired, and representative single optical sections are presented as indicated in the figure legends.

#### Airyscan microscopy

For developmental analysis of GFP::EFA-6 spatial patterning, animals were imaged using a Zeiss LSM 900 microscope (Axio Observer.Z1/7) equipped with Airyscan in super-resolution mode and an α Plan-Apochromat 100×/1.46 NA oil-immersion objective. Laser power was optimized for individual fluorescent markers and maintained consistently across samples expressing the same marker. GFP::EFA-6 was imaged using 488-nm laser excitation at 3% power with the 100× objective and 2% power with the 63× objective. Pmec-7-mRFP (*jsIs973*) was imaged using 561-nm laser excitation at 5% power with both objectives, with a detector gain of 750 V for imaging at 63× magnification. Pmyo-3-mCherry (*juEx4078*) was imaged using 561-nm laser excitation at 3% power and a detector gain of 750 V with the 100× objective. For tissue-specific split-GFP experiments, reconstituted EFA-6-associated GFP fluorescence was imaged using 488-nm laser excitation at 4.5% power, while Pmec-7-RFP was imaged using 561 nm laser excitation at 5% power.

For simultaneous imaging of HIM-4::mNG, mSc::EFA-6, and Pmec-4-BFP, images were acquired using Airyscan with a 63×/1.40 NA oil-immersion objective. The pinhole was set to 224 µm, with a detector gain of 850 V and a digital gain of 1.0. HIM-4::mNG and mScarlet::EFA-6 were imaged using 488 nm and 570 nm laser excitation, respectively, at 2% power. Pmec-4::BFP was imaged using 387 nm laser excitation at 2% power, with a pinhole setting of 221 µm. Z-stacks comprising 20 optical sections spanning 4.75 µm were acquired, and representative single optical sections of the neurites are presented as indicated in the figure legends.

For imaging HIM-4::mNG distribution in the head, vulva, and germline, z-stacks comprising 20 optical sections spanning 9.5 µm were acquired using 488 nm laser excitation at 2% power and default detector gain settings.

#### Compound scope imaging

Standard compound fluorescence imaging was performed on Zeiss Axioplan M2 microscope equipped with Chroma HQ filters and alpha PlanApochromat 40x/1.40 oil immersion objective, with exposure time of 500 ms. We traced the ALM neurites at distal tip or proximal region posterior to the pharynx using segmented line. Next, we converted these traces to straightened cropped sections using the built-in ’Straighten’ function in Fiji.

### Quantitative image analysis

#### Quantification of GFP::EFA-6 fluorescence intensity and foci in confocal images

GFP::EFA-6 fluorescence intensity was quantified using ImageJ (Fiji). For confocal images, a 40 µm long segmented line with a width of 25 pixels was drawn along the distal and proximal regions of interest (ROIs), as defined in Fig. 1A. Mean fluorescence intensity was calculated by averaging the intensity values obtained from each line-scan profile. Each measurement represented an individual animal.

The same line-scan profiles were used to quantify EFA-6 foci. To account for differences in fluorescence intensity between samples, the minimum intensity value within each line-scan profile was subtracted from all intensity measurements. The resulting values were then normalized to the mean intensity of the background-subtracted profile, such that a normalized intensity of 1 represented the mean fluorescence intensity of the corresponding profile. Foci were defined as discrete fluorescence peaks with a normalized intensity exceeding 0.8 over a continuous distance of at least 0.5 µm and reaching a maximum normalized intensity of at least 1.0. For samples exhibiting low fluorescence intensity (mean intensity < 30 arbitrary units) and low signal variability (standard deviation < 25 arbitrary units), foci were counted manually from the original fluorescence images using the same minimum focus size criterion of 0.5 µm. To measure GFP::EFA-6 intensity in the nerve ring, three ROI of 26 µm^2^ was drawn in the nerve ring in a confocal single slice and mean intensity was measured using ImageJ measurement tool. These measurements were plotted after averaging and subtracting the mean background intensity.

#### Quantification of GFP::EFA-6 enrichment at neuron-epidermal interface

To quantify fluorescence intensity of GFP::EFA-6 at the neuron-epidermal interface relative to the surrounding epithelial cytoplasm, a 20 µm long segmented line with a width of 2 pixels was drawn along the edge of the proximal ALM neurite or the distal neurite (Fig. 5J) using ImageJ. Corresponding measurements were obtained from the adjacent epithelial cytoplasm using ROIs of identical dimensions. Two ROIs each were measured at the ALM neuron-epidermal interface and in the epithelial cytoplasm per image. Measurements from the two ROIs within each compartment were averaged to obtain the mean interface and cytoplasmic fluorescence intensities for each animal. These values were either plotted as absolute fluorescence intensities (Supplementary Fig. 18I) or used to calculate the ratio of interface to cytoplasmic fluorescence intensity as a measure of relative enrichment.

#### Quantification of GFP::EFA-6 fluorescence intensity and foci in compound microscope images

For images acquired using a compound microscope, GFP::EFA-6 fluorescence intensity was quantified using rectangular ROIs measuring 2 × 20 µm (40 µm^2^) in ImageJ (Fiji). For the distal region, one ROI per animal was positioned along the terminal 20 µm of the ALM neurite, extending proximally from the axonal tip. For the proximal region, three ROIs of identical dimensions were selected along the remaining neurite. For each ROI, an adjacent region of identical size and shape was selected for background subtraction. Background-corrected fluorescence intensities were used for subsequent quantitative analyses.

For quantification of GFP::EFA-6 foci in the distal neurite of ALM, ROIs were selected as described above. Line-scan profiles and particle analysis in ImageJ were used to facilitate the identification of individual foci. Each focus was manually outlined using the drawing tool in ImageJ, and its area was measured using the *Area* parameter. The total number of foci within each ROI was also recorded.

#### Quantification of HIM-4::mNG fluorescence intensity along ALM neurites

HIM-4::mNG fluorescence intensity was quantified from confocal and Airyscan images using ImageJ. A 20 µm long segmented line with a width of 25 pixels was drawn along the distal and proximal regions of the ALM neurite, as defined in Fig. 1A. Two ROIs were measured per image, and the mean fluorescence intensity from each ROI was plotted as an independent measurement.

#### Quantification of HIM-4::mNG fluorescence intensity in the head, vulva, and germline

For quantification of HIM-4::mNG fluorescence intensity in the head, vulva, and germline, Airyscan z-stacks were converted into sum-intensity projections using the *Z Project* function in ImageJ. For the head region, two ROIs of 100 µm^2^ each were selected within regions containing HIM-4-associated muscle attachment tracks. Fluorescence intensity was measured for each ROI, and background fluorescence from an adjacent region was subtracted. The resulting background-corrected intensity values were plotted independently. For the vulval region, two ROIs of 100 µm² each were selected, one on either side of the vulval musculature.

Fluorescence intensities were measured, and the values from individual ROIs were plotted independently. For the germline, three ROIs of 25 µm^2^ each, encompassing approximately two germ cells per ROI, were selected. Fluorescence intensity was measured, corrected by subtracting the corresponding background fluorescence, and plotted independently for each ROI.

### Developmental analysis of GFP::EFA-6 patterning

Animals of strain CZ29472 were synchronized by allowing 10-15 adult hermaphrodites to lay eggs for 2 h. Eggs were allowed to hatch and grow at 20 °C. Animals were collected at 26 h (L2), 42 h (L3), and 52 h (L4) and imaged using Airyscan microscopy as described above.

### Fluorescence recovery after photo bleaching (FRAP) analysis

FRAP experiments were performed using a Zeiss LSM 900 confocal microscope equipped with a 63×/1.40 NA DIC M27 objective. For all experiments, the selected regions of interest (ROIs) were photobleached using 15 iterations of the 488 nm laser at 100% power. All images during FRAP analysis were acquired with a default of pinhole setting of 1 Airy unit (44 µm), a detector gain of 700 V, and a digital gain of 1.0. For FRAP analysis of endogenous GFP::EFA-6 in Fig. 1, circular ROIs were selected at the distal tip or proximal region of the ALM neurite.

Fluorescence recovery was monitored at 10 s intervals for 8 min. For subsequent experiments involving endogenous GFP::EFA-6, a 3 × 3 µm ROI was selected for photobleaching (10 iterations), and fluorescence recovery was monitored at 5 min intervals for 15 min using 488 nm laser excitation at 2% power. Free GFP FRAP imaging was performed using a 488 nm laser excitation at 0.5% power. A rectangular ROI measuring 2 × 3 µm was selected for photobleaching, and fluorescence recovery was monitored over 10 s, with a total of 30 frames acquired. For epithelial-specific, single-copy transgenes, a 3 × 3 µm ROI was selected for photobleaching, and fluorescence recovery was monitored over 80 s, with a total of 10 frames acquired using 488-nm laser excitation at 2% power. For HIM-4::mNG, a 3 × 3 µm ROI was selected for photobleaching, and fluorescence recovery was monitored at 5 min intervals for 30 min using 488 nm laser excitation at 2% power. region, normalized to prebleach intensity, and expressed as a percentage.

Fluorescence recovery was quantified by subtracting background fluorescence measured from a region outside the animal, correcting for fluorescence loss during image acquisition using a nearby unbleached reference region, and normalizing the corrected fluorescence intensity to the corresponding prebleach value. Normalized fluorescence recovery was expressed as a percentage of the prebleach fluorescence intensity.

### Quantification of ALM and PLM neurite overshooting

Neurite overshooting quantification was performed as described in (Sandhu et al., 2024). Briefly, we used Pmec-7:RFP (jsIs973) to label TRN. Animals were anesthetized with 5 mM levamisole, mounted on 4% agar pads, and immediately examined using a compound microscope equipped with a 63× objective. In control animals, the anterior ALM neurite terminates within the hyp7 epidermal region, before reaching hyp5, approximately 30-40 µm from the nose tip. ALM neurites were scored as exhibiting an overshooting phenotype if their distal tips extended to the nose tip. Similarly, in control animals, the anterior PLM neurite terminates before reaching the ALM cell body. PLM neurites were scored as overshooting if their anterior tips extended beyond the ALM cell body. For each experiment, 20-30 animals were examined, and three independent biological replicates were performed.

### Quantification of ALM age-dependent neurite defects

L4-stage animals expressing zdIs5 or jsIs973 maintained at 15°C were transferred to nematode growth medium (NGM) plates containing 50 µM 5-fluoro-2′-deoxyuridine (FUdR), freshly seeded with 10-fold concentrated *E. coli* OP50 (modified from (Pan et al., 2011)). Plates were maintained at room temperature. D5A (L4 + 5 days) animals were anesthetized with 5 mM levamisole, mounted on 4% agar pads, and examined using a compound microscope equipped with a 63× objective. 5-10% of control animals showed 1-3 swellings in ALM neurite. In Fig 7, we scored mutant animals exhibiting five or more neurite swellings along the entire ALM neurite as positive for the neurite swelling phenotype, whereas animals with fewer than five swellings were classified as negative. Ectopic neurite branching was quantified by counting the total number of ectopic branches along the entire ALM neurite. For each experiment, 20-30 animals were analyzed, and three independent biological replicates were performed.

### Quantification of ALM neurite ensheathment defects

ALM neurite ensheathment defects were assessed from compound microscope images of day 1 adult animals. In wild-type animals, the proximal ALM neurite separates from the epidermal ridge, with the transition identified by the termination of GFP::EFA-6-labeled circumferential epidermal bands. Neurite positioning relative to the epidermal ridge was assessed using Pmec-7-RFP to label the ALM neurite. Ensheathment defects were classified into three categories based on the extent of neurite association with the epidermal ridge. Animals in which the ALM neurite remained associated with the ridge along its entire length were classified as having a complete loss of ensheathment. Animals in which approximately two-thirds of the neurite remained associated with the ridge were classified as having severe ensheathment defects. Animals in which approximately one-third of the neurite exhibited abnormal association with the ridge, typically at three to four discrete regions where the neurite failed to separate from the ridge, were classified as having partial loss of ensheathment.

### Quantification of vulval bursting

15-20 L4-stage animals were transferred to freshly seeded NGM plates containing *E. coli* OP50 and maintained at room temperature. Animals were examined the following day, and individuals exhibiting intestinal extrusion through the vulva were scored as positive for vulval bursting. The assay was repeated independently three to four times.

### Quantification of frequency of hermaphrodites producing male progeny

To assess the frequency of male progeny production, ten L4-stage hermaphrodites were individually transferred to freshly seeded NGM plates containing *E. coli* OP50 (one animal per plate). Plates were maintained at room temperature for three days and subsequently examined for the presence of male progeny. Plates containing at least one male were scored as positive, and the percentage of plates producing male progeny was calculated relative to the total number of plates examined.

### Generation of extrachromosomal transgenic arrays

Expression constructs encoding full-length EFA-6, its intrinsically disordered region (IDR), or the indicated IDR fragments and deletion variants were generated using corresponding cDNAs in-frame fused to mNG at N-terminus and under the control of the epidermal-specific *col-19* promoter (Pcol-19) (Ashley et al., 2021). Information of vector and ligation method are provided in Table S4.

Extrachromosomal transgenic arrays were generated by microinjection of plasmid DNA into the gonads of young adult *C. elegans* hermaphrodites using standard procedures (Mello et al., 1991). Injection mixtures contained the respective expression plasmids at 2 ng/µl, together with Papq-5-RFP co-injection marker at 50 ng/µl. Injected animals were transferred to nematode growth medium (NGM) plates seeded with *E. coli* OP50 and maintained at RT. Transgenic F_1_ progeny were identified based on expression of the co-injection marker, and stable extrachromosomal array lines were established by selecting marker-positive animals in subsequent generations. Transgenic strains, alleles, and plasmids used in this study are listed in Table S1-4.

### Generation of Cas9-mediated single-copy insertion (CasSCI) strains

Expression constructs encoding full-length EFA-6, its IDR, or the indicated deletion and truncation variants were cloned downstream of the epidermis-specific *col-10* promoter (Pcol-10) into CasSCI targeting vectors (Blazie & Jin, 2023; Dickinson et al., 2013; Frøkjær-Jensen et al., 2008). The resulting constructs were designed for insertion at the chromosomal IV genomic site (parent plasmid pCZGY2729). Single-copy transgenic strains were generated using the Cas9-mediated single-copy insertion (CasSCI) method (Blazie & Jin, 2023). CasSCI targeting plasmids were co-injected with Cas9/sgRNA plasmid (pCZGY2750) and co-injection marker plasmids, pCFJ90 (Pmyo-2-mCherry, 5 ng/µl) and pCFJ104 (Pmyo-3-mCherry, 10 ng/µl), into the gonads of young adult hermaphrodites of strain CZ30221[*efa-6(ju1883)*; Pmec-7:RFP(jsIs973)]. Injected animals were maintained at RT, and progeny carrying candidate insertions were identified using mNG expression under compound scope. Candidate single-copy insertion strains were isolated following loss of extrachromosomal co-injection markers and screened for correct genomic integration using PCR with locus-specific primers YJ10503, YJ10504, and YJ10686. The transgenic strains and corresponding alleles are listed in Table S1 and S2. Plasmids and primers used to generate these lines are listed in Table S3 and S5.

### Statistical Analysis

Statistical analyses were performed using GraphPad Prism 9 (GraphPad Software). Data distributions were assessed for normality to guide the selection of appropriate statistical tests. For comparisons between two groups, unpaired Student’s *t*-tests or nonparametric Mann-Whitney U tests were used, as appropriate. Multiple-group comparisons were performed using ordinary one-way ANOVA, Brown-Forsythe and Welch ANOVA, or Kruskal-Wallis tests followed by Dunn’s multiple-comparisons test, as appropriate. Time-course FRAP data were analyzed using repeated-measures two-way ANOVA with Geisser-Greenhouse correction. Categorical data were analyzed using Fisher’s exact test. Data are presented as mean ± standard error of the mean (SEM), unless otherwise indicated. Statistical significance was defined as *P* < 0.05. The statistical tests, sample sizes, and significance levels for individual experiments are specified in the corresponding figure legends.

## Supplemental Materials: 18 Supplemental Figures and 7 Supplemental Tables

Table S1. Strains used in the study

Table S2. Allele information and detection methods

Table S3. Genotyping, genome editing, and cloning primers

Table S4. Overexpression constructs generated for systematic analysis of the EFA-6 IDR

Table S5. Plasmids generated for CRISPR-assisted single-copy insertion (CasSCI) of EFA-6 constructs

Table S6. CRISPR/Cas9 genome editing information

Table S7. Summary of phenotypes associated with different *him-4* alleles

## Acknowledgements

We thank members of the Jin and Chisholm lab for helpful discussions and feedback throughout this study. We are grateful to Dr. Seungmee Park, Dr. Junxiang Zhou, and Dr. E. Jennifer Jin for sharing strains, plasmids, and protocols. We thank the Caenorhabditis Genetics Center (CGC), which is funded by the NIH Office of Research Infrastructure Programs (P40 OD010440), for providing *C. elegans* strains, and wormbase for resources. This work was supported in part by the National Institutes of Health (NIH), including an NIGMS K99/R00 Pathway to Independence Award (1K99GM160792 to A.S.), R35NS127314 to Y.J, and R35GM163671 to A.D.C.

## Declaration of Interests

The authors declare no competing interests.

## Author Contributions

AS, XL, XW, NHT, ADC, YJ designed the study; AS, XL, JS, XW, JX, KY, NHT conducted experiments; AS, XL, JS, XW, JX, KY, NHT, ADC, YJ analyzed data; AS, XL, ADC, YJ wrote the manuscript with all authors’ inputs.

## Resource Availability

All unique materials generated in this study, including strains and plasmids will be shared by the corresponding authors upon request.

## Notes

### Competing Interest Statement

The authors have declared no competing interest.

