## Supplemental Figures and Figure legends for "A specialized extracellular matrix patterns the conserved cortical protein EFA-6 at neuron-epidermal interfaces"

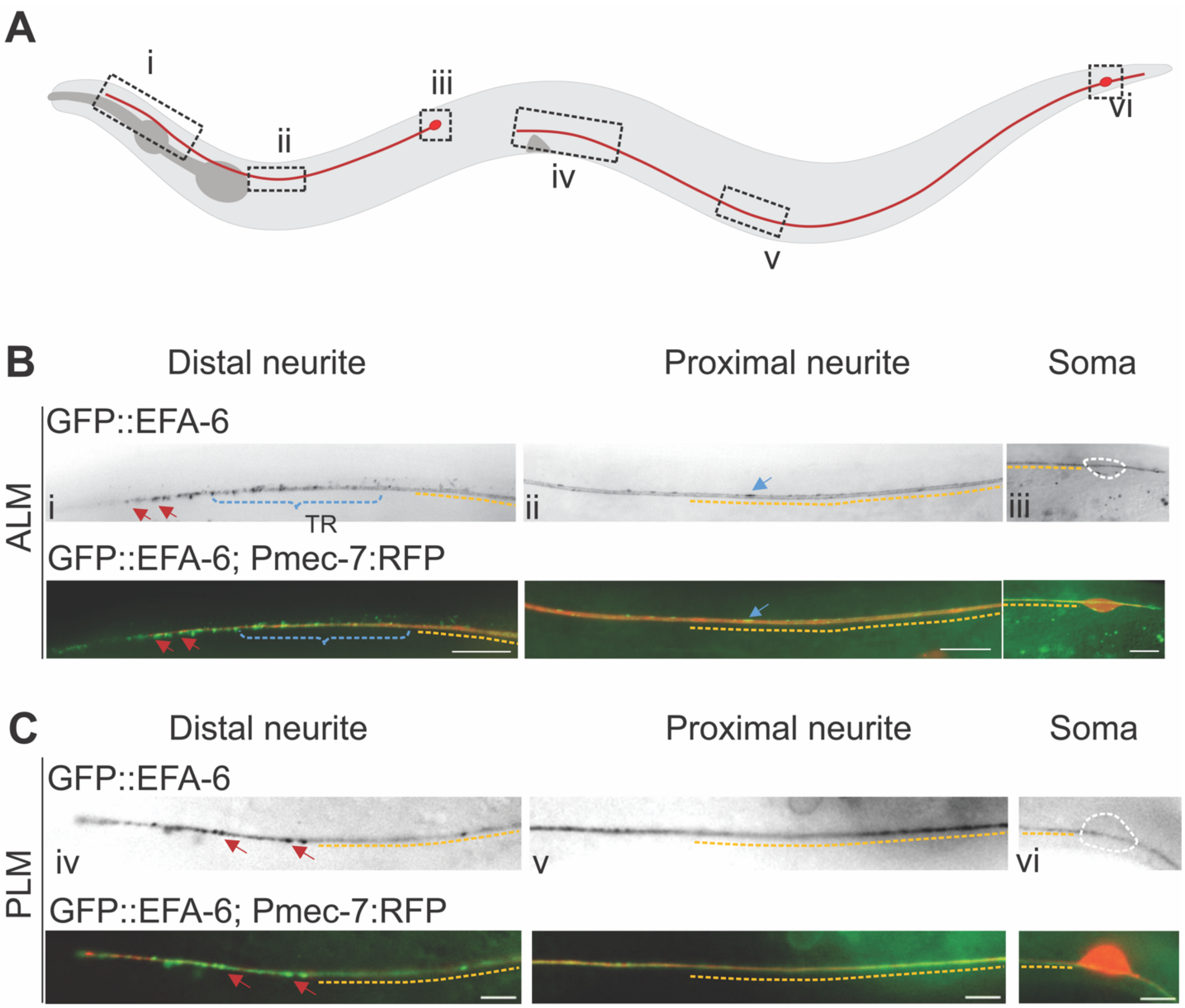


**Supplementary Figure 1: EFA-6 is differentially pattern in touch receptor neurons**

**(A)** Schematic representation of the *C. elegans* touch receptor neurons (TRNs), indicating the regions of interest (ROIs) corresponding to the images shown in subsequent panels: (i) distal ALM neurite, (ii) proximal ALM neurite, (iii) ALM soma, (iv) distal PLM neurite, (v) proximal PLM neurite, and (vi) PLM soma.

**(B-C)** Representative compound fluorescence images of adult animals expressing GFP::EFA-6 and Pmec-7:RFP, which labels the TRNs. The upper panels show GFP::EFA-6 fluorescence alone, whereas the lower panels show merged GFP and RFP channels. (B) Along the ALM neurite, GFP::EFA-6 forms distinct punctate foci in the distal ROI (red arrows) and gradually transitions to a more uniform distribution in the proximal ROI, with occasional elongated foci (blue arrows). The transition region (TR) is indicated by blue dotted lines. The ALM soma is outlined by a white dashed line, and the neurite trajectory is highlighted by an orange dotted line. (C) Along the PLM neurite, GFP::EFA-6 exhibits a spatial distribution similar to that observed along ALM, with distal punctate foci and a more diffuse proximal distribution, although this patterning is less pronounced. n = 10-15 animals per neuron type. Scale bars, 5 μm.

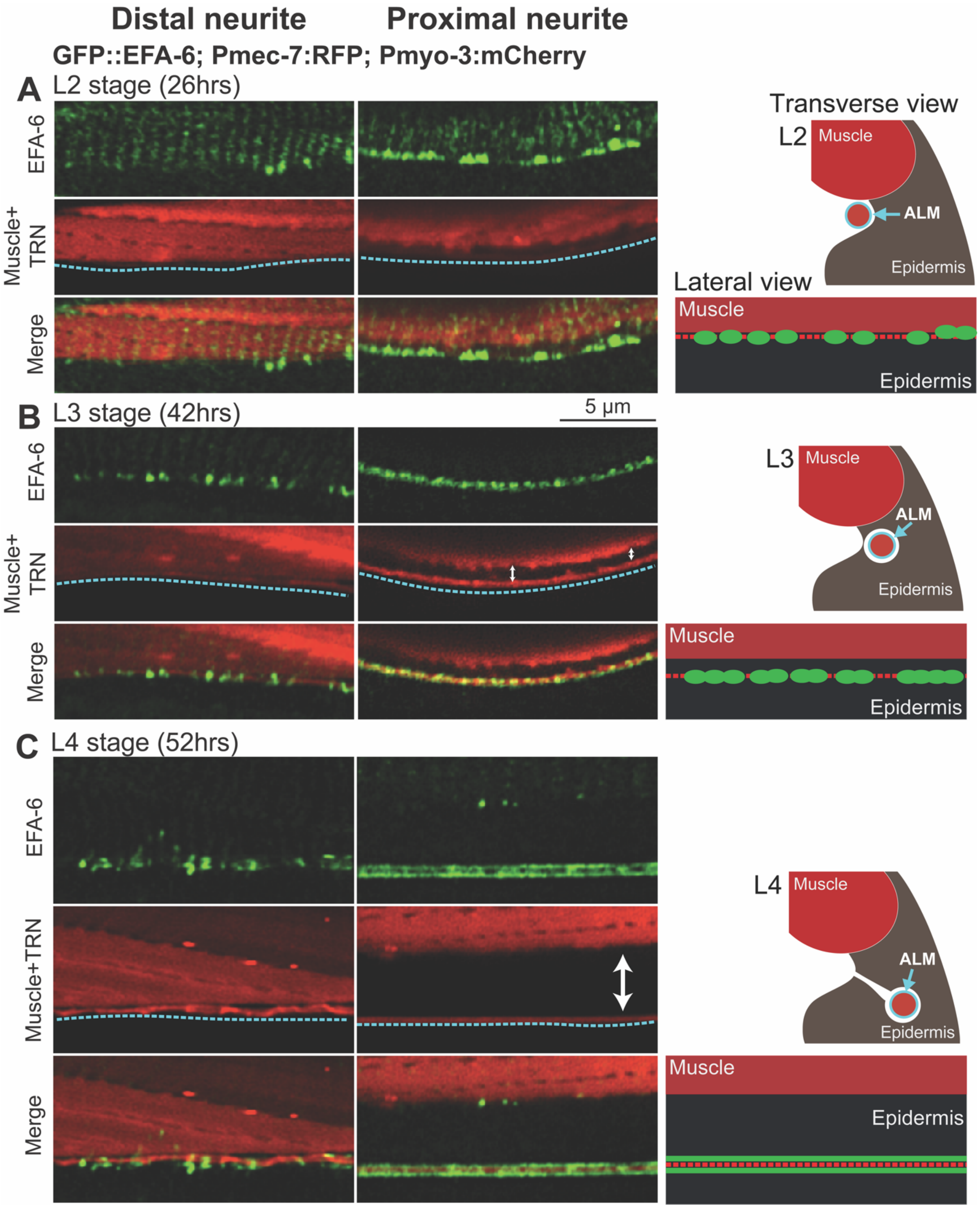

Distal neurite
Proximal neurite
GFP::EFA-6; Pmec-7:RFP; Pmyo-3:mCherry
A
L2 stage (26hrs)
EFA-6
Muscle+TRN
Merge
Transverse view
L2
Muscle
ALM
Epidermis
Lateral view
Muscle
Epidermis
B
L3 stage (42hrs)
5 μm
EFA-6
Muscle+TRN
Merge
L3
Muscle
ALM
Epidermis
Muscle
Epidermis
C
L4 stage (52hrs)
EFA-6
Muscle+TRN
Merge
L4
Muscle
ALM
Epidermis
Muscle
Epidermis

**Supplementary Figure 2: EFA-6 spatial patterning is developmentally correlated to neurite ensheathment.**

**(A-C)** Representative single-plane airyscan images of GFP::EFA-6 co-expressed with Pmec-7:mRFP (labeling ALM neurons) and Pmyo-3:mCherry (labeling body-wall muscle) at different developmental stages. Images show distal and proximal ROIs of ALM neurite (same as Fig.1A). The upper panels show the GFP channel alone, the middle panels show RFP channel, and the bottom panels show merged GFP and RFP channels. Shown on the right is the diagrams depicting transverse and lateral cross-sections of ALM (red circle marked with blue outline) position relative to the epidermis (marked in brown) and body-wall muscle (colored red) along the body axis at each larvae stage. Blue dotted line marks the position of ALM neurite in airyscan images. (A) At L2, the ALM neurite lies closely to the body-wall muscle at both distal and proximal ROIs. GFP::EFA-6 forms punctuated foci at both distal and proximal ROIs. (B) At L3, the distal ALM neurite remained adjacent to body-wall muscle. In contrast, the proximal neurite begins to exhibit separation from the body-wall muscle edge. Along the embedded proximal ROI, GFP::EFA-6 displays periodic and elongated foci, whereas in distal ROI non-embedded regions GFP::EFA-remains punctated. (C) At L4, the distal ALM neurite remained adjacent to body-wall muscle but a clear separation (indicated by white arrows) was observed at the proximal neurite. GFP::EFA-6 remains punctated in the distal ROI, whereas it forms a uniformly distributed expression pattern along proximal ROI. n=5-10 per stage; Scales: 5 μm.

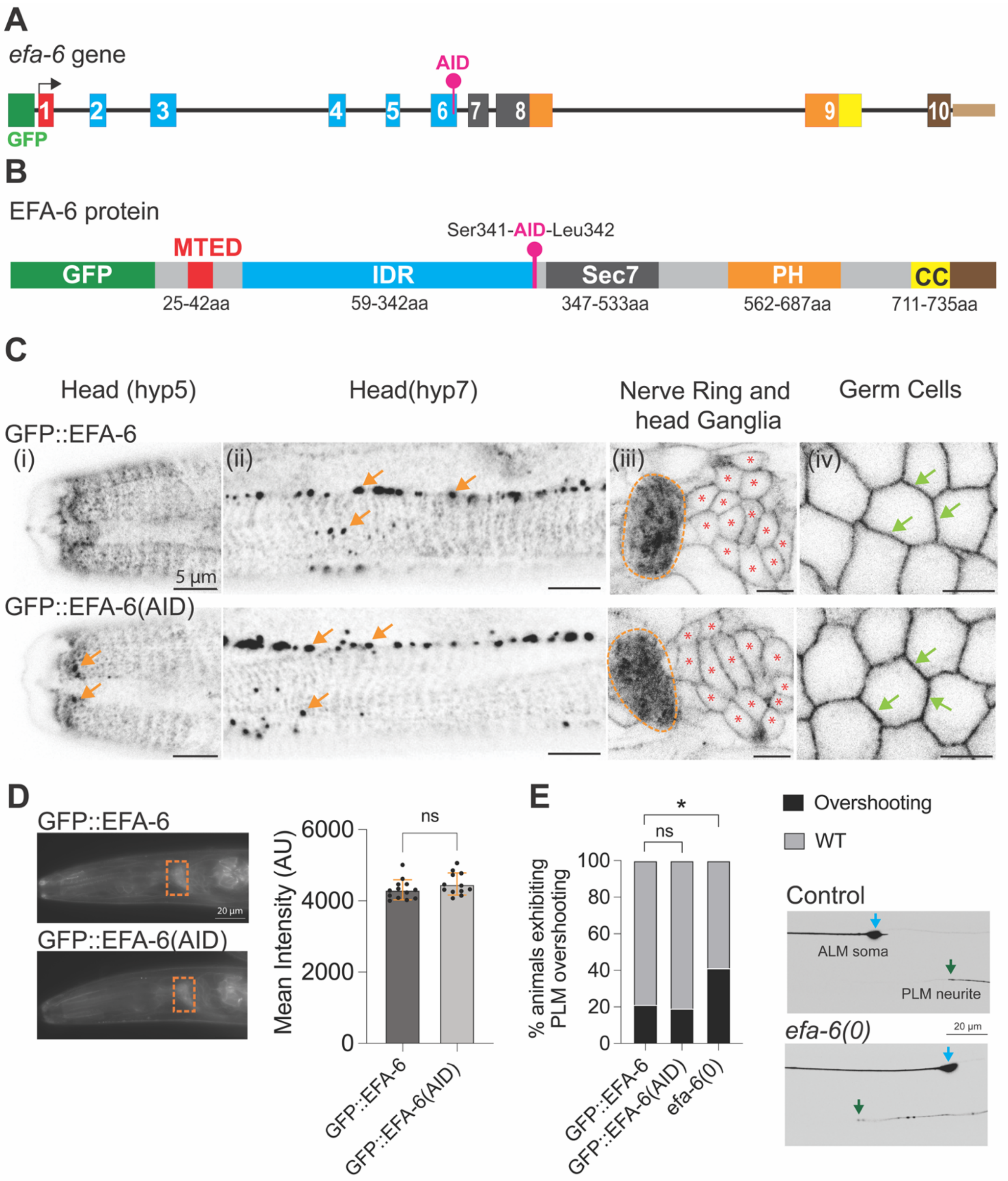

A
efa-6 gene
AID
GFP
B
EFA-6 protein
Ser341-AID-Leu342
MTED
GFP
IDR
Sec7
PH
CC
25-42aa
59-342aa
347-533aa
562-687aa
711-735aa
C
Head (hyp5)
Head(hyp7)
Nerve Ring and head Ganglia
Germ Cells
GFP::EFA-6
(i)
(ii)
(iii)
(iv)
5 µm
GFP::EFA-6(AID)
D
GFP::EFA-6
20 µm
GFP::EFA-6(AID)
Mean Intensity (AU)
ns
6000
4000
2000
0
GFP::EFA-6
GFP::EFA-6(AID)
E
% animals exhibiting PLM overshooting
*
ns
100
80
60
40
20
0
GFP::EFA-6
GFP::EFA-6(AID)
efa-6(0)
Overshooting
WT
Control
ALM soma
PLM neurite
efa-6(0)
20 µm

**Supplementary Figure 3: AID tagging at endogenous GFP::EFA-6 locus does not affect EFA-6 expression or function**

**A)** Illustration showing GFP::EFA-6 genomic organization. Boxes represent exons and introns are marked as connecting lines. Colors of boxes correspond to depicted domains of EFA-6. Green: GFP; Red: Microtubule elimination domain (MTED); Blue: Intrinsically disordered region (IDR); Dark grey: Sec7; Orange: Pleckstrin homology (PH) domain; Yellow: Coiled-coil (CC) domain; Brown: C-terminus region. Pink pin indicates the AID knock-in position.

**B)** Illustration showing GFP::EFA-6 protein architecture. EFA-6 domains are marked in the respective colors and their start to end positions are mention below each domain. Pink pin marks the AID knock-in position.

**C)** Confocal single-plane images of GFP::EFA-6 knock-in control and AID tagged GFP::EFA-6 knock-in in L4 stage animals. GFP::EFA-6(AID) shows expression pattern indistinguishable from control animals. EFA-6 exhibits both diffused and punctuated expression pattern in the head epidermis (i & ii; epidermal foci marked by orange arrows). EFA-6 remains uniformly distributed in the nerve ring (outlined with dotted orange lines) and head ganglia (iii) and germ cells (iv). Head ganglia neurons and cellular cortex of germ cells are marked by red asterisk and green arrows respectively. n=10 per genotype. Scale bar: 5 µm.

**D)** Quantitation of GFP::EFA-6 mean fluorescence intensity at the nerve ring region of control and AID tagged GFP::EFA-6 knock-in. Compound fluorescence images on the left shows the ROIs quantified for the analysis. Scale bar: 20 µm.

**E)** Quantitation of percentage of animals showing PLM overshooting. Confocal single-plane images on the right shows representatives for PLM overshooting compared to control. Blue arrow marks the ALM soma and green arrow indicates PLM anterior neurite tip. Scale bar: 20 µm.

**(D-E)** n=10-15 for (D) and n=30 for (E) per genotype. Data are presented as mean ± SEM. Statistics: Unpaired t-test for (D) and Fisher Exact test for (E); ns, not significant ($P > 0.05$); $^*P < 0.05$.

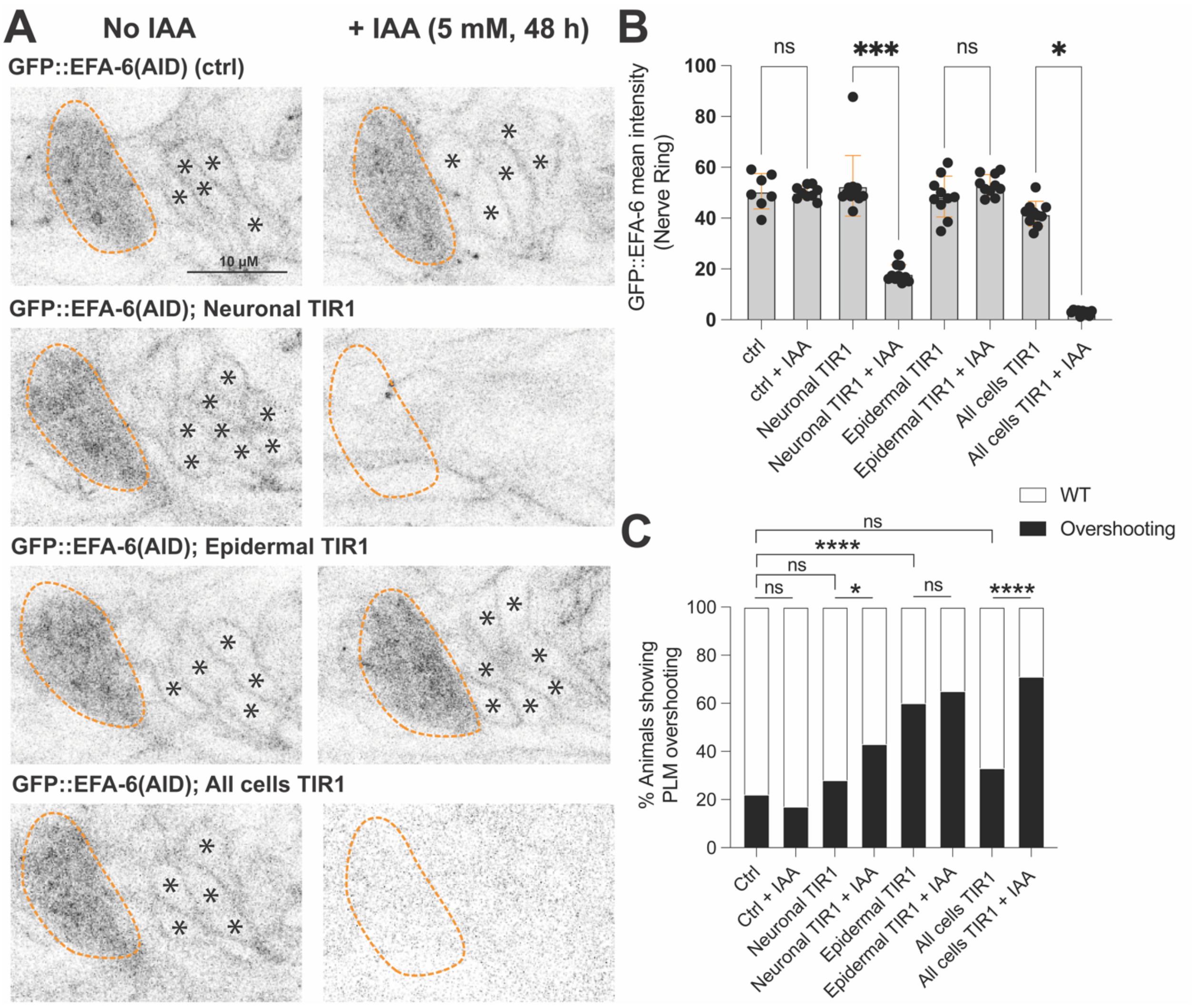


**Supplementary Figure 4: Auxin treatment efficiently depletes AID-tagged GFP::EFA-6 in neurons**

**(A)** Confocal single-plane images of endogenously AID tagged GFP::EFA-6 at the nerve ring (orange outline) and head ganglia neuronal soma (marked by black asterisks) in L4 animals, before and after indole-3-acetic acid (IAA; auxin) treatment for 48 h. Auxin treatment reduced GFP::EFA-6 levels at nerve ring and head ganglia region in animals expressing all cells (Peft-3:TIR1) and neuronal (Prgef-1:TIR1) TIR1. In contrast, GFP::EFA-6 expression remain unaltered in the nerve ring and head ganglia of epidermal (Pcol-10:TIR1) TIR1 expressing animals after auxin treatment. n=10 per genotype; Scale bar, 10 μm.

**(B)** Quantitation of GFP::EFA-6 mean fluorescence intensity at the nerve ring of genotypes indicated in panel A, before and after auxin treatment.

**(C)** Quantitation of percentage of animals showing PLM overshooting for genotype indicated panel A, before and after auxin treatment.

**(B, C)** n=10 for (B) and n=30 for (C) per genotype and condition. Data are presented as mean ± SEM. Statistics: Kruskal-Wallis test followed by Dunn's multiple-comparisons test for (B) and Fisher Exact test for (C); ns, not significant ($P > 0.05$); *$P < 0.05$; ***$P < 0.001$; ****$P < 0.0001$.

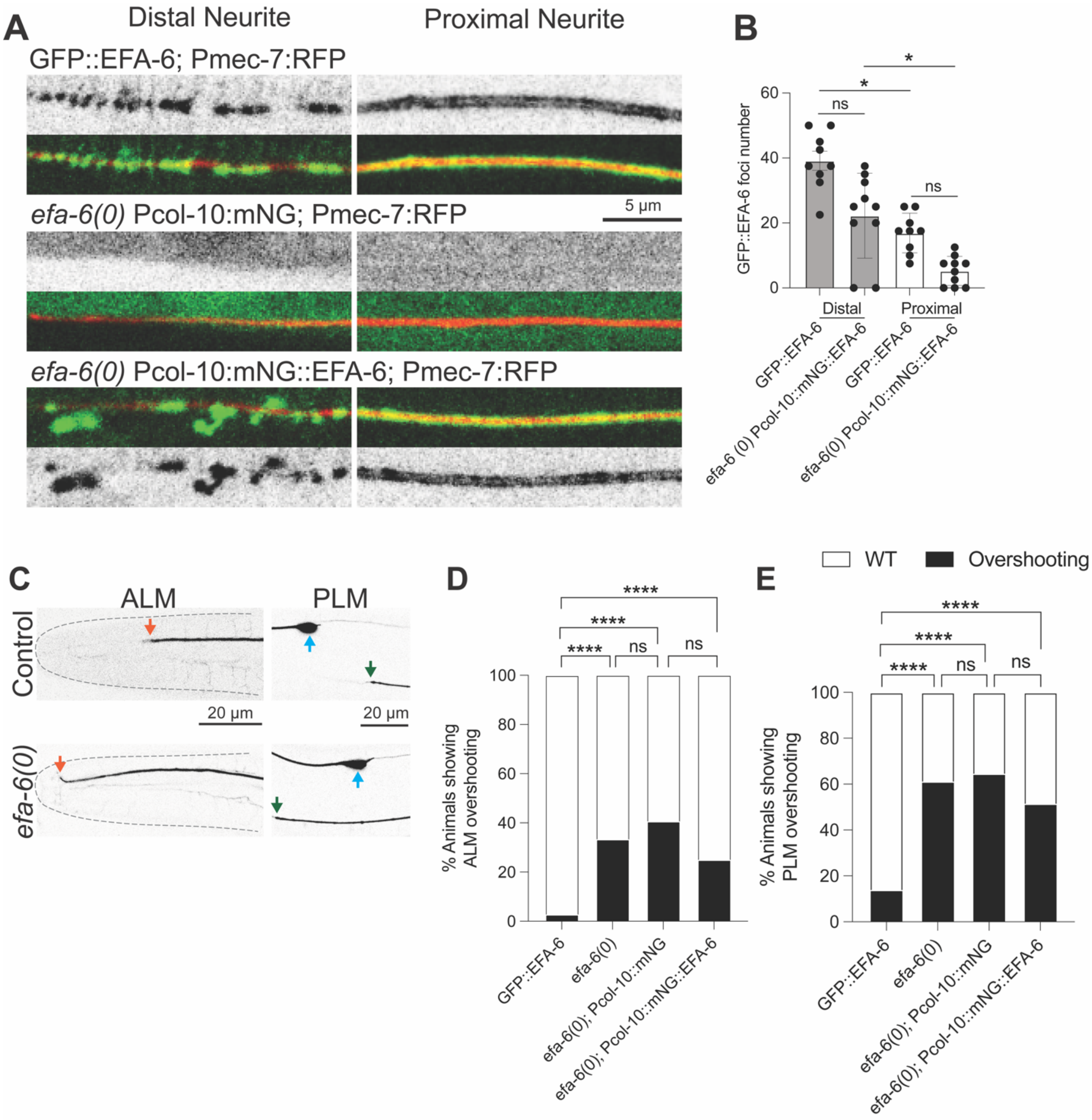


**Supplementary Figure 5: Epithelial EFA-6 recapitulates the endogenous EFA-6 spatial pattern along the ALM neurite.**

**(A)** Representative single-plane confocal images comparing endogenous GFP::EFA-6 knock-in (KI) with epithelial-specific (under Pcol-10) transgene of free mNG or mNG::EFA-6, expressed in the *efa-6(ju1883)* null background as single-copy insertion. Images show the distal and proximal ROIs (same as in Fig.1A) of the ALM neurite in L4 stage animals. To mark ALM neurite, merged images of mNG or GFP and Pmec-7:RFP labeling the neurite are shown in the bottom panel. In animals expressing epithelial free mNG, mNG is diffusely distributed throughout the epithelial cells and shows no specific enrichment along the ALM neurite. In *efa-6* null animals expressing epithelial mNG::EFA-6, EFA-6 recapitulates the spatial pattern of endogenous GFP::EFA-6. mNG::EFA-6 forms punctate foci at the distal ROI while exhibiting a predominantly uniform distribution along the proximal ROI. n=10 animals per genotype. Scale bar, 5 μm.

**(B)** Quantitation of foci density at the distal and proximal ROIs (same as in Fig,1A) in endogenous GFP::EFA-6 KI and epithelial-specific transgene expressing mNG::EFA-6 in the *efa-6* null background as single-copy insertion.

**(C)** Confocal single-plane images showing ALM and PLM overshooting representative images. Orange arrow marks the ALM anterior neurite tip, blue arrow marks the ALM soma, and green arrow indicates PLM anterior neurite tip. Scale bar: 20 μm.

**(D)** Quantitation of percentage of animals showing ALM overshooting in endogenous GFP::EFA-6 KI and epithelial-specific transgene expressing mNG::EFA-6 in the *efa-6* null background as single-copy insertion.

**(E)** Quantitation of percentage of animals showing PLM overshooting in endogenous GFP::EFA-6 KI and epithelial-specific transgene expressing mNG::EFA-6 in the *efa-6* null background as single-copy insertion.

**(B, D, E)** n=10 for (B) and n=30 per genotype for (D, E). Data are presented as mean ± SEM. Statistics: Kruskal-Wallis test followed by Dunn’s multiple-comparisons test for (B) and Fisher Exact test for (D) and (E); ns, not significant ($P > 0.05$); *$P < 0.05$; ****$P < 0.0001$.

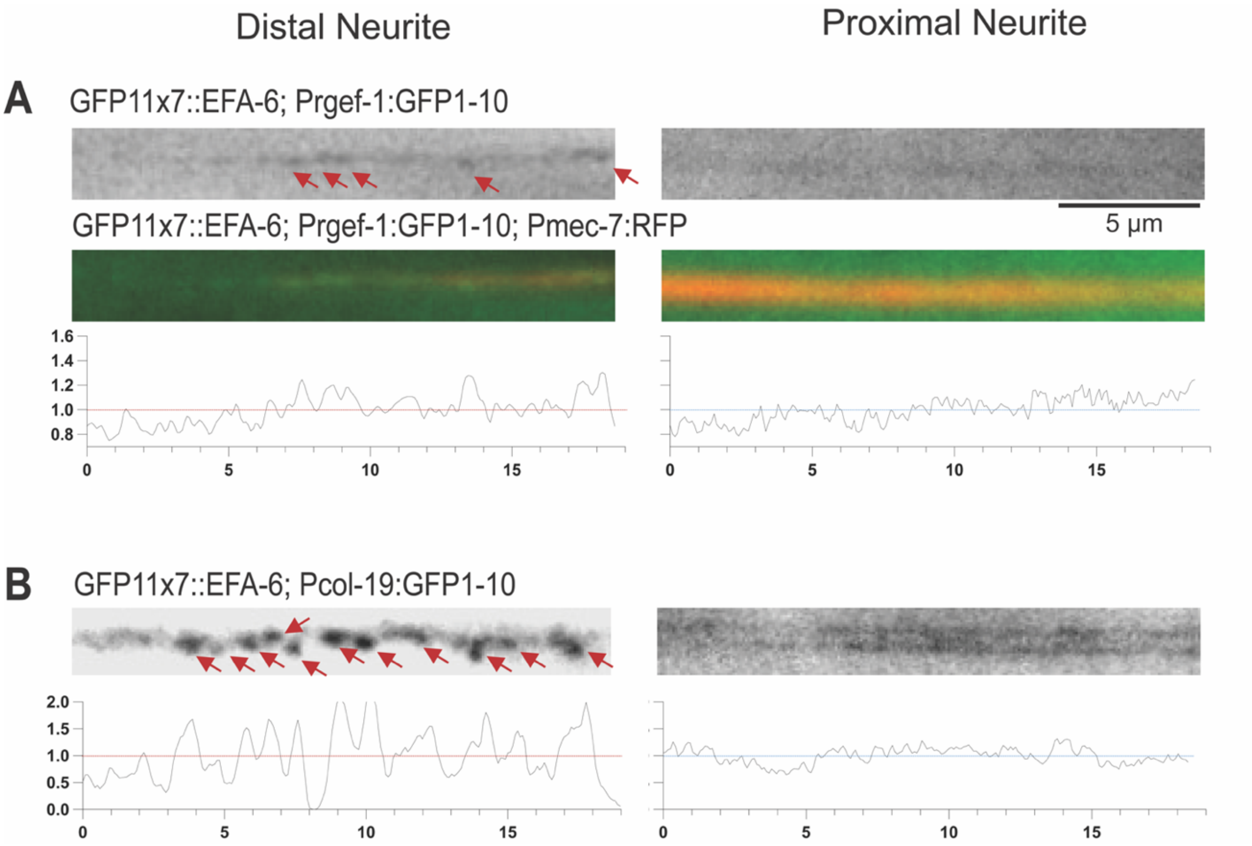


**Supplementary Figure 6: Neuronal and epidermal reconstituted GFP-11::EFA-6 phenocopies endogenous GFP::EFA-6 expression pattern.**

**(A-B)** Compound fluorescence images and line scan for tissue-specific split-GFP reconstitution of EFA-6 in the neuron (GFP11x7::EFA-6; Prgef-1:GFP1-10) and epidermis (GFP11x7::EFA-6; Pcol-19:GFP1-10), with and without neuronal marker Pmec-7:RFP. Images show distal and proximal neurite regions in adult animals. In both tissues, EFA-6 forms foci (red arrows) at distal neurite but exhibits uniformly distributed expression pattern at proximal neurite region. n=5-6 per genotype; Scales: 5µm.

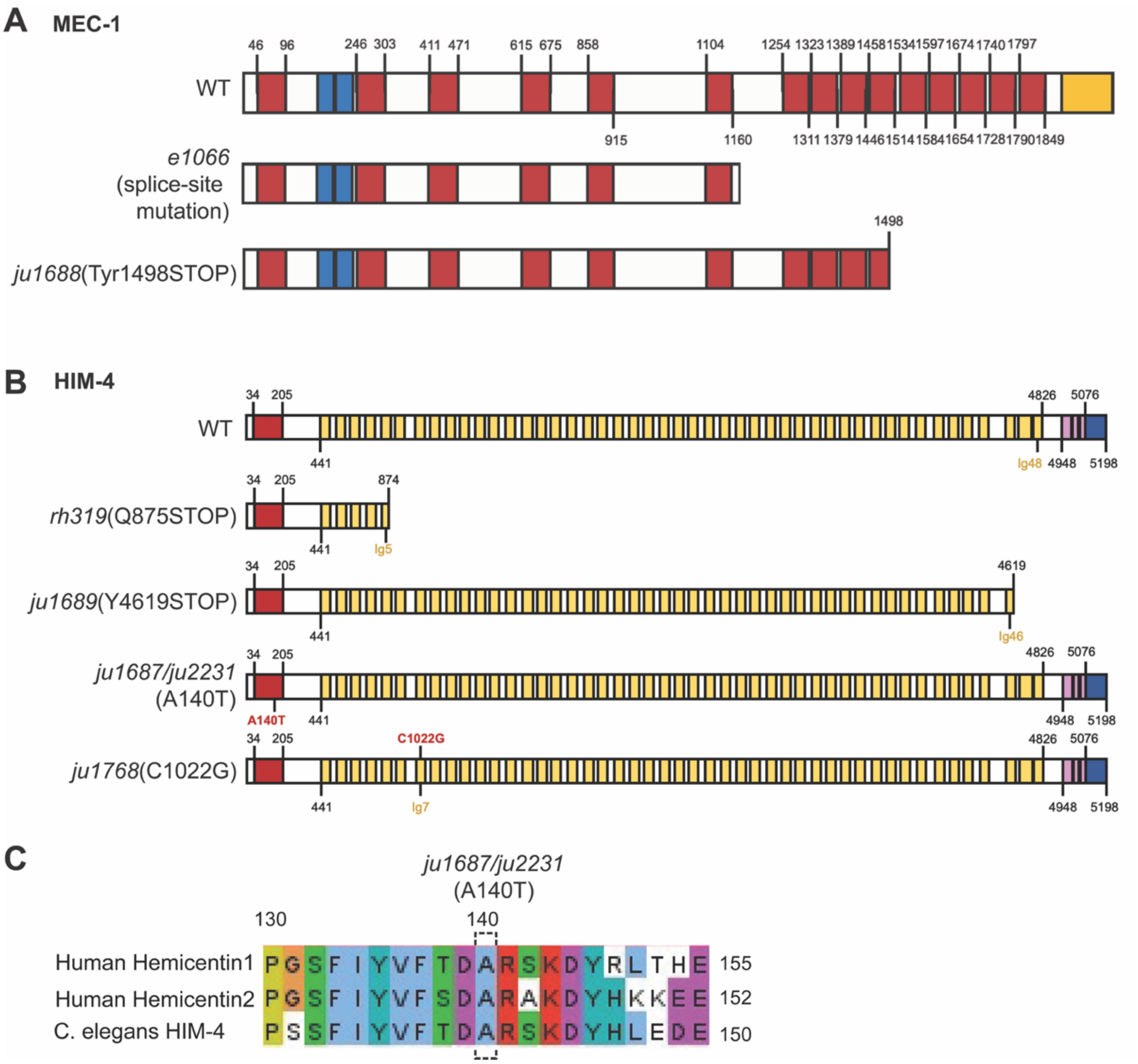


**Supplementary Figure 7. Domain organization of MEC-1 and HIM-4 and conservation of the HIM-4 VWA domain.**

**(A, B)** Schematic representations of the domain structure of the *C. elegans* extracellular matrix proteins MEC-1 (A) and HIM-4 (hemicentin) (B), showing mutations identified in the forward genetic screen and previously characterized loss-of-function alleles used in this study. (A) MEC-1 contains Kunitz (red), epidermal growth factor (EGF)-like (blue), and cysteine-rich C-terminal (yellow) domains. The predicted protein products of the *mec-1(e1066)* and *mec-1(ju1688/Tyr1498STOP)* alleles are illustrated. (B) HIM-4 contains an N-terminal von Willebrand factor type A (VWA) domain (red), multiple immunoglobulin (Ig)-like repeats (yellow), EGF-like domains (pink), and a C-terminal fibulin-type (FC) domain (blue). The positions of the A140T (ju1687/ju2231) and C1022G (ju1768) missense mutations, identified in the forward genetic screen, are indicated, along with the predicted truncated protein products of *him-4(rh319, Q875STOP)* and *him-4(ju1689, Y4619STOP)* and the C-terminal deletion allele *him-4(ju2236)*. Numbers indicate amino acid positions and domain boundaries. (C) Multiple sequence alignment of the VWA domain region containing the HIM-4 A140T substitution with the corresponding sequences of human hemicentin-1 (HMCN1) and hemicentin-2 (HMCN2). The conserved alanine residue affected by the A140T substitution is highlighted.

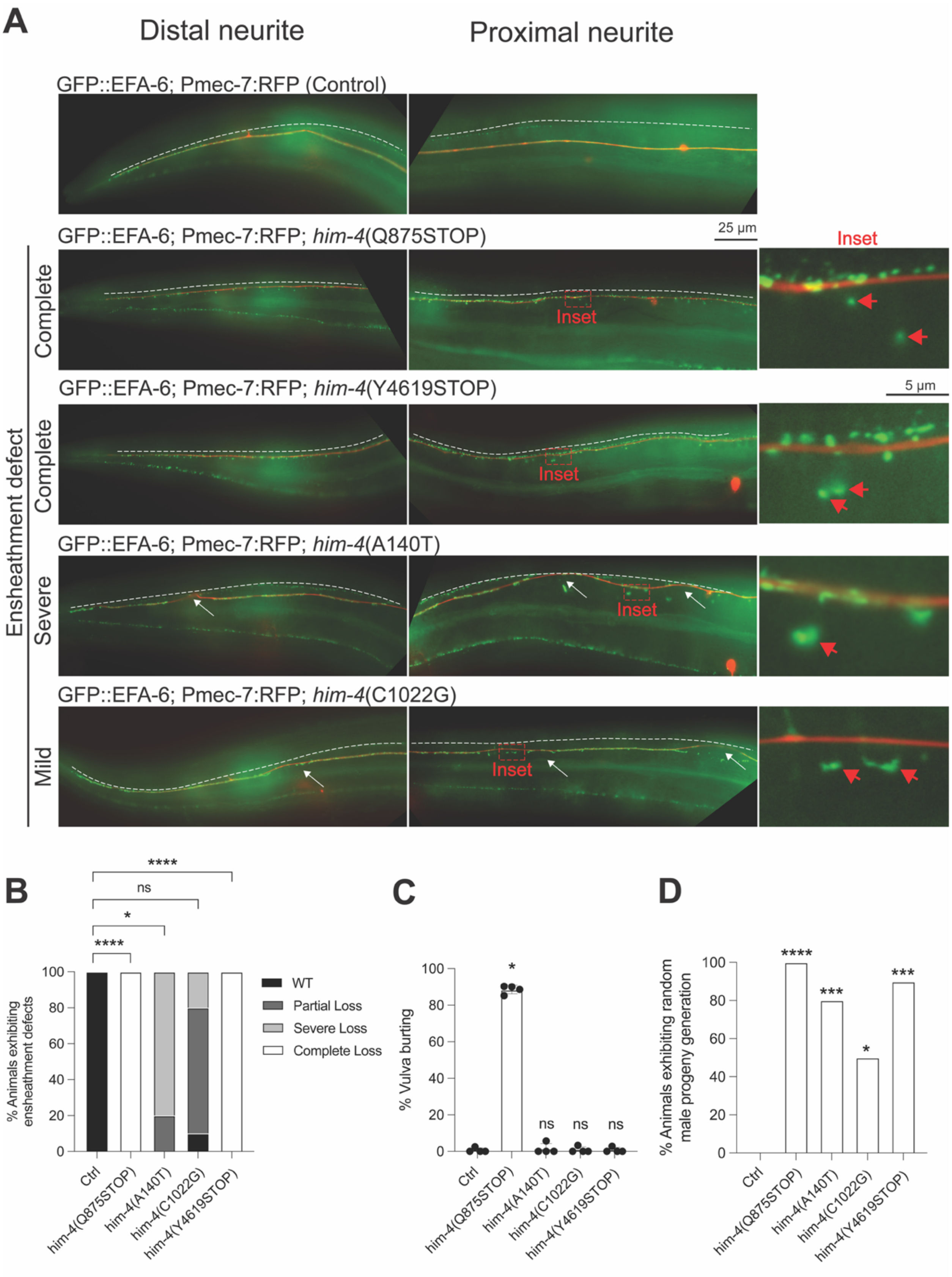
A
Distal neurite
Proximal neurite
GFP::EFA-6; Pmec-7:RFP (Control)
GFP::EFA-6; Pmec-7:RFP; him-4(Q875STOP)
25 μm
Inset
Complete
GFP::EFA-6; Pmec-7:RFP; him-4(Y4619STOP)
5 μm
Complete
GFP::EFA-6; Pmec-7:RFP; him-4(A140T)
Severe
GFP::EFA-6; Pmec-7:RFP; him-4(C1022G)
Mild
Ensheathment defect
B
% Animals exhibiting ensheathment defects
WT
Partial Loss
Severe Loss
Complete Loss
****
ns
*
****
Ctrl
him-4(Q875STOP)
him-4(A140T)
him-4(C1022G)
him-4(Y4619STOP)
C
% Vulva burting
*
ns
ns
ns
D
% Animals exhibiting random male progeny generation
****
***
*
***

**Supplementary Figure 8: HIM-4 mutations isolated from the forward genetic screen differentially disrupt distinct HIM-4-associated phenotypes.**

**(A)** Representative compound fluorescence images of adult animals expressing GFP::EFA-6 together with Pmec-7:RFP, which labels the ALM neurite. The genotypes shown include wild-type and *him-4(rh319, Q875STOP)* null controls, followed by three *him-4* alleles isolated from the forward genetic screen: *him-4(ju1689,* Y4619STOP), *him-4(ju1687,* A140T), and *him-4(ju1768,* C1022G). The dorsolateral epidermal ridge is indicated by the white dotted line, and the position of the ALM neurite relative to the ridge was used to assess neurite ensheathment. In wildtype control animals, the proximal ALM neurite is fully separated from the epidermal ridge, consistent with complete ensheathment. In the distal region, the anterior neurite tip remains closely associated with the ridge and progressively separates from it posteriorly. In contrast, in *him-4(rh319)* null mutants, the ALM neurite remains closely associated with the ridge throughout its length, indicating a complete loss of ensheathment. Among the screen-derived alleles, *him-4(ju1689,* Y4619STOP) phenocopies *him-4* null, with complete loss of ensheathment. *him-4(ju1687,* A140T) exhibits severe ensheathment defects, characterized by extended regions of the ALM neurite that remain associated with the ridge (marked by white arrows) interspersed with normally ensheathed regions. *him-4(ju1768,* C1022G) predominantly exhibits mild ensheathment defects, in which most of the neurite is normally ensheathed except for two to three discrete regions that remain associated with the ridge. Insets highlight EFA-6 foci displaced from the ALM neurite (red arrows). n=10 animals per genotype. Scale bar, 25 μm.

**(B)** Quantification of animals showing ensheathment defects for genotypes shown in panel (A).

**(C)** Quantification of animals showing vulva bursting defects for genotypes shown in panel (A).

**(D)** Quantification of percentage of animals exhibiting random male generation at RT for genotypes shown in panel (A).

**(B-D)** n=10 per genotype. Data are presented as mean ± SEM. Statistics: Kruskal-Wallis test followed by Dunn's multiple-comparisons test for (B-D); ns, not significant ($P > 0.05$); *$P < 0.05$; ***$P < 0.001$; ****$P < 0.0001$.

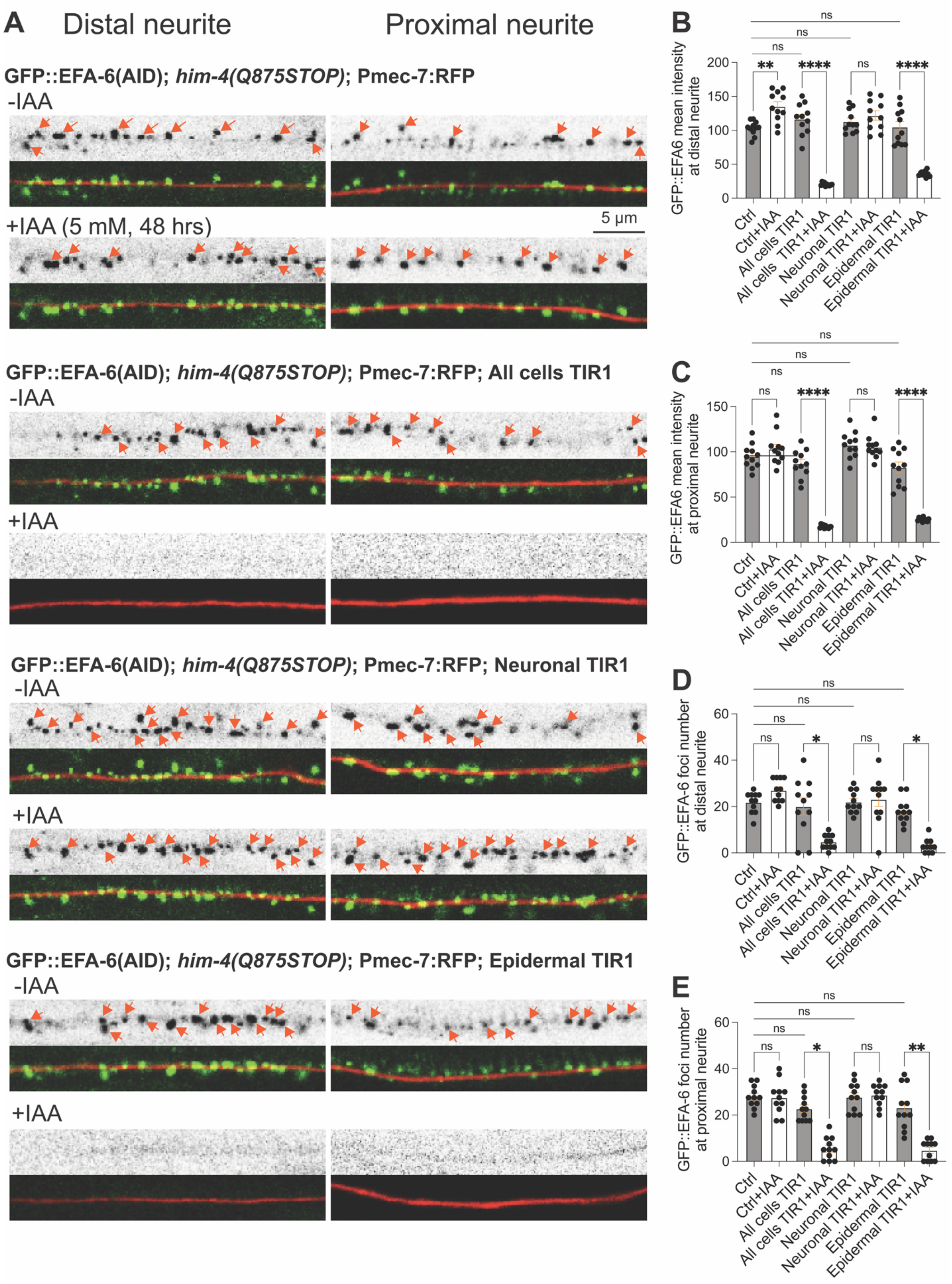

A
Distal neurite
Proximal neurite
GFP::EFA-6(AID); him-4(Q875STOP); Pmec-7:RFP
-IAA
+IAA (5 mM, 48 hrs)
5 μm
GFP::EFA-6(AID); him-4(Q875STOP); Pmec-7:RFP; All cells TIR1
-IAA
+IAA
GFP::EFA-6(AID); him-4(Q875STOP); Pmec-7:RFP; Neuronal TIR1
-IAA
+IAA
GFP::EFA-6(AID); him-4(Q875STOP); Pmec-7:RFP; Epidermal TIR1
-IAA
+IAA
B
GFP::EFA6 mean intensity at distal neurite
C
GFP::EFA6 mean intensity at proximal neurite
D
GFP::EFA-6 foci number at distal neurite
E
GFP::EFA-6 foci number at proximal neurite
Ctrl
Ctrl+IAA
All cells TIR1
All cells TIR1+IAA
Neuronal TIR1
Neuronal TIR1+IAA
Epidermal TIR1
Epidermal TIR1+IAA
ns
**
****
*

**Supplementary Figure 9: Ectopic EFA-6 foci associated with HIM-4 defects predominantly originate from epithelial cells.**

**(A)** Representative single-plane confocal images of endogenous AID-tagged GFP::EFA-6 (KI) in *him-4 (rh319, Q875STOP)* background. Images show distal and proximal ROIs (same as Fig.1A) of the ALM neurite (labeled with Pmec-7:RFP) in L4 animals before and after indole-3-acetic acid (IAA; auxin) treatment for 48 h. The upper panels show the GFP channel alone, whereas the lower panels show merged GFP and RFP channels. Orange arrows mark EFA-6 foci. Auxin treatment reduced GFP::EFA-6 levels at both distal and proximal neurite regions in animals expressing TIR1 in all cells (Peft-3:TIR1). Animals expressing neuronal TIR1 (Prgef-1:TIR1) showed little to no change in GFP::EFA-6 levels following auxin treatment. In contrast, epithelial TIR1 expression (Pcol-10:TIR1) resulted in a marked reduction in GFP::EFA-6 at both regions. n=10 animals per genotype. Scale bar, 5 µm.

**(B-C)** Quantification of mean GFP::EFA-6 fluorescence intensity at the (B) distal and (C) proximal regions of the ALM neurite in L4-stage *him-4(Q875STOP)* null animals expressing all cells, neuronal, or epithelial TIR1, before and after auxin treatment for 48hrs.

**(D-E)** Quantification of GFP::EFA-6 foci density at the (D) distal and (E) proximal ROIs (same as Fig. 1A) of the ALM neurite in L4-stage *him-4(Q875STOP)* null animals expressing all cells, neuronal, or epithelial TIR1, before and after auxin treatment for 48hrs.

**(B-E)** n=10-12 animals per genotype; Data are presented as mean ± SEM. Statistics: Brown-Forsythe and Welch ANOVA tests for (B-C) and Kruskal-Wallis test followed by Dunn's multiple-comparisons test for (D-E); ns, not significant ($P > 0.05$); *$P < 0.05$; **$P < 0.01$; ****$P < 0.0001$.

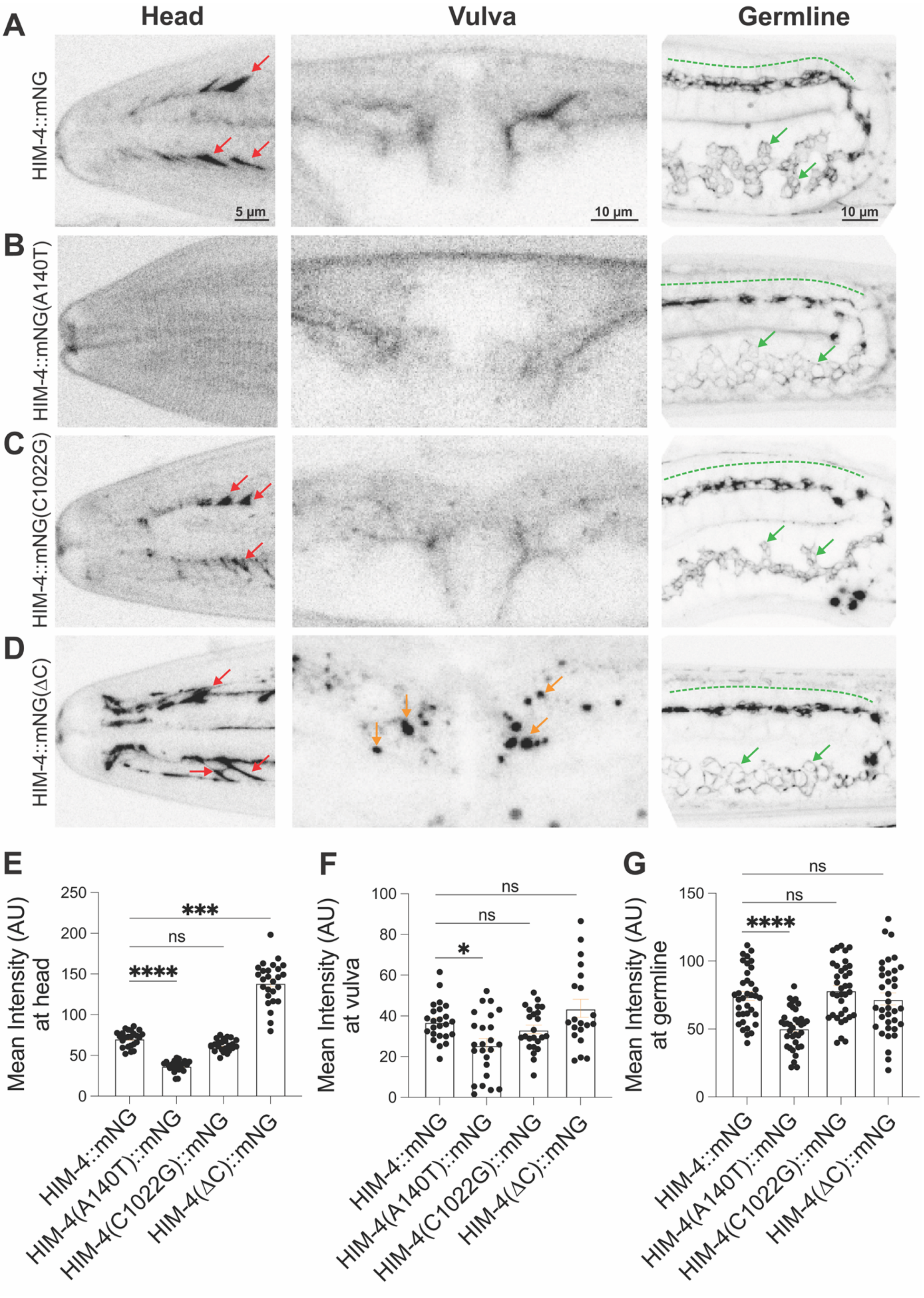

A
B
C
D
Head
Vulva
Germline
HIM-4::mNG
HIM-4::mNG(A140T)
HIM-4::mNG(C1022G)
HIM-4::mNG(ΔC)
5 μm
10 μm
10 μm
E
F
G
Mean Intensity (AU) at head
Mean Intensity (AU) at vulva
Mean Intensity (AU) at germline
***
ns
****
*
****
HIM-4::mNG
HIM-4(A140T)::mNG
HIM-4(C1022G)::mNG
HIM-4(ΔC)::mNG

**Supplementary Figure 10. HIM-4 mutations isolated from the forward genetic screen differentially disrupt HIM-4 expression and localization across tissues.**

**(A-D)** Representative airyscan z-projection images of HIM-4::mNG in L4 stage animals expressing (A) wild-type HIM-4::mNG (control), (B) HIM-4(A140T)::mNG (*ju2232*), (C) HIM-4(C1022G)::mNG (*ju2233*), or (D) HIM-4(ΔC)::mNG (*ju2235*), containing an in-frame C-terminal deletion. Images show HIM-4 organization in the head, vulva, and germline. In control animals, HIM-4::mNG forms distinct tracks in the head region (marked by red arrows). HIM-4(C1022G)::mNG shows no obvious alteration in these tracks, whereas the tracks are largely absent in HIM-4(A140T)::mNG animals. HIM-4(ΔC)::mNG retains track formation in the head, but their spatial organization is severely disrupted. At the vulva, control HIM-4::mNG exhibits a predominantly diffuse distribution throughout the vulval region. HIM-4(C1022G)::mNG shows no obvious alteration in vulval expression, whereas HIM-4(A140T)::mNG exhibits reduced HIM-4 signal. In HIM-4(ΔC)::mNG animals, HIM-4 exhibits a predominantly punctate (indicated by orange arrows) distribution at the vulva. In the germline, control HIM-4::mNG labels germ cells in the distal mitotic region (marked by green arrows) and forms a relatively diffuse sheet along the central rachis of the proximal gonad (indicated by green dotted line). HIM-4(A140T)::mNG exhibits reduced germline expression, whereas the C1022G and ΔC variants show no obvious reduction in germline expression. n=10 animals per genotype. Scale bars: head, 5 µm; vulva and germline, 10 µm.

**(E-G)** Quantification of mean HIM-4::mNG fluorescence intensity in the (E) head, (F) vulva, and (G) germline for the indicated HIM-4 variants. n = 10-12 animals per genotype. Data are presented as mean ± SEM. Statistics: Kruskal–Wallis test followed by Dunn's multiple-comparisons test. ns, not significant ($P > 0.05$); *$P < 0.05$; **$P < 0.01$; ***$P < 0.001$; ****$P < 0.0001$.

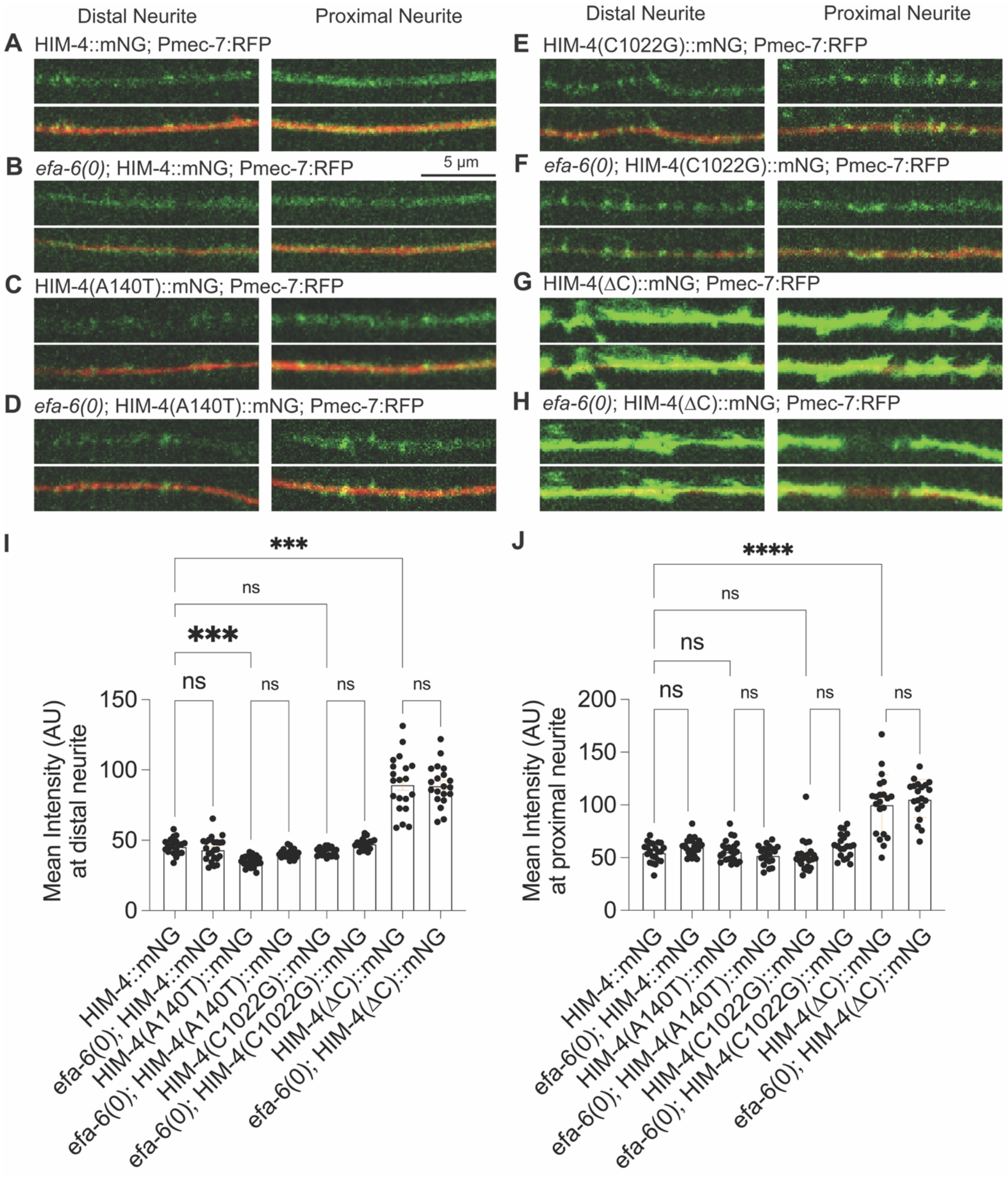
Distal Neurite
Proximal Neurite
A HIM-4::mNG; Pmec-7:RFP
B efa-6(0); HIM-4::mNG; Pmec-7:RFP
5 µm
C HIM-4(A140T)::mNG; Pmec-7:RFP
D efa-6(0); HIM-4(A140T)::mNG; Pmec-7:RFP
E HIM-4(C1022G)::mNG; Pmec-7:RFP
F efa-6(0); HIM-4(C1022G)::mNG; Pmec-7:RFP
G HIM-4(ΔC)::mNG; Pmec-7:RFP
H efa-6(0); HIM-4(ΔC)::mNG; Pmec-7:RFP
I
Mean Intensity (AU) at distal neurite
***
ns
***
ns
0
50
100
150
J
Mean Intensity (AU) at proximal neurite
****
ns
0
50
100
150
200
HIM-4::mNG
efa-6(0); HIM-4::mNG
HIM-4(A140T)::mNG
efa-6(0); HIM-4(A140T)::mNG
HIM-4(C1022G)::mNG
efa-6(0); HIM-4(C1022G)::mNG
HIM-4(ΔC)::mNG
efa-6(0); HIM-4(ΔC)::mNG

**Supplementary Figure 11. Loss of *efa-6* does not alter HIM-4 expression or organization along the ALM neurite.**

**(A-H)** Representative single-plane confocal images of wild-type HIM-4::mNG (control) and the indicated HIM-4 variants, HIM-4(A140T)::mNG, HIM-4(C1022G)::mNG, and HIM-4(ΔC)::mNG, in either wild-type or *efa-6(ju1883)* null backgrounds. Images show the distal and proximal ROIs (same as Fig.1A) of the ALM neurite, labeled with Pmec-7:RFP. The upper panels show the mNG channel alone, whereas the lower panels show merged mNG and RFP channels. Loss of *efa-6* does not detectably alter the expression level or spatial organization of wild-type HIM-4::mNG or the indicated HIM-4 variants along the ALM neurite. n=10 animals per genotype. Scale bar, 5 μm.

**(I, J)** Quantification of mean HIM-4::mNG fluorescence intensity at the (I) distal and (J) proximal ROIs (same as Fig.1A) of the ALM neurite for wild-type HIM-4::mNG and the indicated HIM-4 variants in wild-type or *efa-6* null backgrounds. n=10-12 animals per genotype. Data are presented as mean ± SEM. Statistics: Kruskal-Wallis test followed by Dunn's multiple-comparisons test. ns, not significant ($P > 0.05$); ***$P < 0.001$; ****$P < 0.0001$.

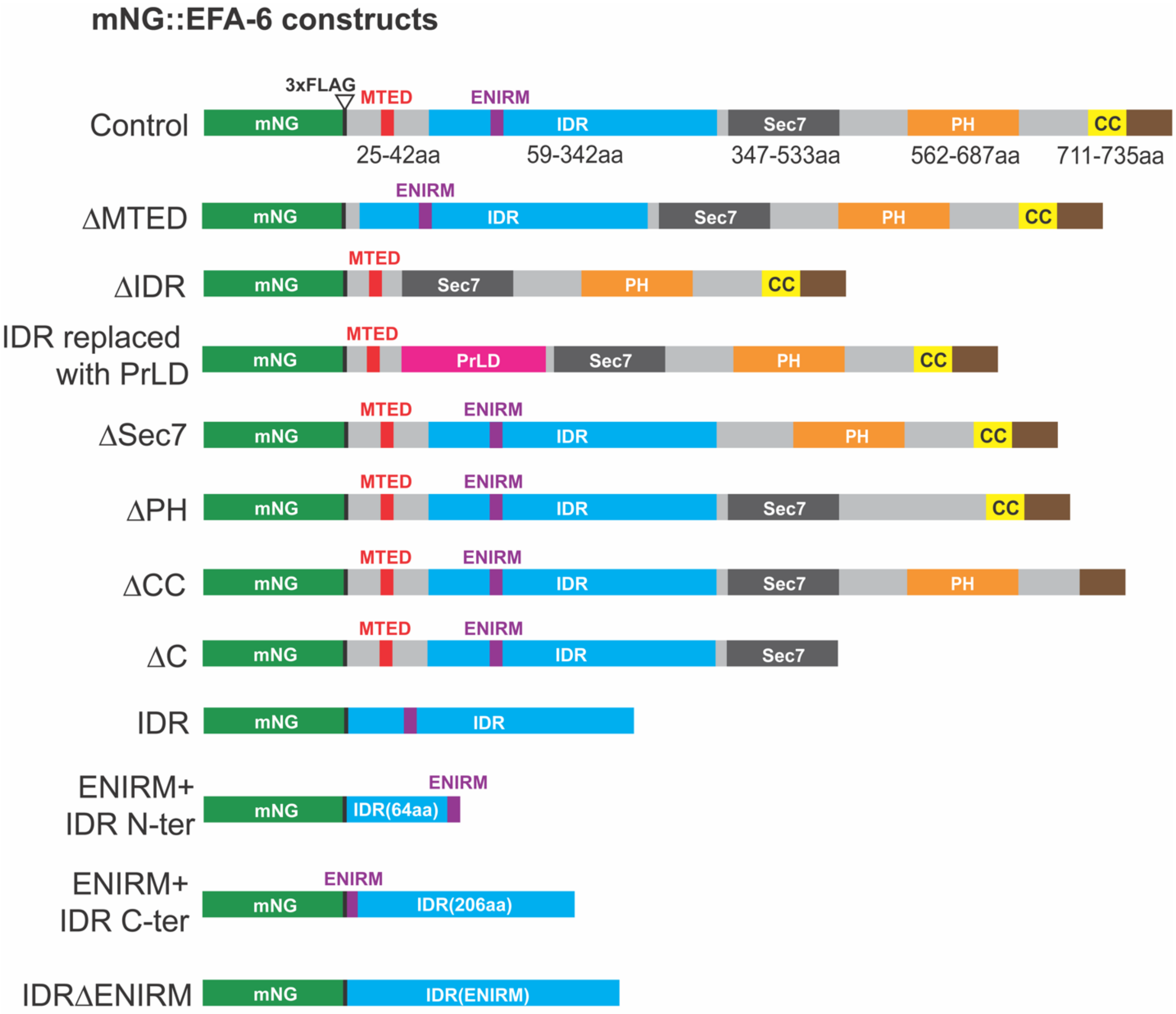


**Supplementary Figure 12. Schematic representation of EFA-6 domain deletion constructs.**

Schematic representations of N-terminally mNG tagged full-length EFA-6 (control) and the indicated domain deletion or modification constructs. Colored boxes indicate the respective domains, and gray regions represent intervening sequences. Amino acid positions of the indicated domains are shown below the full-length construct. EFA-6 contains a microtubule elimination domain (MTED), an intrinsically disordered region (IDR), a Sec7 domain, a pleckstrin homology (PH) domain, and a coiled-coil (CC) domain. All constructs are driven by epithelial-specific *col-10* promoter. All transgenes were expressed as single-copy insertions using cxTi10882 site on Chr IV in *efa-6(ju1883)* null background.

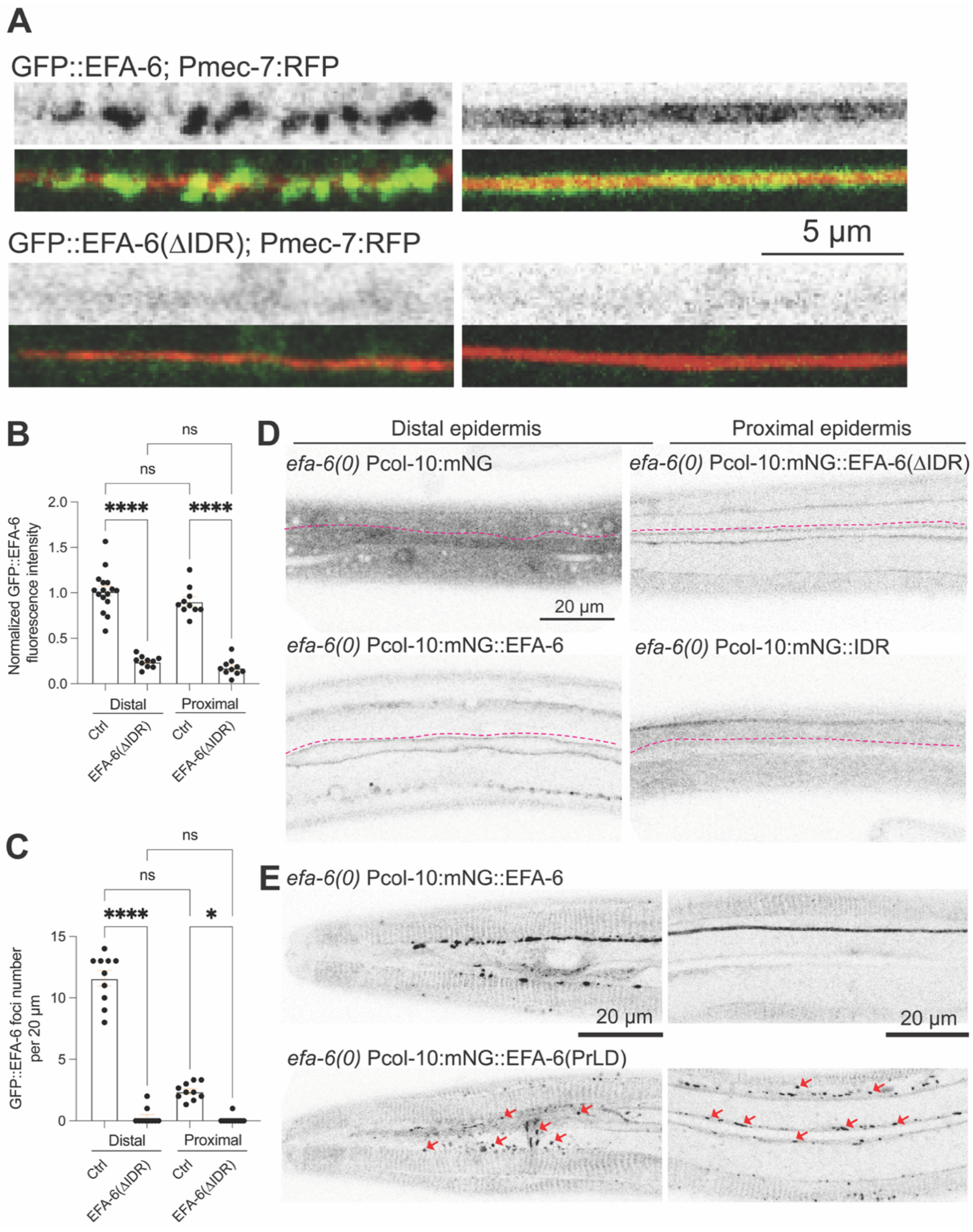

A
GFP::EFA-6; Pmec-7:RFP
GFP::EFA-6(ΔIDR); Pmec-7:RFP
5 μm
B
ns
ns
****
****
Normalized GFP::EFA-6 fluorescence intensity
2.0
1.5
1.0
0.5
0.0
Distal
Proximal
Ctrl
EFA-6(ΔIDR)
Ctrl
EFA-6(ΔIDR)
C
ns
ns
****
*
GFP::EFA-6 foci number per 20 μm
15
10
5
0
Distal
Proximal
Ctrl
EFA-6(ΔIDR)
Ctrl
EFA-6(ΔIDR)
D
Distal epidermis
Proximal epidermis
efa-6(0) Pcol-10:mNG
efa-6(0) Pcol-10:mNG::EFA-6(ΔIDR)
20 μm
efa-6(0) Pcol-10:mNG::EFA-6
efa-6(0) Pcol-10:mNG::IDR
E
efa-6(0) Pcol-10:mNG::EFA-6
20 μm
20 μm
efa-6(0) Pcol-10:mNG::EFA-6(PrLD)

**Supplementary Figure 13. EFA-6 IDR is specifically target EFA-6 to the neuron-epidermal interface, but not other cell-cell boundaries.**

**(A)** Representative single-plane confocal images of L4-stage animals expressing endogenous GFP::EFA-6 knock-in or GFP::EFA-6 lacking the IDR. Images show the distal and proximal ROIs (same as in Fig.1A) of the ALM neurite, labeled with Pmec-7:RFP. The upper panels show the GFP channel alone, whereas the lower panels show merged images. Deletion of the IDR reduces endogenous GFP::EFA-6 expression levels as well as foci formation along both distal and proximal ALM regions. n=10-15 per genotype. Scale bar, 5 μm.

**(B)** Quantification of normalized GFP::EFA-6 fluorescence intensity at the distal and proximal ROIs (same as in Fig.1A) of the ALM neurite in L4 animals expressing endogenous GFP::EFA-6 or GFP::EFA-6(ΔIDR).

**(C)** Quantification of GFP::EFA-6 foci density at the distal and proximal ROIs (same as in Fig.1A) of the ALM neurite in L4 animals endogenous expressing GFP::EFA-6 or GFP::EFA-6(ΔIDR).

**(B-C)** n=10-15 animals per genotype. Data are presented as mean ± SEM. Statistical significance was assessed using the Kruskal-Wallis test followed by Dunn's multiple-comparisons test for (B-C). ns, not significant ($P > 0.05$); *$P < 0.05$; ****$P < 0.0001$.

**(D)** Representative single-plane confocal images of L4-stage animals expressing epithelial-specific constructs: free mNG, full-length mNG::EFA-6, full-length mNG::EFA-6 lacking the IDR (mNG::EFA-6(ΔIDR)), and the isolated EFA-6 IDR tagged with mNG (mNG::IDR). All constructs are expressed as single-copy transgenes, driven by the *col-10* promoter, in the *efa-6* null background. Images show the distribution of mNG-tagged proteins in epidermis and seam cells, with seam cell dorsal boundaries outlined by magenta dotted lines. Free mNG is diffusely distributed throughout the epidermal cytoplasm without specific enrichment at the seam cell cortex. In contrast, full-length mNG::EFA-6 is enriched at the cellular cortex, prominently labeling seam cell boundaries. Deletion of the IDR does not abolish EFA-6 recruitment to seam cell membranes, indicating that the IDR is dispensable for this aspect of cortical localization. In contrast, the isolated mNG::IDR is distributed throughout the epidermis without detectable enrichment at seam cell boundaries. n=10-15 per genotype. Scale bar, 20 μm.

**(E)** Representative single-plane confocal images of L4-stage animals expressing epidermis-specific constructs: full-length mNG::EFA-6 and mNG::EFA-6(PrLD), in which the native intrinsically disordered region (IDR) of EFA-6 is replaced with the prion-like domain (PrLD) of TIAR-2. All constructs are expressed as single-copy transgenes, driven by the *col-10* promoter, in the *efa-6(ju1883)* null background. Images show the epidermis in the head and midbody regions. Full-length mNG::EFA-6 exhibits characteristic membrane localization and spatial patterning, whereas mNG::EFA-6(PrLD) forms ectopic foci within the epidermis (red arrows).

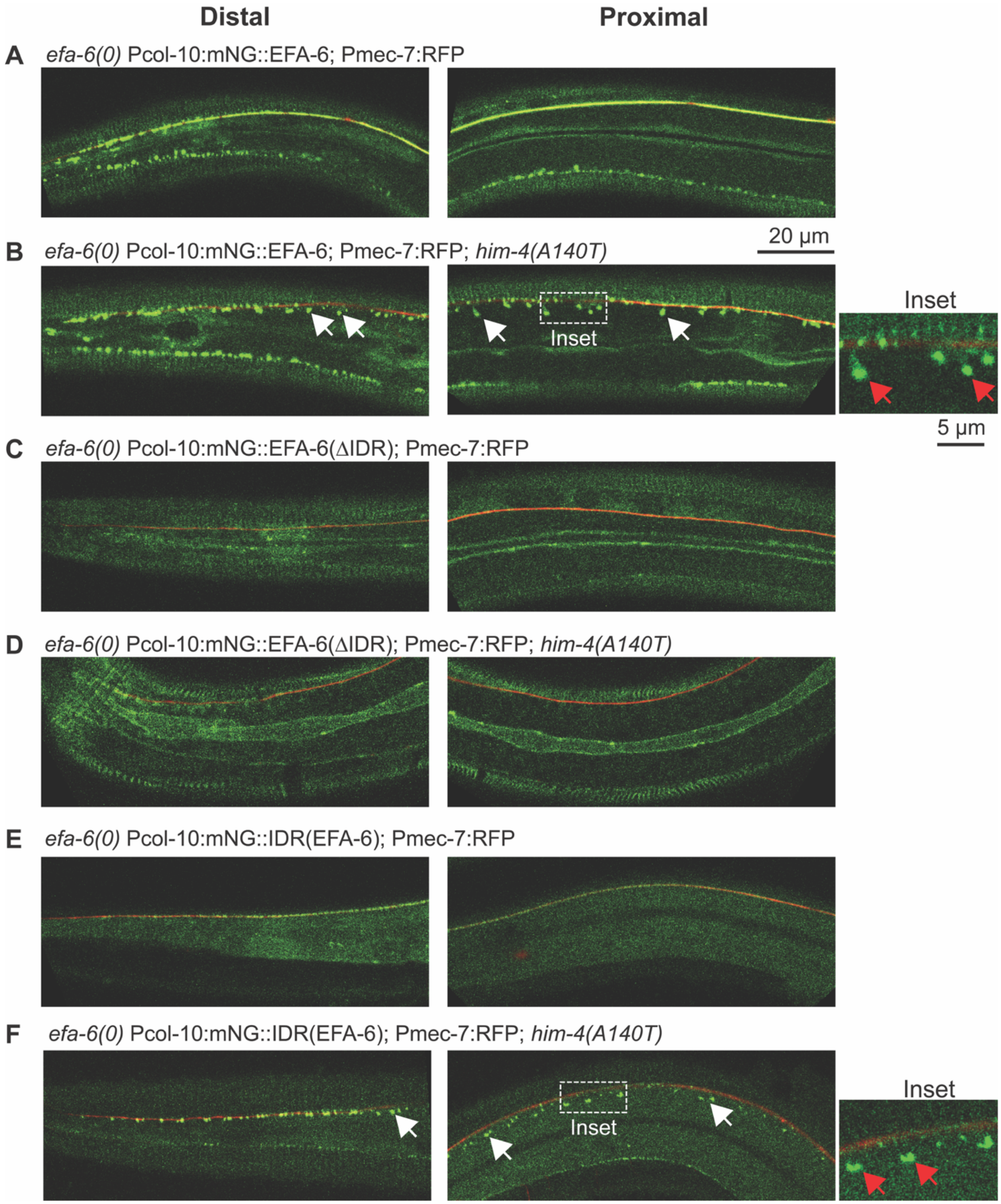
Distal
Proximal
A efa-6(0) Pcol-10:mNG::EFA-6; Pmec-7:RFP
20 µm
B efa-6(0) Pcol-10:mNG::EFA-6; Pmec-7:RFP; him-4(A140T)
Inset
Inset
5 µm
C efa-6(0) Pcol-10:mNG::EFA-6(ΔIDR); Pmec-7:RFP
D efa-6(0) Pcol-10:mNG::EFA-6(ΔIDR); Pmec-7:RFP; him-4(A140T)
E efa-6(0) Pcol-10:mNG::IDR(EFA-6); Pmec-7:RFP
F efa-6(0) Pcol-10:mNG::IDR(EFA-6); Pmec-7:RFP; him-4(A140T)
Inset
Inset

**Supplementary Figure 14: The EFA-6 IDR mediates ECM-dependent spatial patterning at the neuron-epidermal interface.**

**(A-F)** Representative single-plane confocal images of L4-stage animals expressing epithelial-specific, single-copy transgenes driven by the Pcol-10 promoter in the *efa-6(ju1883)* null background, either with wild-type or *him-4(A140T)* mutation. The constructs include full-length mNG::EFA-6 (A, B), mNG::EFA-6 lacking the intrinsically disordered region (mNG::EFA-6(ΔIDR)) (C, D), and EFA-6 IDR alone tagged with mNG (mNG::IDR) (E, F). Images show wider views of the epidermis surrounding the distal and proximal ROIs of the ALM neurite shown in Fig.1A, labeled with Pmec-7:RFP. In wild-type, full-length mNG::EFA-6 (A) is enriched along the ALM neurite and exhibits distinct spatial patterning in the distal and proximal regions. In *him-4(A140T)* mutants (B), full-length mNG::EFA-6 forms ectopic foci along the entire neurite, some of which are displaced away from the neurite while others remain closely associated with it. Deletion of the IDR (C, D) abolishes detectable neurite-associated EFA-6 fluorescence in both genetic backgrounds. In contrast, the isolated mNG::IDR (E) is sufficient for enrichment along the ALM neurite in wild-type, exhibiting both diffuse and punctate distributions. In *him-4(A140T)* mutants (F), mNG::IDR forms distinct ectopic foci, most of which are displaced away from the ALM neurite. White arrows indicate ectopic foci located away from the neurite. Insets highlight EFA-6 foci displaced from the ALM neurite (red arrows).

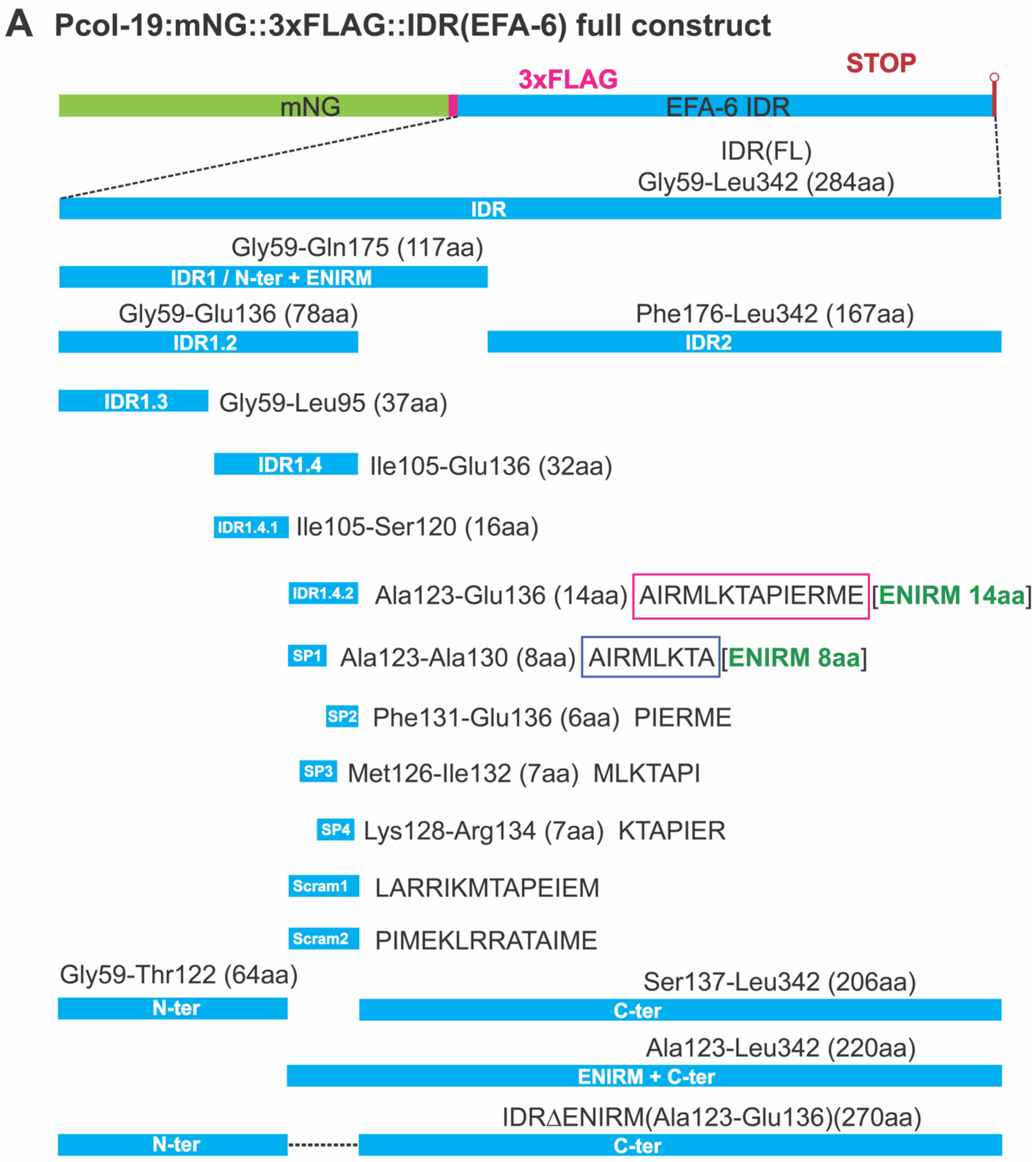


**B**

**IDR sequence-**

GTTVASSIHPHQLSRATASAGVSFPSMNRNGAAAQKLSRLPVSTSQIERRGSLARKTSEESS
PT**AIRMLKTA**PIERMESTDVEESEEETVMMTTDEKENQKKPNENDDEVMVVDEEQFIVVSND
MKSPNEEIVAKSLRSAMFTMPTDNHHHSYNSSPQISTLSPHLRSNGDGPSRSPVYDDVDDD
LNGSLDAKDMSNNSHQQSFRSPENYSEKDTPSKHSVVTIDGSGVSNHYDQDGMFSHVYYS
TQDTTPKHGSPSLRKQIFESRTTPNTAASNSSASASPSL

**Net side chain charge-**

IDR N-terminus sequence flanking **ENIRM** (marked by magenta line): +5.08
IDR C-terminus sequence flanking **ENIRM** (marked by yellow line): -20.66

**Supplementary Figure 15. Systematic truncation and mutation of the EFA-6 IDR.**

**(A)** Schematic representation of the full-length EFA-6 IDR (Gly59-Leu342, 284 aa) and the series of truncated fragments, overlapping short peptides, scrambled-sequence variants, and internal deletion constructs generated to identify sequences mediating neuron-epidermal interface targeting. The full-length IDR was divided into IDR1 (Gly59-Gln175, 117 aa) and IDR2 (Phe176-Leu342, 167 aa). Progressive truncation of IDR1 generated IDR1.2 (Gly59-Glu136, 78 aa), IDR1.3 (Gly59-Leu95, 37 aa), IDR1.4 (Ile105-Glu136, 32 aa), IDR1.4.1 (Ile105-Ser120, 16 aa), and IDR1.4.2 (Ala123-Glu136, 14 aa; referred in the study as ENIRM 14aa). Further dissection of IDR1.4.2 (AIRMLKTAPIERME) generated four overlapping short peptides: SP1 (AIRMLKTA, 8 aa; referred in the study as ENIRM 8aa), SP2 (PIERME, 6 aa), SP3 (MLKTAPI, 7 aa), and SP4 (KTAPIER, 7 aa), along with two scrambled-sequence variants (Scram1 and Scram2). Additional constructs include the N-terminal IDR fragment (Gly59-Thr122, 64 aa), the C-terminal IDR fragment (Ser137-Leu342, 206 aa), the C-terminal fragment containing the 14-amino-acid targeting sequence (Ala123-Leu342, 220 aa), and the full-length IDR lacking the 14-amino-acid IDR1.4.2 sequence (IDRΔENIRM, 270 aa). All IDR fragments and sequence variants were fused to mNG for visualization. Amino acid boundaries, fragment lengths, and short peptide sequences are indicated. ENIRM: Epidermal-Neuronal Interface Recognition Motif (ENIRM) sequence is highlighted by dark blue box for ENIRM 8aa and pink box for ENIRM 14aa.

**(B)** EFA-6 intrinsically disordered region (IDR) sequence showing positively charged amino acids highlighted in green and negatively charged amino acids in red. The eight-amino-acid ENIRM motif (A123-E130) is highlighted in blue. The N- and C-terminal sequences flanking the ENIRM motif are underlined in magenta and yellow, respectively. The net side-chain charge of each flanking sequence is indicated below the sequence.

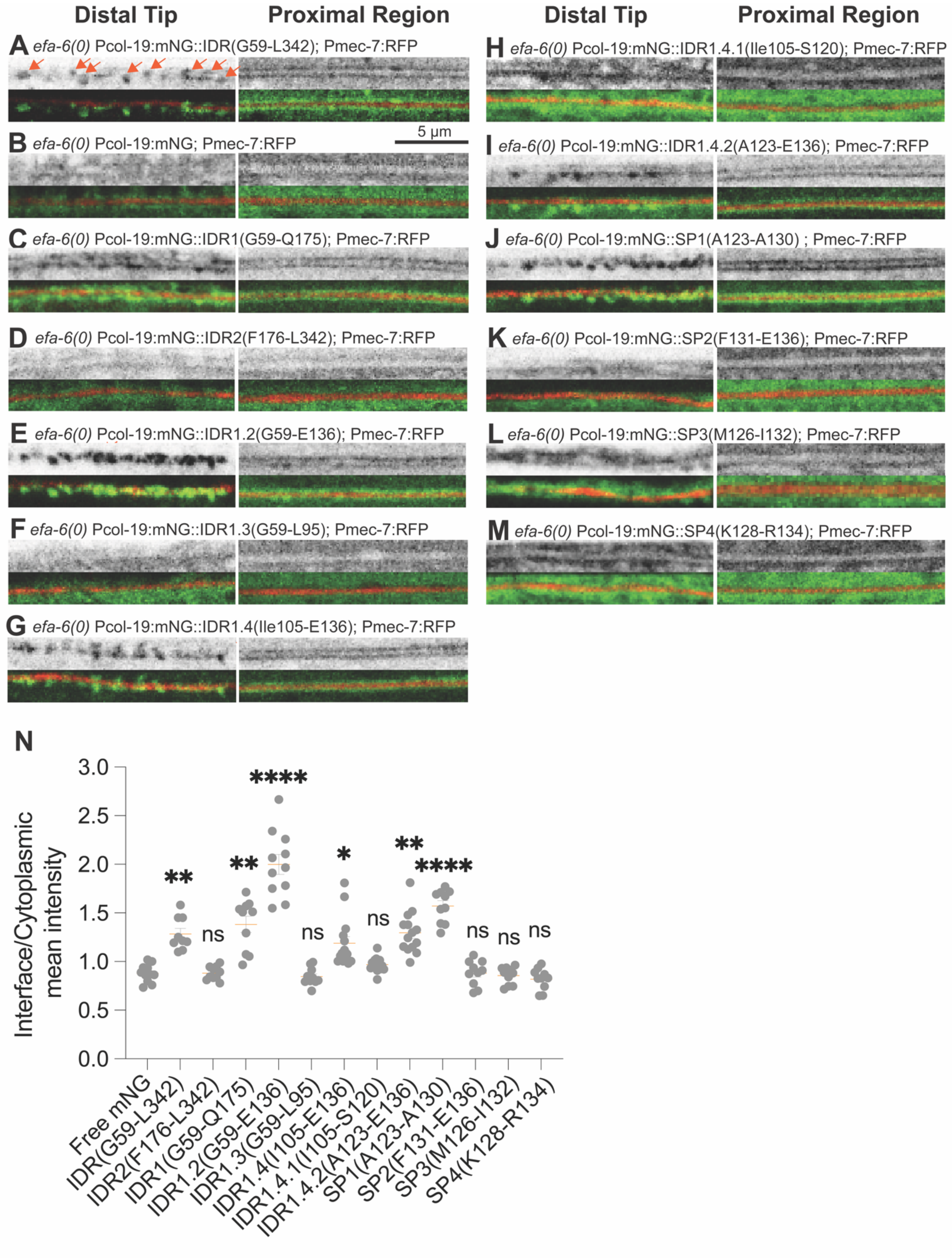

Distal Tip
Proximal Region
A efa-6(0) Pcol-19:mNG::IDR(G59-L342); Pmec-7:RFP
B efa-6(0) Pcol-19:mNG; Pmec-7:RFP
5 μm
C efa-6(0) Pcol-19:mNG::IDR1(G59-Q175); Pmec-7:RFP
D efa-6(0) Pcol-19:mNG::IDR2(F176-L342); Pmec-7:RFP
E efa-6(0) Pcol-19:mNG::IDR1.2(G59-E136); Pmec-7:RFP
F efa-6(0) Pcol-19:mNG::IDR1.3(G59-L95); Pmec-7:RFP
G efa-6(0) Pcol-19:mNG::IDR1.4(Ile105-E136); Pmec-7:RFP
H efa-6(0) Pcol-19:mNG::IDR1.4.1(Ile105-S120); Pmec-7:RFP
I efa-6(0) Pcol-19:mNG::IDR1.4.2(A123-E136); Pmec-7:RFP
J efa-6(0) Pcol-19:mNG::SP1(A123-A130) ; Pmec-7:RFP
K efa-6(0) Pcol-19:mNG::SP2(F131-E136); Pmec-7:RFP
L efa-6(0) Pcol-19:mNG::SP3(M126-I132); Pmec-7:RFP
M efa-6(0) Pcol-19:mNG::SP4(K128-R134); Pmec-7:RFP
N
Interface/Cytoplasmic mean intensity
3.0
2.5
2.0
1.5
1.0
0.5
0.0
**
ns
**
****
ns
*
ns
**
****
ns
ns
ns
Free mNG
IDR(G59-L342)
IDR2(F176-L342)
IDR1(G59-Q175)
IDR1.2(G59-E136)
IDR1.3(G59-L95)
IDR1.4(I105-E136)
IDR1.4.1(I105-S120)
IDR1.4.2(A123-E136)
SP1(A123-A130)
SP2(F131-E136)
SP3(M126-I132)
SP4(K128-R134)

**Supplementary Figure 16. Systematic dissection of the EFA-6 IDR identifies sequences mediating neuron-epidermal interface targeting.**

**(A-M)** Representative single-slice confocal images of adult animals expressing epithelial-specific mNG-tagged EFA-6 IDR fragments. All constructs are expressed as single-copy transgenes, driven by the *col-10* promoter, in the *efa-6* null background. Images show the distal and proximal ROIs (same as in Fig.1A) of the ALM neurite, labeled with Pmec-7:RFP. The upper panels show the mNG channel alone, whereas the lower panels show merged mNG and RFP channels. Constructs include (A) the full-length EFA-6 IDR (Gly59-Leu342, 284 aa), (B) free mNG, (C) IDR1 (Gly59-Gln175, 117 aa), (D) IDR2 (Phe176-Leu342, 167 aa), and progressively truncated fragments of IDR1: (E) IDR1.2 (Gly59-Glu136, 78 aa), (F) IDR1.3 (Gly59-Leu95, 37 aa), (G) IDR1.4 (Ile105-Glu136, 32 aa), (H) IDR1.4.1 (Ile105-Ser120, 16 aa), and (I) IDR1.4.2 (Ala123-Glu136, ENIRM 14 aa). Further dissection of the 14-amino-acid fragment (AIRMLKTAPIERME) generated (J-M) four overlapping short peptides: SP1 (AIRMLKTA, 8 aa; referred in the study as ENIRM 8aa), SP2 (PIERME, 6 aa), SP3 (MLKTAPI, 7 aa), and SP4 (KTAPIER, 7 aa) to identify minimal sequences sufficient for interface targeting. IDR1, IDR1.2, IDR1.4, and IDR1.4.2 recapitulate the spatial enrichment pattern of the full-length IDR along the ALM neurite, whereas the remaining IDR fragments fail to reproduce this pattern. Among the short peptides, SP1 (AIRMLKTA; ENIRM 8 aa) retains neuron-epidermal interface enrichment comparable to that of the full-length IDR, whereas SP2-SP4 do not exhibit detectable enrichment. Orange arrows mark EFA-6 foci. n=10-12 animals per genotype. Scale bar, 5 μm.

**(N)** Quantification of fluorescence enrichment in the proximal ROIs (same as in Fig.1A) of the ALM neurite in adult animals expressing the epithelial-specific constructs shown in panel (A-M). n=10-15 animals per genotype. Data are presented as mean ± SEM. Statistical significance was assessed using the Kruskal-Wallis test followed by Dunn's multiple-comparisons test. ns, not significant ($P > 0.05$); $^{*}P < 0.05$; $^{**}P < 0.01$; $^{****}P < 0.0001$.

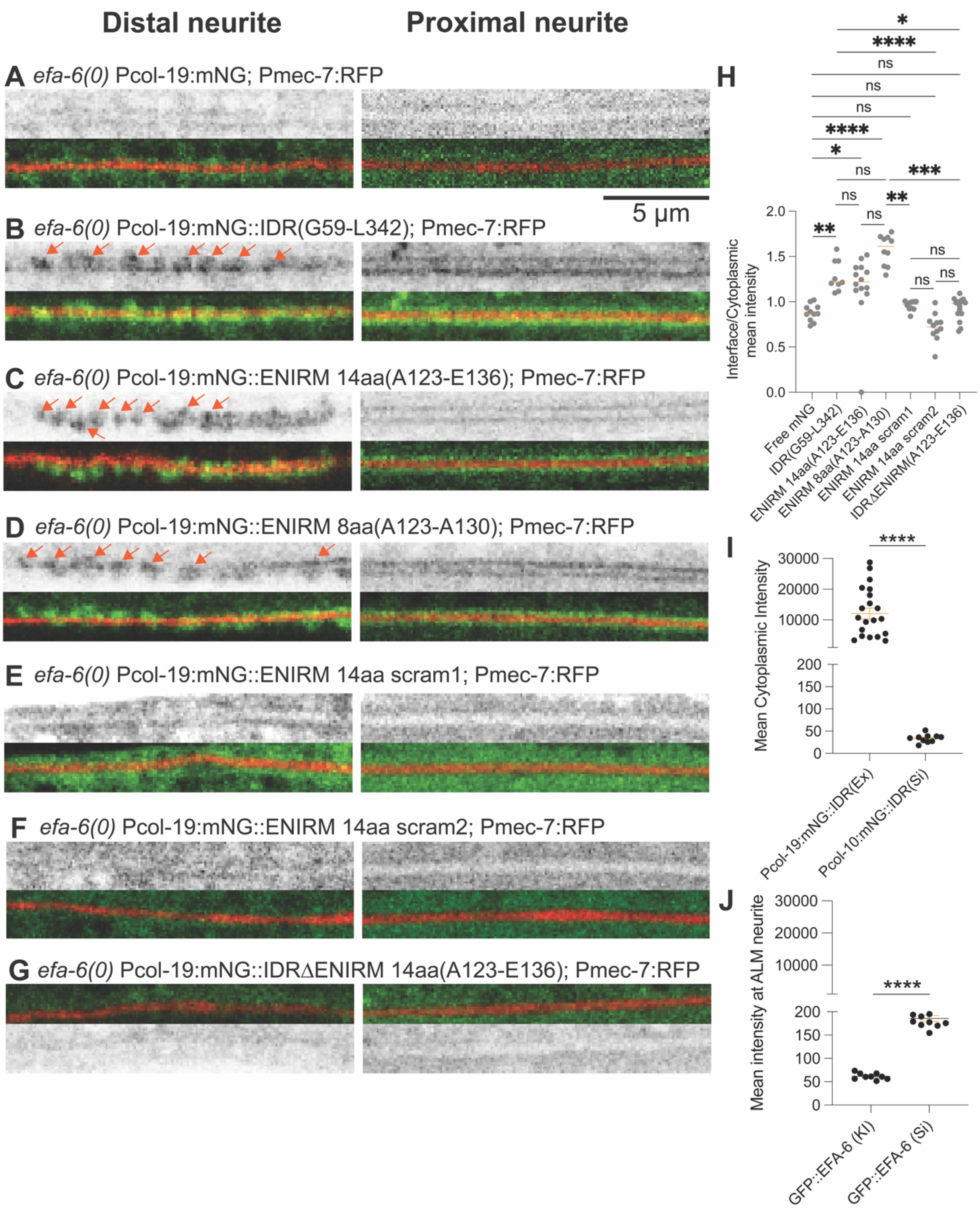

Distal neurite
Proximal neurite
A efa-6(0) Pcol-19:mNG; Pmec-7:RFP
5 µm
B efa-6(0) Pcol-19:mNG::IDR(G59-L342); Pmec-7:RFP
C efa-6(0) Pcol-19:mNG::ENIRM 14aa(A123-E136); Pmec-7:RFP
D efa-6(0) Pcol-19:mNG::ENIRM 8aa(A123-A130); Pmec-7:RFP
E efa-6(0) Pcol-19:mNG::ENIRM 14aa scram1; Pmec-7:RFP
F efa-6(0) Pcol-19:mNG::ENIRM 14aa scram2; Pmec-7:RFP
G efa-6(0) Pcol-19:mNG::IDRΔENIRM 14aa(A123-E136); Pmec-7:RFP
H
Interface/Cytoplasmic mean intensity
Free mNG
IDR(G59-L342)
ENIRM 14aa(A123-E136)
ENIRM 8aa(A123-A130)
ENIRM 14aa scram1
ENIRM 14aa scram2
IDRΔENIRM(A123-E136)
I
Mean Cytoplasmic Intensity
Pcol-19:mNG::IDR(Ex)
Pcol-10:mNG::IDR(Si)
J
Mean intensity at ALM neurite
GFP::EFA-6 (KI)
GFP::EFA-6 (Si)

**Supplementary Figure 17. ENIRM is necessary and sufficient for EFA-6 IDR targeting to the neuron-epidermal interface under overexpression conditions.**

**(A-G)** Representative single-plane confocal images of adult animals expressing epithelial-specific, mNG-tagged EFA-6 IDR constructs driven by the Pcol-19 promoter in the *efa-6(ju1883)* null background. Images show the distal and proximal regions of the ALM neurite, labeled with Pmec-7::RFP. The upper panels show the mNG channel alone, whereas the lower panels show merged mNG and RFP channels. Constructs include (A) free mNG, (B) the full-length EFA-6 IDR (G59-L342, 284 aa), (C) the 14-amino-acid ENIRM-containing fragment (IDR1.4.2; AIRMLKTAPIERME), (D) the 8-amino-acid ENIRM sequence (SP1; AIRMLKTA), (E, F) scrambled variants of the 14-amino-acid fragment, and (G) the full-length IDR lacking the 14-amino-acid ENIRM 14aa sequence (IDRΔENIRM 14aa(A123-E136)). Please see Fig. S15 for construct details. Both the 14-amino-acid and 8-amino-acid fragments recapitulate the spatial enrichment pattern of the full-length IDR along the distal and proximal regions of the ALM neurite. In contrast, scrambling the 14-amino-acid sequence or deleting 14aa ENIRM sequence from the full-length IDR disrupts neuron-epidermal interface enrichment. Orange arrows mark EFA-6 foci. n=10 animals per genotype. Scale bar, 5 μm.

**(H)** Quantification of fluorescence enrichment in the proximal ROI (same as in Fig.1A) of the ALM neurite in animals expressing the epithelial-specific constructs shown in (A-G).

**(I)** Quantification of mean cytoplasmic fluorescence intensity in the epidermis, within the proximal ROI as shown in Fig.1A, in *efa-6* null animals expressing epidermis-specific mNG::IDR driven by the *col-19* promoter from extrachromosomal arrays or by the *col-10* promoter as a single-copy insertion transgene.

**(J)** Quantification of mean fluorescence intensity at the proximal ROI (same as in Fig.1A) of the ALM neurite of animals expressing endogenous GFP::EFA-6 knock-in or epidermis-specific mNG::EFA-6 from a *col-10*-driven single-copy insertion transgene in *efa-6* null animals.

**(H-J)** n=10-15 animals per genotype. Data are presented as mean ± SEM. Statistical significance was assessed using the Kruskal-Wallis test followed by Dunn's multiple-comparisons test for (H) and unpaired *t*-test for (I, J). ns, not significant ($P > 0.05$); *$P < 0.05$; **$P < 0.01$; ***$P < 0.001$; ****$P < 0.0001$.

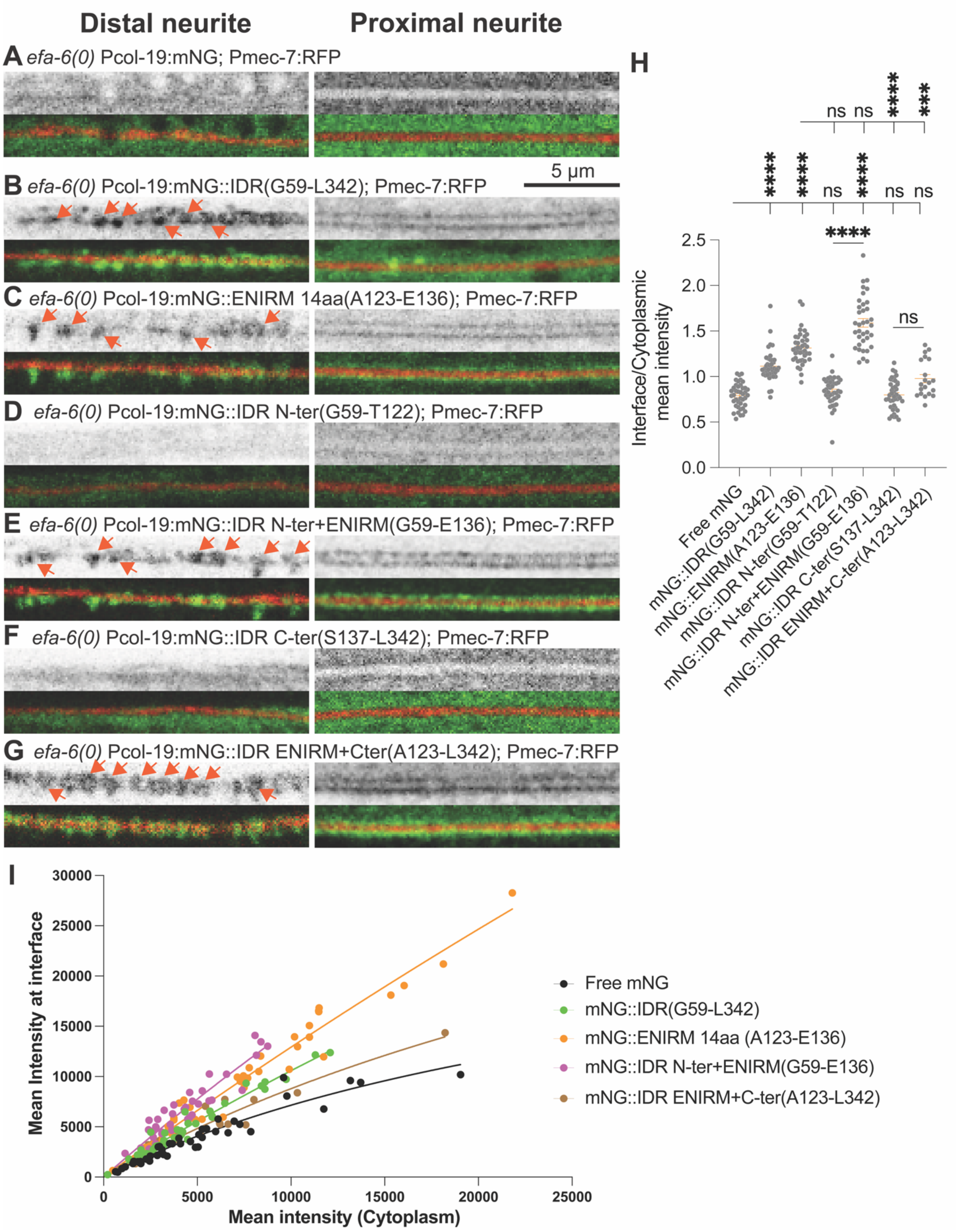

Distal neurite
Proximal neurite
A efa-6(0) Pcol-19:mNG; Pmec-7:RFP
5 μm
B efa-6(0) Pcol-19:mNG::IDR(G59-L342); Pmec-7:RFP
C efa-6(0) Pcol-19:mNG::ENIRM 14aa(A123-E136); Pmec-7:RFP
D efa-6(0) Pcol-19:mNG::IDR N-ter(G59-T122); Pmec-7:RFP
E efa-6(0) Pcol-19:mNG::IDR N-ter+ENIRM(G59-E136); Pmec-7:RFP
F efa-6(0) Pcol-19:mNG::IDR C-ter(S137-L342); Pmec-7:RFP
G efa-6(0) Pcol-19:mNG::IDR ENIRM+Cter(A123-L342); Pmec-7:RFP
H
Interface/Cytoplasmic mean intensity
Free mNG
mNG::IDR(G59-L342)
mNG::ENIRM(A123-E136)
mNG::IDR N-ter(G59-T122)
mNG::IDR N-ter+ENIRM(G59-E136)
mNG::IDR C-ter(S137-L342)
mNG::IDR ENIRM+C-ter(A123-L342)
I
Mean Intensity at interface
Mean intensity (Cytoplasm)
Free mNG
mNG::IDR(G59-L342)
mNG::ENIRM 14aa (A123-E136)
mNG::IDR N-ter+ENIRM(G59-E136)
mNG::IDR ENIRM+C-ter(A123-L342)

**Supplementary Figure 18. C-terminal flanking disordered sequences attenuate ENIRM-mediated targeting to the neuron-epidermal interface under overexpression conditions.**

**(A-G)** Representative single-slice confocal images of adult animals overexpressing epithelial-specific, mNG-tagged EFA-6 IDR constructs driven by the Pcol-19 promoter in the *efa-6(ju1883)* null background. Constructs include **(A)** free mNG, **(B)** the full-length EFA-6 IDR tagged with mNG (mNG::IDR(G59-L342)), **(C)** the isolated 14-amino-acid ENIRM-containing fragment (mNG::ENIRM14aa (A123-E136)), **(D)** the N-terminal disordered sequence flanking ENIRM (mNG::IDR(G59-T122)), **(E)** the 14-amino-acid ENIRM-containing fragment combined with its N-terminal flanking disordered sequence (mNG::IDR(G59-E136)), **(F)** the C-terminal disordered sequence flanking ENIRM (mNG::IDR(S137-L342)), and **(G)** the 14-amino-acid ENIRM-containing fragment combined with its C-terminal flanking disordered sequence (mNG::IDR(A123-L342)). Please see Fig. S15 for construct details. Images show the distal and proximal ROIs (same as in Fig.1A) of the ALM neurite, labeled with Pmec-7:RFP. The upper panels show the mNG channel alone, whereas the lower panels show merged images. The full-length IDR, isolated 14-amino-acid ENIRM-containing fragment, and ENIRM-containing fragments with either N-terminal or C-terminal flanking disordered sequences exhibit enrichment along the ALM neurite. In contrast, neither the N-terminal nor the C-terminal flanking disordered sequence alone shows detectable enrichment at the neuron-epidermal interface. Notably, the isolated ENIRM-containing fragment and its N-terminal extension exhibit stronger interface enrichment than the full-length IDR or the ENIRM-containing fragment with its C-terminal extension, suggesting that C-terminal flanking disordered sequences attenuate ENIRM-mediated interface targeting under overexpression conditions. Orange arrows mark EFA-6 foci. n=40-45 animals per genotype. Scale bar, 5 μm.

**(H)** Quantification of fluorescence enrichment in the proximal ROI (same as in Fig.1A) of the ALM neurite in animals expressing the epithelial-specific constructs shown in (A-G).

**(I)** Relationship between mean fluorescence intensity at the neuron-epidermal interface and mean cytoplasmic fluorescence intensity in the proximal region of the ALM neurite for the epithelial-specific constructs shown in (A-G). Each data point represents measurements from an individual animal.

**(H, I)** n=20-40 animals per genotype for (H); n=40-45 animals per genotype for (I). Data are presented as mean ± SEM. Statistics: Kruskal-Wallis test followed by Dunn's multiple-comparisons test for (H). ns, not significant ($P > 0.05$); *$P < 0.05$; **$P < 0.01$; ***$P < 0.001$; ****$P < 0.0001$.
