## Supplemental Tables for "A specialized extracellular matrix patterns the conserved cortical protein EFA-6 at neuron-epidermal interfaces"

**Sandhu, Lyu, et al. Supplemental Tables 1-7**

**Table S1:** Strains used in the study

| Strain number | Allele | Reference |
|---|---|---|
| CZ26544 | P*mec-7*-mRFP + CB*unc-119* (*jsIs973*); *gfp::efa-6(ju1658)* | (Sandhu et al., 2024) |
| CZ29798 | P*mec-7*-mRFP + CB*unc-119* (*jsIs973*); *efa-6(ju1990ju1658)* | (Sandhu et al., 2024) |
| CZ30221 | P*mec-7*-mRFP + CB*unc-119* (*jsIs973*); *efa-6(ju1833ju1658)* | (Sandhu et al., 2024) |
| CZ10175 | Pmec-4:GFP(zdls5) | (Chen et al., 2011; Clark & Chiu, 2003) |
| CZ31574 | P*mec-7*-mRFP + CB*unc-119* (*jsIs973*); *efa-6(ju1658ju2193)* | (Sandhu et al., 2024) |
| CZ31871 | Prgef-1-TIR1(reSi7[rgef-1p::TIR1::F2A::mTagBFP2:: AID*::NLS::tbb-2 3'UTR]); Pmec-7-mRFP + CBunc-119(jsIs973); gfp::efa-6(ju1658ju2193) | This study |
| CZ31575 | Peft-3-TIR1(wrdSi22[eft-3p::TIR1:F2A:mTagBFP::tbb2 3' UTR]); Pmec-7-mRFP + CBunc-119 (jsIs973); gfp::efa-6(ju1658ju2193) | This study |
| CZ31872 | Pcol-10-TIR1(reSi2[col-10p::TIR1::F2A::mTagBFP2:: AID*::NLS::tbb-2 3'UTR]); Pmec-7-mRFP + CBunc-119(jsIs973); gfp::efa-6(ju1658ju2193) | This study |
| CZ27176 | *gfp::efa-6(ju1658); mec-1(ju1688)* | This study |
| CZ27177 | *gfp::efa-6(ju1658); mec-1(ju1688)* | This study |
| CZ27336 | P*mec-7*-mRFP + CB*unc-119* (*jsIs973*); *gfp::efa-6(ju1658); mec-1(ju1688)* | This study |
| CZ27696 | P*mec-7*-mRFP + CB*unc-119* (*jsIs973*); *gfp::efa-6(ju1658); mec-1(e1066)* | This study |
| CZ27174 | *gfp::efa-6(ju1658); him-4(ju1687)* | This study |
| CZ27175 | *gfp::efa-6(ju1658); him-4(ju1687)* | This study |
| CZ28616 | P*mec-7*-mRFP + CB*unc-119* (*jsIs973*); *gfp::efa-6(ju1658); him-4(ju1687)* | This study |
| CZ27332 | *gfp::efa-6(ju1658); him-4(ju1768)* | This study |
| CZ27463 | *gfp::efa-6(ju1658); him-4(ju1768)* | This study |
| CZ27586 | P*mec-7*-mRFP + CB*unc-119* (*jsIs973*); *gfp::efa-6(ju1658); him-4(ju1768)* | This study |
| CZ27178 | *gfp::efa-6(ju1658); him-4(ju1689)* | This study |
| CZ27179 | *gfp::efa-6(ju1658); him-4(ju1689)* | This study |
| CZ27348 | P*mec-7*-mRFP + CB*unc-119* (*jsIs973*); *gfp::efa-6(ju1658); him-4(ju1689)* | This study |
| CZ31445 | P*mec-7*-mRFP + CB*unc-119* (*jsIs973*); *gfp::efa-6(ju1658); him-4(ju2231)* | This study |
| CZ31701 | P*mec-7*-mRFP + CB*unc-119* (*jsIs973*); *efa-6(ju1990ju1658); him-4(ju2231)* | This study |
| CZ29382 | P*mec-7*-mRFP + CB*unc-119* (*jsIs973*); *gfp::efa-6(ju1658); him-4(rh319)* | This study |
| CZ32262 | mScarlet::EFA-6(*syb6998*); him-4::mNG(qy33); Pmec-4:BFP + Pttx-3:RFP (juEx8629) | This study |
| CZ32187 | mScarlet::EFA-6(*syb6998*); him-4(qy33ju2232); Pmec-4::BFP + Pttx-3 RFP(juEx8629) | This study |
| CZ32188 | mScarlet::EFA-6(*syb6998*); him-4(qy33ju2233); Pmec-4::BFP + Pttx-3 RFP(juEx8629) | This study |
| CZ32189 | mScarlet::EFA-6(*syb6998*); him-4(qy33ju2235); Pmec-4::BFP + Pttx-3 RFP(juEx8629) | This study |
| CZ31908 | Pmec-7-mRFP + CBunc-119(jsIs973); efa-6(ju1883ju1658) Pcol-10:mNG::3xFLAG(juSi415) | This study |

| | | |
|---|---|---|
| CZ31874 | Pmec-7-mRFP + CBunc-119(jsIs973); efa-6(ju1883ju1658) Pcol-10:mNG::3xFLAG::EFA-6(juSi408) | This study |
| CZ32488 | Pmec-7-mRFP + CBunc-119(jsIs973); efa-6(ju1883ju1658) Pcol-10:mNG::3xFLAG::EFA-6ΔPH(juSi746) | This study |
| CZ32222 | Pmec-7-mRFP + CBunc-119(jsIs973); efa-6(ju1883ju1658) Pcol-10:mNG::3xFLAG::EFA-6(ΔCC)(juSi422) | This study |
| CZ31911 | Pmec-7-mRFP + CBunc-119(jsIs973); efa-6(ju1883ju1658) Pcol-10:mNG::3xFLAG::EFA-6(ΔC) (juSi417) | This study |
| CZ32221 | Pmec-7-mRFP + CBunc-119(jsIs973); efa-6(ju1883ju1658) Pcol-10:mNG::3xFLAG::EFA-6(Δsec7) (juSi421) | This study |
| CZ31909 | Pmec-7-mRFP + CBunc-119(jsIs973); efa-6(ju1883ju1658) Pcol-10:mNG::3xFLAG::EFA-6(ΔMTED)(juSi416) | This study |
| CZ31910 | Pmec-7-mRFP + CBunc-119(jsIs973); efa-6(ju1883ju1658) Pcol-10:mNG::3xFLAG::EFA-6(ΔIDR)(juSi414) | This study |
| CZ31870 | Pmec-7-mRFP + CBunc-119(jsIs973); efa-6(ju1883ju1658) Pcol-10:mNG::3xFLAG::EFA-6(TIAR2 PrLD) (juSi411) | This study |
| CZ32223 | Pmec-7-mRFP + CBunc-119(jsIs973); efa-6(ju1883ju1658) Pcol-10:mNG::3xFLAG::IDR(G59-L342) (juSi423) | This study |
| CZ32490 | Pmec-7-mRFP + CBunc-119(jsIs973); efa-6(ju1883ju1658) Pcol-10:mNG::3xFLAG::IDRΔENIRM(A123-E136) (juSi752) | This study |
| CZ32487 | Pmec-7-mRFP + CBunc-119(jsIs973); efa-6(ju1883ju1658) Pcol-10:mNG::3xFLAG::ENIRM(A123-E136) (juSi749) | This study |
| CZ32484 | Pmec-7-mRFP + CBunc-119(jsIs973); efa-6(ju1883ju1658) Pcol-10:mNG::3xFLAG::IDR(G59-E136)(juSi750) | This study |
| CZ32491 | Pmec-7-mRFP + CBunc-119(jsIs973); efa-6(ju1883ju1658) Pcol-10:mNG::3xFLAG::IDR(A123-L342) (juSi751) | This study |
| NM3336 | Pmec-7-mRFP + CBunc-119(jsIs973) | (Zheng et al., 2011) |
| CZ32228 | Pmec-7-mRFP + CBunc-119(jsIs973); efa-6(ju1883ju1658); him-4(ju2231) | This study |
| CZ32224 | Pmec-7-mRFP + CBunc-119(jsIs973); efa-6(ju1883ju1658); Pcol-10:mNG::3xFLAG::EFA-6(juSi408); him-4(ju2231) | This study |
| CZ32226 | Pmec-7-mRFP + CBunc-119(jsIs973); efa-6(ju1883ju1658); Pcol-10:mNG::3xFLAG::EFA-6(ΔIDR)(juSi414); him-4(ju2231) | This study |
| CZ32234 | Pmec-7-mRFP + CBunc-119(jsIs973); *gfp::efa-6*(ju2193ju1658); him-4(rh319) | This study |
| CZ32236 | Peft-3-TIR1(wrdSi22[eft-3p::TIR1:F2A:mTagBFP::tbb2 3' UTR]); Pmec-7-mRFP + CBunc-119(jsIs973); *gfp::efa-6*(ju2193ju1658); him-4(rh319) | This study |
| CZ32235 | Prgef-1-TIR1(reSi7[rgef-1p::TIR1::F2A::mTagBFP2::AID*::NLS::tbb-2 3'UTR]); Pmec-7-mRFP + CBunc-119(jsIs973); *gfp::efa-6*(ju2193ju1658); him-4(rh319) | This study |
| CZ32237 | Pcol-10-TIR1(reSi2[col-10p:TIR1::F2A::mTagBFP2::AID*::NLS::tbb-2 3'UTR]); Pmec-7-mRFP + CBunc-119(jsIs973); *gfp::efa-6*(ju2193ju1658); him-4(rh319) | This study |
| CZ32467 | Pmec-7-mRFP + CBunc-119(jsIs973); *him-4::mNG(qy33)* | This study |
| CZ32468 | Pmec-7-mRFP + CBunc-119(jsIs973); *efa-6(ju1833ju1658); him-4::mNG(qy33)* | This study |
| CZ32469 | Pmec-7-mRFP + CBunc-119(jsIs973); *him-4::mNG(qy33ju2232)* | This study |
| CZ32470 | Pmec-7-mRFP + CBunc-119(jsIs973); *efa-6(ju1833ju1658); him-4::mNG(qy33ju2232)* | This study |
| CZ32479 | Pmec-7-mRFP + CBunc-119(jsIs973); *him-4::mNG(qy33ju2233)* | This study |
| CZ32480 | Pmec-7-mRFP + CBunc-119(jsIs973); *efa-6(ju1833ju1658); him-4::mNG(qy33ju2233)* | This study |
| CZ32481 | Pmec-7-mRFP + CBunc-119(jsIs973); *him-4::mNG(qy33ju2235)* | This study |

| | | |
|---|---|---|
| CZ32482 | Pmec-7-mRFP + CBunc-119(jsIs973); *efa-6(ju1833ju1658); him-4::mNG(qy33ju2235)* | This study |
| CZ30212 | *mSc::efa-6(syb6998); him-4::mNG(qy33)* | This study |
| CZ31618 | *mSc::efa-6(syb6998); him-4(qy33ju2232)* | This study |
| CZ31619 | *mSc::efa-6(syb6998); him-4(qy33ju2233)* | This study |
| CZ31622 | *mSc::efa-6(syb6998); him-4(qy33ju2235)* | This study |
| CZ32455 | Pmec-7-mRFP + CBunc-119(jsIs973); efa-6(ju1883ju1658); Pcol-19: mNG::3xFLAG (juEx8682) line-1 | This study |
| CZ32456 | Pmec-7-mRFP + CBunc-119(jsIs973); efa-6(ju1883ju1658); Pcol-19: mNG::3xFLAG (juEx8682) line-1 | This study |
| CZ32244 | Pmec-7-mRFP + CBunc-119(jsIs973); efa-6(ju1883ju1658); Pcol-19:mNG::3xFLAG::IDR(G59-L342) (juEx8630) | This study |
| CZ32245 | Pmec-7-mRFP + CBunc-119(jsIs973); efa-6(ju1883ju1658); Pcol-19: mNG::3xFLAG::IDR(G59-L342) (juEx8630) | This study |
| CZ32453 | Pmec-7-mRFP + CBunc-119(jsIs973); efa-6(ju1883ju1658); Pcol-19: mNG::3xFLAG::IDRΔENIRM(A123-E136) (juEx8690) line-1 | This study |
| CZ32454 | Pmec-7-mRFP + CBunc-119(jsIs973); efa-6(ju1883ju1658); Pcol-19: mNG::3xFLAG::IDRΔENIRM(A123-E136) (juEx8690) line-2 | This study |
| CZ32459 | Pmec-7-mRFP + CBunc-119(jsIs973); efa-6(ju1883ju1658); Pcol-19: mNG::3xFLAG::ENIRM 14aa(A123-E136) (juEx8685) line-1 | This study |
| CZ32460 | Pmec-7-mRFP + CBunc-119(jsIs973); efa-6(ju1883ju1658); Pcol-19: mNG::3xFLAG::ENIRM 14aa(A123-E136) (juEx8685) line-2 | This study |
| CZ32457 | Pmec-7-mRFP + CBunc-119(jsIs973); efa-6(ju1883ju1658); Pcol-19: mNG::3xFLAG::IDR(G59-T122) (juEx8693) line-1 | This study |
| CZ32458 | Pmec-7-mRFP + CBunc-119(jsIs973); efa-6(ju1883ju1658); Pcol-19: mNG::3xFLAG::IDR(G59-T122) (juEx8693) line-2 | This study |
| CZ32250 | Pmec-7-mRFP + CBunc-119(jsIs973); efa-6(ju1883ju1658); Pcol-19: mNG::3xFLAG::IDR(G59-E136) (juEx8633) line-1 | This study |
| CZ32251 | Pmec-7-mRFP + CBunc-119(jsIs973); efa-6(ju1883ju1658); Pcol-19: mNG::3xFLAG:: IDR(G59-E136) (juEx8633) line-2 | This study |
| CZ32451 | Pmec-7-mRFP + CBunc-119(jsIs973); efa-6(ju1883ju1658); Pcol-19: mNG::3xFLAG::IDR(S137-L342) (juEx8694) line-1 | This study |
| CZ32452 | Pmec-7-mRFP + CBunc-119(jsIs973); efa-6(ju1883ju1658); Pcol-19: mNG::3xFLAG::IDR(S137-L342) (juEx8694) line-2 | This study |
| CZ32447 | Pmec-7-mRFP + CBunc-119(jsIs973); efa-6(ju1883ju1658); Pcol-19: mNG::3xFLAG::IDR(A123-L342) (juEx8695) line-1 | This study |
| CZ32466 | Pmec-7-mRFP + CBunc-119(jsIs973); efa-6(ju1883ju1658); Pcol-19: mNG::3xFLAG::IDR(A123-L342) (juEx8695) line-2 | This study |
| CZ32436 | Pmec-7-mRFP + CBunc-119(jsIs973); efa-6(ju1883ju1658); Pcol-19: mNG::3xFLAG::ENIRM 14aa Scram1(juEx8691) line-1 | This study |
| CZ32437 | Pmec-7-mRFP + CBunc-119(jsIs973); efa-6(ju1883ju1658); Pcol-19: mNG::3xFLAG::ENIRM 14aa Scram1(juEx8691) line-2 | This study |
| CZ32438 | Pmec-7-mRFP + CBunc-119(jsIs973); efa-6(ju1883ju1658); Pcol-19: mNG::3xFLAG::ENIRM 14aa Scram2 (juEx8692) line-1 | This study |
| CZ32439 | Pmec-7-mRFP + CBunc-119(jsIs973); efa-6(ju1883ju1658); Pcol-19: mNG::3xFLAG::ENIRM 14aa Scram2 (juEx8692) line-2 | This study |
| CZ32246 | Pmec-7-mRFP + CBunc-119(jsIs973); efa-6(ju1883ju1658); Pcol-19: mNG::3xFLAG::IDR1(G59-Q175) (juEx8631) | This study |
| CZ32247 | Pmec-7-mRFP + CBunc-119(jsIs973); efa-6(ju1883ju1658); Pcol-19: mNG::3xFLAG::IDR1(G59-Q175) (juEx8631) | This study |
| CZ32248 | Pmec-7-mRFP + CBunc-119(jsIs973); efa-6(ju1883ju1658); Pcol-19: mNG::3xFLAG::IDR2(F176-L342) (juEx8632) | This study |
| CZ32249 | Pmec-7-mRFP + CBunc-119(jsIs973); efa-6(ju1883ju1658); Pcol-19: mNG::3xFLAG::IDR2(F176-L342) (juEx8632) | This study |
| CZ32250 | Pmec-7-mRFP + CBunc-119(jsIs973); efa-6(ju1883ju1658); Pcol-19: | This study |

| | | |
|---|---|---|
| | mNG::3xFLAG::IDR1.2(G59-E136) (juEx8633) | |
| CZ32251 | Pmec-7-mRFP + CBunc-119(jsIs973); efa-6(ju1883ju1658); Pcol-19: mNG::3xFLAG::IDR1.2(G59-E136) (juEx8633) | This study |
| CZ32252 | Pmec-7-mRFP + CBunc-119(jsIs973) III ; efa-6(ju1883) IV ; Pcol-19: mNG::3xFLAG::IDR1.3(G59-L95) (juEx8634) | This study |
| CZ32253 | Pmec-7-mRFP + CBunc-119(jsIs973); efa-6(ju1883ju1658); Pcol-19: mNG::3xFLAG::IDR1.3(G59-L95) (juEx8634) | This study |
| CZ32463 | Pmec-7-mRFP + CBunc-119(jsIs973); efa-6(ju1883ju1658); Pcol-19: mNG::3xFLAG::IDR1.4(I105-E136) (juEx8683) line-1 | This study |
| CZ32464 | Pmec-7-mRFP + CBunc-119(jsIs973); efa-6(ju1883ju1658); Pcol-19: mNG::3xFLAG::IDR1.4(I105-E136) (juEx8683) line-2 | This study |
| CZ32461 | Pmec-7-mRFP + CBunc-119(jsIs973); efa-6(ju1883ju1658); Pcol-19: mNG::3xFLAG::IDR1.4.1(I105-S120) (juEx8684) line-1 | This study |
| CZ32462 | Pmec-7-mRFP + CBunc-119(jsIs973); efa-6(ju1883ju1658); Pcol-19: mNG::3xFLAG::IDR1.4.1(I105-S120) (juEx8684) line-2 | This study |
| CZ32459 | Pmec-7-mRFP + CBunc-119(jsIs973); efa-6(ju1883ju1658); Pcol-19: mNG::3xFLAG::IDR1.4.2(A123-E136) (juEx8685) line-1 | This study |
| CZ32460 | Pmec-7-mRFP + CBunc-119(jsIs973); efa-6(ju1883ju1658); Pcol-19: mNG::3xFLAG::IDR1.4.2(A123-E136) (juEx8685) line-2 | This study |
| CZ32443 | Pmec-7-mRFP + CBunc-119(jsIs973); efa-6(ju1883ju1658); Pcol-19: mNG::3xFLAG::SP1(A123-A130) (juEx8686) line-1 | This study |
| CZ32444 | Pmec-7-mRFP + CBunc-119(jsIs973); efa-6(ju1883ju1658); Pcol-19: mNG::3xFLAG::SP1(A123-A130) (juEx8686) line-2 | This study |
| CZ32440 | Pmec-7-mRFP + CBunc-119(jsIs973); efa-6(ju1883ju1658); Pcol-19: mNG::3xFLAG::SP2(F131-E136) (juEx8687) line-1 | This study |
| CZ32441 | Pmec-7-mRFP + CBunc-119(jsIs973); efa-6(ju1883ju1658); Pcol-19: mNG::3xFLAG::SP2(F131-E136) (juEx8687) line-2 | This study |
| CZ32442 | Pmec-7-mRFP + CBunc-119(jsIs973); efa-6(ju1883ju1658); Pcol-19: mNG::3xFLAG::SP3(M126-I132) (juEx8688) line-1 | This study |
| CZ32465 | Pmec-7-mRFP + CBunc-119(jsIs973); efa-6(ju1883ju1658); Pcol-19: mNG::3xFLAG::SP3(M126-I132) (juEx8688) line-2 | This study |
| CZ32445 | Pmec-7-mRFP + CBunc-119(jsIs973); efa-6(ju1883ju1658); Pcol-19: mNG::3xFLAG::SP4(K128-R134) (juEx8689) line-1 | This study |
| CZ32446 | Pmec-7-mRFP + CBunc-119(jsIs973); efa-6(ju1883ju1658); Pcol-19: mNG::3xFLAG::SP4(K128-R134) (juEx8689) line-1 | This study |
| CZ28323 | P*mec-7*-mRFP + CB*unc-119* (*jsIs973*); *7xGFP11::efa-6(syb2598); Prgef-1-GFP1-10(juIs436)* | This study |
| CZ28451 | P*mec-7*-mRFP + CB*unc-119* (*jsIs973*); *7xGFP11::efa-6(syb2598); Pcol-19-GFP1-10(juEx8111)* | This study |
| CZ29472 | P*mec-7*-mRFP + CB*unc-119* (*jsIs973*); *efa-6(ju1658); Pmyo-3-mCherry(juEx4078)* | This study |

**Table S2:** Allele information and detection methods

| Allele name | Molecular detail | allele information and predicted protein/ Reference |
|---|---|---|
| *Pmec-7-mRFP + CBunc-119 (jsIs973)* | Genotyped using dissection scope | (Zheng et al., 2011) |
| Pmec-4:GFP(zdls5) | Genotyped using dissection scope | (Clark & Chiu, 2003) |
| GFP::EFA-6(ju1658) | Primers: YJ12371 + YJ12372<br>WT:1542bp KI:507bp | (Sandhu et al., 2024) |
| mSc::EFA-6(syb6998) | Primers: SD21116 + YJ12372<br>WT:no band KI: 429bp | (Sandhu et al., 2024) |
| GFP::EFA-6(ju1658ju1990) | Primers: SD20611 + SD20090<br>WT: 3764bp MUT: 360b<br><br>Primers: SD20611 + SD20610<br>WT: 1113bp MUT: no band | (Sandhu et al., 2024) |
| GFP::EFA-6(ju1658ju1883) | SD21128 + SD21129<br>WT: 739 bp MUT: no band<br><br>Primers: SD21128 + SD21127<br>WT: no band MUT: 436 bp | (Sandhu et al., 2024) |
| GFP::EFA-6(ju1658ju2193) | Primers: SD23144 + SD23145<br>WT: 365bp MUT: 509bp | 144 bp AID sequence inserted between nt 3945 and 3946 bp |
| Peft-3-TIR1(wrdSi22[eft-3p::TIR1:F2A:mTagBFP::tbb2 3' UTR]) | Primers: SD23146 + SD23147 + SD23148<br>WT:623bp MUT:881bp | (Zhang et al., 2015) |
| Prgef-1-TIR1(reSi7[rgef-1p::TIR1::F2A::mTagBFP2::AID*::NLS::tbb-2 3'UTR]) | Primers: SD23146 + SD23147 + SD23148<br>WT:623bp MUT:881bp | (Ashley et al., 2021) |
| Pcol-10-TIR1(reSi2[col-10p:TIR1::F2A::mTagBFP2::AID*::NLS::tbb-2 3'UTR]) | Primers: SD23149 + SD23150 + SD23148 WT:684bp MUT:881 | (Ashley et al., 2021) |
| mec-1(ju1688) | Primers: SD21845 + SD21846 Sanger sequencing of the 150bp product, also detected by lethargic phenotype. | T5800A nucleotide change, resulting in Tyr1498*. |
| mec-1(e1066) | Primers: SD21807 + SD21808 Sanger sequencing of the 572 bp product, also detected by lethargic animals. | G11705A nucleotide change at 5' intron splice site which result at truncated after Ku6. (Emtage et al., 2004) |
| him-4(ju1687) | Primers: SD22076 + SD22077 Sanger sequencing of the 257bp product | a G>A transition (GCC > ACC) in *him-4*, resulting in A140T missense mutation at VWA domain of HIM-4. |
| him-4(ju1768) | Primers: SD22330 + SD22331 Sanger sequencing of the 197bp product | a T>G transition (TGT > GGT) in *him-4*, resulting in C1022G missense mutation at 7th Ig repeat of HIM-4. |
| him-4(ju1689) | Primers: SD21847 + SD21848 Sanger sequencing of the 556bp product, also detected by higher male occurrence. | T13857A nucleotide resulting in Tyr4619*, truncating last three Ig domains, EGF domain and FC domain. |
| *him-4(ju2231)* | Primers: SD22076 + SD22077 | a G>A transition (GCC > ACC) in *him-4*, resulting in A140T missense mutation at VWA domain of HIM-4. |

| | | |
|---|---|---|
| | Digest the 257bp PCR product with MseI, WT product is cut into 161bp+96bp, MUT isn't cut. | |
| him-4(rh319) | Primers: SD21809 + SD21810<br>Sanger sequencing of the 286 bp product, also detected by exploding gut and higher male occurrence. | C2998T nucleotide change in cDNA resulting in Q875*.<br>(Vogel & Hedgecock, 2001) |
| *him-4::mNG(qy33)* | Primers: SD22336 + SD22329<br>WT: 405bp<br>MUT: ~1500bp | mNG insertion at C-terminus of HIM-4 with a 18aa linker sequence:<br>GASGASGASGASGASGAS<br>(Keeley et al., 2020) |
| *him-4::mNG(qy33ju2232)* | SD22076 + SD22077<br>Digest the 257bp PCR product with MseI, WT product is cut into 161bp+96bp, MUT isn't cut | a G>A transition (GCC > ACC) in *him-4*, resulting in A140T missense mutation at VWA domain of HIM-4. |
| *him-4::mNG(qy33ju2233)* | Primers: SD22330 + SD22331<br>Digest the 197bp PCR product with BspHI, WT product is cut into 127bp+70bp, MUT isn't cut. | a T>G transition (TGT > GGT) in *him-4*, resulting in C1022G missense mutation at 7th Ig repeat of HIM-4. |
| *him-4::mNG(qy33ju2235)* | Primers: SD22471 + SD22328<br>WT: 830 bp<br>MUT: ~500bp | Exact alignment unknown, contain ~300bp deletion between exon 57-59, this allele was generated in *qy33* and mNG was observed in *ju2235* animals, suggesting a in-frame deletion. |
| Pmec-4:BFP + Pttx-3:GFP *(juEx8629)* | GFP visible under dissection scope; BFP visible under compound scope | Pmec-4:BFP (50 ng/ul) co-injected with Pttx-3:GFP (50 ng/ul) |
| efa-6(ju1883) Pcol-10:mNG::3xFLAG(juSi415) | Visible under compound scope | Pcol-10:mNG::3xFLAG transgene inserted as single copy at cxTi10882 site on ChrIV. Expressed only in epidermis. |
| efa-6(ju1883) Pcol-10:mNG::3xFLAG::EFA-6(juSi408) | Visible under compound scope | Pcol-10:mNG::3xFLAG::EFA-6 transgene inserted as single copy at cxTi10882 site on ChrIV. Expressed only in epidermis. |
| efa-6(ju1883) Pcol-10:mNG::3xFLAG::EFA-6(ΔPH) (juSi746) | Visible under compound scope | Pcol-10:mNG::3xFLAG::EFA-6(ΔPH) transgene inserted as single copy at cxTi10882 site on ChrIV. Expressed only in epidermis.<br>375bp deletion, Pro563-Pro687<br>Left flanking sequence:<br>CGGCAGCCACAACAAATATATGA AGTTGAT<br>Right flanking sequence:<br>TTGCCAGTCACTTCGAAACCTGA AACTGCA |
| efa-6(ju1883) Pcol-10:mNG::3xFLAG::EFA-6(ΔCC) (juSi422) | Visible under compound scope | Pcol-10:mNG::3xFLAG::EFA-6(ΔCC) transgene inserted as single copy at |

| | | |
|---|---|---|
| | | cxTi10882 site on ChrIV. Expressed only in epidermis.<br>75bp deletion, Ile711-Ser735 deletion<br>Left flanking sequence: TCCACGGATTCCGTGTCTCGCTCCG<br>Right flanking sequence: GTATCACCTAATCAACCACAGCAGC |
| efa-6(ju1883) Pcol-10:mNG::3xFLAG::EFA-6(ΔC) (juSi417) | Visible under compound scope | Pcol-10:mNG::3xFLAG::EFA-6(ΔC) transgene inserted as single copy at cxTi10882 site on ChrIV. Expressed only in epidermis.<br>255bp deletion, Pro563-STOP817 flanking sequence: CGGCAGCCACAACAAATATATGAAGTTGAT |
| efa-6(ju1883) Pcol-10:mNG::3xFLAG::EFA-6(ΔSec7) (juSi421) | Visible under compound scope | Pcol-10:mNG::3xFLAG::EFA-6(ΔSec7) transgene inserted as single copy at cxTi10882 site on ChrIV. Expressed only in epidermis.<br>His343-Asp562<br>Left flanking sequence: AATAGTTCAGCATCAGCCAGTCCGTCACTC<br>Right flanking sequence: CCAGATTCAGTTGTCGAATATTATAGTGGA |
| efa-6(ju1883) Pcol-10:mNG::3xFLAG::EFA-6(ΔMTED) (juSi416) | Visible under compound scope | Pcol-10:mNG::3xFLAG::EFA-6(ΔMTED) transgene inserted as single copy at cxTi10882 site on ChrIV. Expressed only in epidermis.<br>54bp deletion, lys24 to arg41<br>Left flanking sequence - CATTGATGGAGCCCCTCGCCGGAATGTC<br>Right flanking sequence- AACGTTTCGAGCACTTATGCAAAGCTTC |
| efa-6(ju1883) Pcol-10:mNG::3xFLAG::EFA-6(ΔIDR) (juSi414) | Visible under compound scope | Pcol-10:mNG::3xFLAG::EFA-6(ΔIDR) transgene inserted as single copy at cxTi10882 site on ChrIV. Expressed only in epidermis.<br>849bp deletion, Gly58 to Ser340<br>Left flanking sequence- CAAAGCTTCTCGGCGACCAACTGCCTCCA<br>Right flanking sequence- CTCCATGCAACTTCTGAAAGTCGAGGAG |
| efa-6(ju1883) Pcol-10:mNG::3xFLAG::IDR(G59-L342)(juSi423) | Visible under compound scope | Pcol-10::mNG::3xFLAG::IDR(G59-L342) transgene inserted as single copy at cxTi10882 site on ChrIV. Expressed only in epidermis. |

| | | |
|---|---|---|
| efa-6(ju1883) Pcol-10:mNG::3xFLAG::EFA-6(PrLD) (juSi411) | Visible under compound scope | Pcol-10::mNG::3xFLAG::EFA-6(PrLD) transgene inserted as single copy at cxTi10882 site on ChrIV. Expressed only in epidermis. |
| efa-6(ju1883) Pcol-10:mNG::3xFLAG::ENIRM(A123-E136) (juSi749) | Visible under compound scope | Pcol-10::mNG::3xFLAG:: ENIRM(A123-E136) transgene inserted as single copy at cxTi10882 site on ChrIV. Expressed only in epidermis. |
| efa-6(ju1883) Pcol-10:mNG::3xFLAG::IDR(G59-E136) (juSi750) | Visible under compound scope | Pcol-10::mNG::3xFLAG:: IDR(G59-E136) inserted as single copy at cxTi10882 site on ChrIV. Expressed only in epidermis. |
| efa-6(ju1883) Pcol-10:mNG::3xFLAG::IDR(A123-L342) (juSi751) | Visible under compound scope | Pcol-10::mNG::3xFLAG::IDR(A123-L342) transgene inserted as single copy at cxTi10882 site on ChrIV. Expressed only in epidermis. |
| efa-6(ju1883) Pcol-10:mNG::3xFLAG::IDR Δ ENIRM(A123-E136) (juSi752) | Visible under compound scope | Pcol-10::mNG::3xFLAG::IDRΔENIRM(A123-E136) transgene inserted as single copy at cxTi10882 site on ChrIV. Expressed only in epidermis. |
| *7xGFP11::efa-6(syb2598)* | Primers: YJ12371 + YJ12372 WT:945bp KI:507bp | 7 copies of GFP11 insertion in-frame to ATG of EFA-6 |
| *Prgef-1-GFP1-10(juIs436)* | Detect by Pttx-3-rfp using compound scope | Expression of GFP1-10 in pan-neuron (Noma K et al., 2017) |
| *Pcol-19-GFP1-10(juEx8111)* | Correlate with co-injection marker: Pmyo-2-mCherry, visible under dissection scope | Extrachromosomal expression of GFP1-10 in adult epidermis |
| Pcol-19:mNG::3xFLAG(juEx8682) | Visible under compound scope | pCZGY3757 (2 ng/ul) injected with Papq-5:mKO2 (50 ng/ul) |
| Pcol:19:mNG::3xFLAG::IDR(G59-L342)(juEx8630) | Visible under compound scope | pCZGY3747 (2 ng/ul) injected with Papq-5:mKO2 (50 ng/ul) |
| Pcol-19:mNG::3xFLAG::IDR1(G59-Q175)(juEx8631) | Visible under compound scope | pCZGY3753 (2 ng/ul) injected with Papq-5:mKO2 (50 ng/ul) |
| Pcol-19:mNG::3xFLAG::IDR2(F176-L342)(juEx8632) | Visible under compound scope | pCZGY3750 (2 ng/ul) injected with Papq-5:mKO2 (50 ng/ul) |
| Pcol-19:mNG::3xFLAG::IDR1.2(G59-E136)(juEx8633) | Visible under compound scope | pCZGY3752 (2 ng/ul) injected with Papq-5:mKO2 (50 ng/ul) |
| Pcol-19:mNG::3xFLAG::IDR1.3(G59-L95)(juEx8634) | Visible under compound scope | pCZGY3751 (2 ng/ul) injected with Papq-5:mKO2 (50 ng/ul) |
| Pcol-19:mNG::3xFLAG::IDR1.4(I105-E136) (juEx8683) | Visible under compound scope | pCZGY3754 (2 ng/ul) injected with Papq-5:mKO2 (50 ng/ul) |
| Pcol-19:mNG::3xFLAG::IDR1.4.1(I105-S120) (juEx8684) | Visible under compound scope | pCZGY3755 (2 ng/ul) injected with Papq-5:mKO2 (50 ng/ul) |
| Pcol-19:mNG::3xFLAG::IDR1.4.2(A123-E136) (juEx8685) | Visible under compound scope | pCZGY3756 (2 ng/ul) injected with Papq-5:mKO2 (50 ng/ul) |

| | | |
|---|---|---|
| Pcol-19:mNG::3xFLAG::SP-1(A123-A130) (juEx8686) | Visible under compound scope | pCZGY3762 (2 ng/ul) injected with Papq-5:mKO2 (50 ng/ul) |
| Pcol-19:mNG::3xFLAG::SP-2(F131-E136) (juEx8687) | Visible under compound scope | pCZGY3760 (2 ng/ul) injected with Papq-5:mKO2 (50 ng/ul) |
| Pcol-19:mNG::3xFLAG::SP-3(M126-I132) (juEx8688) | Visible under compound scope | pCZGY3761 (2 ng/ul) injected with Papq-5:mKO2 (50 ng/ul) |
| Pcol-19:mNG::3xFLAG::SP-4(K128-R134) (juEx8689) | Visible under compound scope | pCZGY3763 (2 ng/ul) injected with Papq-5:mKO2 (50 ng/ul) |
| Pcol-19:mNG::3xFLAG::IDR ∆ENIRM(A123-E136) (juEx8690) | Visible under compound scope | pCZGY3767 (2 ng/ul) injected with Papq-5:mKO2 (50 ng/ul) |
| Pcol-19:mNG::3xFLAG::ENIRM 14aa Scram1 (juEx8691) | Visible under compound scope | pCZGY3758 (2 ng/ul) injected with Papq-5:mKO2 (50 ng/ul) |
| Pcol-19:mNG::3xFLAG::ENIRM 14aa Scram2 (juEx8692) | Visible under compound scope | pCZGY3759 (2 ng/ul) injected with Papq-5:mKO2 (50 ng/ul) |
| Pcol-19:mNG::3xFLAG::IDR(G59-T122) (juEx8693) | Visible under compound scope | pCZGY3765 (2 ng/ul) injected with Papq-5:mKO2 (50 ng/ul) |
| Pcol-19:mNG::3xFLAG::IDR(S137-L342) (juEx8694) | Visible under compound scope | pCZGY3766 (2 ng/ul) injected with Papq-5:mKO2 (50 ng/ul) |
| Pcol-19:mNG::3xFLAG::IDR(A123-L342) (juEx8695) | Visible under compound scope | pCZGY3764 (2 ng/ul) injected with Papq-5:mKO2 (50 ng/ul) |
| *Pmyo-3-mCherry(juEx4078)* | Visible under dissection scope | Extrachromosomal expression of mCherry labeled muscle cells |

**Table S3:** Genotyping, genome editing, and cloning primers

| Primer name | Sequence |
|---|---|
| **Genotyping primers** | |
| YJ12371 | ctcacctccatcacgtgttt |
| YJ12372 | tgaggcctcaaatcgcagagc |
| SD21116 | gcgttccgagggacgtcact |
| SD20611 | gaaataatgctctgcgatttgag |
| SD20090 | atactcctccagtggctcctcgac |
| SD20610 | ggttaaaaatcaatggaaaatcg |
| SD21128 | gctcgaaatctgtatgagttgaaga |
| SD21129 | ttgattctattgcgagaatatcttg |
| SD21128 | gctcgaaatctgtatgagttgaaga |
| SD21127 | ctcgaaactcaaaagcacccat |
| SD23144 | CCGTCGAAACATTCAGTGGT |
| SD23145 | ATACTCCTCCAGTGGCTCCTCGAC |
| SD23146 | tgtcgaccgctagtgtagcttac |
| SD23147 | cgtctctccacgatttacacactatttg |
| SD23148 | CTGGCCGTCGTTTTACAGGA |
| SD23149 | ACCGGAAACCAAAGGACGAGAG |
| SD23150 | ACGCCCAGGAGAACACGTTAG |
| SD21845 | AAGCTTGTCTGCAACCATTC |
| SD21846 | TCTCTCACAGATCCATTGGC |
| SD21807 | ACACTTGGGTTGTCAGAAGC |
| SD21808 | GCTAAATCCTCTCTGCCCTC |
| SD22076 | GTATTCTCAAGGCACTTCAAAT |
| SD22077 | TTCTAATACGGTACTGACATCACTC |
| SD22330 | AAGGAGGAATCGTGTTCAAG |
| SD22331 | CGTGACAGTGAAATCTCTGC |
| SD21847 | AGGACCAAGATCCTCTCGAAC |
| SD21848 | GCTTGGTTTCTCTCATAACAGGTGATC |
| SD21809 | GGCTACACTCAACAATCCAAGATAT |
| SD21810 | GACTGGTTTATCATCCTTAATCCAT |
| SD22336 | GGTGATCATCACAAAGTACGAG |
| SD22329 | gtggggtctggcatactttgc |
| SD22471 | CCATTGACAACCACAATCATTCCAGg |
| SD22328 | CGCACTCATCGAGATCTTCGC |
| **Genome editing primers** | |
| SD21595 | TCTTCGAATCTAGAACGACTCCAAACACGGCGGCGAGCAATAGTTCAGCATCAG CCAGTCCGTCAGGAGCCGGAGCCCCTAAAGATCCAG |
| SD21596 | aataaattttctggaaattcctggaattctcaaaaaacatttcccatacTTTCAGAAGTTGCATGGAGCTT CACGAACGCCGCCGCCTC |
| **Cloning primers** | |
| SD22111 | CCAATATACCAAACATAACTGTTTGAAGTTCCTATTCTCTAGAAAGTATAGGAAC TTCGTCAATCAACTTCATATATTTGTTGTGGCTG |
| SD21602 | atggtgtcgaagggagaagaggataacatg |
| SD21637 | cataatttacagtaaaattatcaaaaaatagcaaaaatcaatagaggaccgctacacAaccgatcttcatcccttcaa catttggcag |
| SD21638 | CCAATATACCAAACATAACTGTTTGAAGTTCCTATTCTCTAGAAAGTATAGGAAC TTCGTCAATGAGCAGCAGCTCCATTAACAGCATC |
| SD23137 | CACTGCCGCCACCACCGCCGGATCCTCGCTCGGATGGCTTTCCAGAATCTCCA CTTTTTCCCCATGGAGGCAGTTGGTCGCCGAGAAGC |
| SD23138 | TTTCTCTTAATCGGAACGTTTCGAGCACTTATGCAAAGCTTCTCGGCGACCAAC TGCCTCCATGGGGAAAAAGTGGAGATTCTGGAAAG |

| | |
|---|---|
| SD23139 | GCTCTCGGCGGATCGAAGTGATACTCCTCCAGTGGCTCCTCGACTTTCAGAAG TTGCATGGAGATTTCCTCCAGTAGATGAAGCTGCTG |
| SD23140 | ATTGGCAAAAGGATGGACCACCACCACCACCAGCTGCAGCAGCTTATCTACTG GAGGAAATCTCCATGCAACTTCTGAAAGTCGAGG |
| SD23151 | TTCCCAATATACCAAACATAACTGTTTGAAGTTCCTATTCTCTAGAAAGTATAGG AACTTCGCTActtgtcatcgtcatccttgtaatc |
| SD21961 | TCAATCAACTTCATATATTTGTTGTGGC |
| SD21690 | GACATTCCGGCGAGGGGCTCCATCAATG |
| SD21691 | AACGTTTCGAGCACTTATGCAAAGCTTC |
| SD21692 | TGGAGGCAGTTGGTCGCCGAGAAGCTTTG |
| SD21693 | CTCCATGCAACTTCTGAAAGTCGAGGAG |
| SD22374 | caaagaccatgacggtgattataaagatcatgacatcgattacaaggatgacgatgacaagGGTACCACAG TTGCATCGAGCATTCAC |
| SD22375 | GAGCAATTATTTTTTATAATGCCAACTTTGTACAAGAAAGCTGGGTCGAATTCGC CCTTCAGAGTGACGGACTGGCTGATGCTG |
| SD22376 | CAGAGGCGGTGGCTCTTGACAGCTGATGTGGGTGAATGCTCGATGCAACTGTG GTACCcttgtcatcgtcatccttgtaatcgatg |
| SD22377 | CTAGAACGACTCCAAACACGGCGGCGAGCAATAGTTCAGCATCAGCCAGTCCG TCACTCTGAAGGGCGAATTCGACCCAGCTTTCTTG |
| SD22378 | GAGTGACGGACTGGCTGATGCTGAAC |
| SD22379 | CCAGATTCAGTTGTCGAATATTATAGTGG |
| SD22830 | ATCAACTTCATATATTTGTTGTGGCTGCCG |
| SD22831 | TTGCCAGTCACTTCGAAACCTGAAACTGCA |
| SD22832 | CGGAGCGAGACACGGAATCCGTGGAAGTCG |
| SD22833 | GTATCACCTAATCAACCACAGCAGCTG |
| SD22834 | CCAATATACCAAACATAACTGTTTGAAGTTCCTATTCTCTAGAAAGTATAGGAAC TTCGTCAGAGTGACGGACTGGCTGATGCTGAACT |
| SD22839 | TTGTTCTTCATCAACAACCATCACTTCA |
| SD22840 | CTCCATTCGTTCAATTGGAGCTGTTTTAAG |
| SD22841 | CAGTTTCTGGGCCGCCGCTCCGTTTCGATT |
| SD22842 | TGAAGGGCGAATTCGACCCAGCTTTCTTGT |
| SD22843 | TTTATTGTAGTATCAAACGACATGAAATCT |
| SD22844 | ACCcttgtcatcgtcatccttgtaatcgatg |
| SD22949 | GCTATTCGAATGCTTAAAACAGCTCCAATTGAACGAATG |
| SD22950 | TGAAGACTCTTCTGACGTTTTTCGTGCCAAAC |
| SD22951 | TGCCAAACTACCACGTCTTTCGATACCcttgtcatcgtcatccttgtaatcgatgtc |
| SD22952 | CGAAAAACGTCAGAAGAGTCTTCATGAAGGGCGAATTCGACCCAGCTTTCTTGT ACAAAG |
| SD22953 | AGCTGTTTTAAGCATTCGAATAGCACCcttgtcatcgtcatccttgtaatcgatgtc |
| SD22954 | CCAATTGAACGAATGGAGTGAAGGGCGAATTCGACCCAGCTTTCTTGTACAAAG TTGGC |
| SD22955 | TGGTGAAGACTCTTCTGACGTTTTTCGTGCCAAACTACCACGTCTTTCGATACCc ttgtcatcgtcatccttgtaatcgatgtcatg |
| SD22956 | ACAGCTATTCGAATGCTTAAAACAGCTCCAATTGAACGAATGGAGTGAAGGGCG AATTCGACCCAGCTTTCTTGTACAAAGTTGGC |
| SD22957 | CCAATATACCAAACATAACTGTTTGAAGTTCCTATTCTCTAGAAAGTATAGGAAC TTCGTCACTCCATTCGTTCAATTGGAGCTGT |
| SD23064 | CGCCCTTCACATCTCAATTTCTGGAGCTGTCATTTTAATTCGTCGAGCAAGACCc ttgtcatcgtcatccttgtaatcgatgtc |
| SD23065 | CGCCCTTCACTCCATAATAGCTGTAGCTCGTCGAAGTTTTTCCATAATTGGACCc ttgtcatcgtcatccttg |
| SD23066 | CGCCCTTCACTCCATTCGTTCAATTGGACCcttgtcatcgtcatccttgtaatcgatgtcatgatcttta taatcaccg |

| | |
|---|---|
| SD23067 | CGCCCTTCAAATTGGAGCTGTTTTAAGCATACCcttgtcatcgtcatccttgtaatcgatgtcatgatctttataatcacc |
| SD23068 | CGCCCTTCAAGCTGTTTTAAGCATTCGAATAGCACCcttgtcatcgtcatccttg |
| SD23069 | AATTCGACCCAGCTTTCTTGTACAAAGTTGGCATTATAAAAAATAATTGCTCATC |
| SD23070 | CGCCCTTCATCGTTCAATTGGAGCTGTTTTACCcttgtcatcgtcatccttgtaatcgatgtcatgatctttataatcacc |
| SD23142 | TGTTGGTGAAGACTCTTCTGACGTTTTTCGTGCCAAAC |
| SD23143 | AGCACTGATGTAGAGGAAAGTGAAGAAGAAACTGTAATG |

**Table S4:** Overexpression constructs generated for systematic analysis of the EFA-6 IDR

| Plasmid # | Name | Construction |
|---|---|---|
| pCZGY 3757 | Pcol-19: mNG::3xFLAG | Plasmid pCZGY3721 (mNG::3xFLAG -pCR8) was generated by TOPO cloning. The resulting construct was subsequently recombined with the Pcol-19 Gateway destination vector pCZGY1434 using Gateway recombination cloning to generate an epidermal-specific expression construct. |
| pCZGY 3747 | Pcol-19: mNG::3xFLAG::IDR(G59-L342) | Full-length EFA-6 IDR(G59-L342) was cloned into a pCR8 vector containing mNG::3×FLAG (pCZGY3721) using Gibson assembly with primers SD22376, SD22377, SD22374, and SD22375, generating plasmid pCZGY3713 (mNG::3×FLAG::IDR(G59-L342)-pCR8). The resulting construct was subsequently recombined with the Pcol-19 Gateway destination vector pCZGY1434 using Gateway recombination cloning to generate an epidermal-specific expression construct. |
| pCZGY 3753 | Pcol-19: mNG::3xFLAG::IDR1(G59-Q175) | Plasmid pCZGY3717 (mNG::3×FLAG:: IDR1(G59-Q175)-pCR8) was generated from pCZGY3713 (mNG::3×FLAG::IDR(G59-L342)-pCR8) by PCR-based mutagenesis using primers SD22842 and SD22839 to delete the C-terminal 167 amino acids of the EFA-6 IDR. The resulting construct was subsequently recombined with the Pcol-19 Gateway destination vector pCZGY1434 using Gateway recombination cloning to generate final an epidermal-specific expression construct. |
| pCZGY 3750 | Pcol-19: mNG::3xFLAG::IDR2(F176-L342) | Plasmid pCZGY3714 (mNG::3×FLAG::I IDR2(F176-L342)-pCR8) was generated from pCZGY3713 (mNG::3×FLAG::IDR(G59-L342)-pCR8) by PCR-based mutagenesis using primers SD22844 and SD22843 to delete the N-terminal 117 amino acids of the EFA-6 IDR. The resulting construct was subsequently recombined with the Pcol-19 Gateway destination vector pCZGY1434 using Gateway recombination cloning to generate final epidermal-specific expression construct. |
| pCZGY 3752 | Pcol-19: mNG::3xFLAG::IDR1.2(G59-E136) | Plasmid pCZGY3716 (mNG::3×FLAG:: IDR1.2(G59-E136)-pCR8) was generated from pCZGY3713 (mNG::3×FLAG::IDR(G59-L342)-pCR8) by PCR-based mutagenesis using primers SD22842 and SD22840 to delete the C-terminal 206 amino acids of the EFA-6 IDR. The resulting construct was subsequently recombined with the Pcol-19 Gateway destination vector pCZGY1434 using Gateway recombination cloning to generate final epidermal-specific expression construct. |
| pCZGY 3751 | Pcol-19: mNG::3xFLAG::IDR1.3(G59-L95) | Plasmid pCZGY3715 (mNG::3×FLAG::IDR1.3(G59-L95)-pCR8) was generated from pCZGY3713 (mNG::3×FLAG::IDR(G59-L342)-pCR8) by PCR-based mutagenesis using primers SD22842 and SD22841 to delete the C-terminal 247 amino acids of the EFA-6 IDR. The resulting construct was subsequently recombined with the Pcol-19 Gateway destination vector pCZGY1434 using Gateway recombination cloning to generate an epidermal-specific expression construct. |
| pCZGY 3754 | Pcol-19: mNG::3xFLAG::IDR1.4(I105-E136) | Plasmid pCZGY3718 (mNG::3×FLAG:: IDR1.4(I105-E136)-pCR8) was generated from pCZGY3716 (mNG::3×FLAG::IDR1.2(G59-E136)-pCR8) by PCR-based mutagenesis using primers SD22955 and SD22956. The resulting construct was subsequently recombined with the Pcol- |

| | | |
|---|---|---|
| | | 19 Gateway destination vector pCZGY1434 using Gateway recombination cloning to generate final epidermal-specific expression construct. |
| pCZGY 3755 | Pcol-19: mNG::3xFLAG::IDR1.4.1(I 105-S120) | Plasmid pCZGY3719 (mNG::3×FLAG::IDR1.4.1(I105-S120)-pCR8) was generated from pCZGY3718 (mNG::3×FLAG:: IDR1.4(I105-E136)-pCR8) by PCR-based mutagenesis using primers SD22951 and SD22952. The resulting construct was subsequently recombined with the Pcol-19 Gateway destination vector pCZGY1434 using Gateway recombination cloning to generate final epidermal-specific expression construct. |
| pCZGY 3756 | Pcol-19: mNG::3xFLAG::IDR1.4.2( A123-E136) | Plasmid pCZGY3720 (mNG::3×FLAG::IDR1.4.2(A123-E136)-pCR8) was generated from pCZGY3718 (mNG::3×FLAG:: IDR1.4(I105-E136)-pCR8) by PCR-based mutagenesis using primers SD22953 and SD22954. The resulting construct was subsequently recombined with the Pcol-19 Gateway destination vector pCZGY1434 using Gateway recombination cloning to generate final epidermal-specific expression construct. |
| pCZGY 3762 | Pcol-19: mNG::3xFLAG::SP1(A123 -A130) | Plasmid pCZGY3726 (mNG::3×FLAG::SP1(A123-A130)-pCR8) was generated from pCZGY3721 (mNG::3×FLAG-pCR8) by PCR-based mutagenesis using primers SD23068 and SD23069. The resulting construct with was subsequently recombined with the Pcol-19 Gateway destination vector pCZGY1434 using Gateway recombination cloning to generate final epidermal-specific expression construct. |
| pCZGY 3760 | Pcol-19: mNG::3xFLAG::SP2(F131-E136) | Plasmid pCZGY3724 (mNG::3×FLAG::SP2(F131-E136)-pCR8) was generated from pCZGY3721 (mNG::3×FLAG-pCR8) by PCR-based mutagenesis using primers SD23066 and SD23069. The resulting construct with was subsequently recombined with the Pcol-19 Gateway destination vector pCZGY1434 using Gateway recombination cloning to generate final epidermal-specific expression construct. |
| pCZGY 3761 | Pcol-19: mNG::3xFLAG::SP3(M126 -I132) | Plasmid pCZGY3725 (mNG::3×FLAG::SP3(M126-I132)-pCR8) was generated from pCZGY3721 (mNG::3×FLAG-pCR8) by PCR-based mutagenesis using primers SD23067 and SD23069. The resulting construct with was subsequently recombined with the Pcol-19 Gateway destination vector pCZGY1434 using Gateway recombination cloning to generate final epidermal-specific expression construct. |
| pCZGY 3763 | Pcol-19: mNG::3xFLAG::SP4(K128 -R134) | Plasmid pCZGY3727 (mNG::3×FLAG::SP4(K128-R134)-pCR8) was generated from pCZGY3721 (mNG::3×FLAG-pCR8) by PCR-based mutagenesis using primers SD23070 and SD23069. The resulting construct with was subsequently recombined with the Pcol-19 Gateway destination vector pCZGY1434 using Gateway recombination cloning to generate final epidermal-specific expression construct. |
| pCZGY 3758 | Pcol-19: mNG::3xFLAG::ENIRM 14aa Scram1 | Plasmid pCZGY3722 (mNG::3×FLAG::ENIRM 14aa Scram-pCR8) was generated from pCZGY3721 (mNG::3×FLAG-pCR8) by PCR-based mutagenesis using primers SD23064 and SD23069. The resulting construct with was subsequently recombined with the Pcol-19 Gateway destination vector pCZGY1434 using Gateway recombination cloning to generate final epidermal-specific expression construct. |
| pCZGY 3759 | Pcol-19: mNG::3xFLAG::ENIRM 14aa Scram2 | Plasmid pCZGY3723 (mNG::3×FLAG::ENIRM 14aa Scram-pCR8) was generated from pCZGY3721 (mNG::3×FLAG-pCR8) by PCR-based mutagenesis using primers SD23065 and |

| | | |
|---|---|---|
| | | SD23069. The resulting construct with was subsequently recombined with the Pcol-19 Gateway destination vector pCZGY1434 using Gateway recombination cloning to generate final epidermal-specific expression construct. |
| pCZGY 3765 | Pcol-19: mNG::3xFLAG::IDR(G59-T122) | Plasmid pCZGY3729 (mNG::3×FLAG::IDR(G59-T122)-pCR8) was generated from pCZGY3713 (mNG::3×FLAG::IDR(G59-L342)-pCR8) by PCR-based mutagenesis using primers SD22950 and SD22842. The resulting construct with was subsequently recombined with the Pcol-19 Gateway destination vector pCZGY1434 using Gateway recombination cloning to generate final epidermal-specific expression construct. |
| pCZGY 3766 | Pcol-19: mNG::3xFLAG::IDR(S137-L342) | Plasmid pCZGY3730 (mNG::3×FLAG::IDR(S137-L342)-pCR8) was generated from pCZGY3713 (mNG::3×FLAG::IDR(G59-L342)-pCR8) by PCR-based mutagenesis using primers SD22844 and SD23143. The resulting construct with was subsequently recombined with the Pcol-19 Gateway destination vector pCZGY1434 using Gateway recombination cloning to generate final epidermal-specific expression construct. |
| pCZGY 3764 | Pcol-19: mNG::3xFLAG::IDR(A123-L342) | Plasmid pCZGY3728 (mNG::3×FLAG:: IDR(A123-L342)-pCR8) was generated from pCZGY3713 (mNG::3×FLAG::IDR(G59-L342)-pCR8) by PCR-based mutagenesis using primers SD22844 and SD22949. The resulting construct with was subsequently recombined with the Pcol-19 Gateway destination vector pCZGY1434 using Gateway recombination cloning to generate final epidermal-specific expression construct. |
| pCZGY 3767 | Pcol-19: mNG::3xFLAG::IDRΔENIRM(A123-E136) | Plasmid pCZGY3731 (mNG::3×FLAG::IDRΔENIRM(A123-E136)-pCR8) was generated from pCZGY3713 (mNG::3×FLAG::IDR(G59-L342)-pCR8) by PCR-based mutagenesis using primers SD23142 and SD23143. The resulting construct with was subsequently recombined with the Pcol-19 Gateway destination vector pCZGY1434 using Gateway recombination cloning to generate final epidermal-specific expression construct. |
| pCZGY 3784 | Pmec-4:BFP | Plasmid pCZGY3784 (Pmec-4:BFP) was generated by Gateway recombination cloning of pCZGY3783 (BFP-pCR8) with Pmec-4 Gateway destination vector pCZGY553. |

**Table S5:** Plasmids generated for CRISPR-assisted single-copy insertion (CasSCI) of EFA-6 constructs

| **Plasmid #** | **construct/backbone** | **Detail** |
|---|---|---|
| pCZGY3768 | Pcol-10: mNG::3xFLAG::EFA-6/ pCZGY2729 | Full-length EFA-6<br>Starting clone-pCZGY3704(mNG::3xFLAG::EFA-6-pCR8)<br>Intermediate clone: pCZGY3732 (Pcol-10: mNG::3xFLAG::EFA-6)<br>Primers used cloning into pCZGY2729: SD21637 + SD21638 |
| pCZGY3772 | Pcol-10: mNG::3xFLAG::EFA-6(ΔIDR)/ pCZGY2729 | EFA-6 lacking IDR<br>Deletion generated by pcr-mutagenesis.<br>Mutagenesis Primers: SD21692 + SD21693<br>Starting clone-pCZGY3706 (mNG::3xFLAG::EFA-6(ΔIDR)-pCR8)<br>Intermediate clone: pCZGY3735(Pcol-10: mNG::3xFLAG::EFA-6(ΔIDR))<br>Primers used cloning into pCZGY2729: SD21637 + SD21638 |
| pCZGY3769 | Pcol-10: mNG::3xFLAG/ pCZGY2729 | mNeonGreen-only control<br>Intermediate clone: pCZGY3732 (Pcol-10::mNG::3xFLAG::EFA-6)<br>Primers used cloning into pCZGY2729: SD23151 + SD21637 |
| pCZGY3771 | Pcol-10: mNG::3xFLAG::EFA-6(ΔMTED)/ pCZGY2729 | EFA-6 lacking the MTED<br>Deletion generated by pcr-mutagenesis.<br>Mutagenesis Primers: SD21690 + SD21691<br>Starting clone-pCZGY3705 (mNG::3xFLAG::EFA-6(ΔMTED)-pCR8)<br>Intermediate clone: pCZGY3734(Pcol-10: mNG::3xFLAG::EFA-6(ΔIDR))<br>Primers used cloning into pCZGY2729: SD21637 + SD21638 |
| pCZGY3773 | Pcol-10: mNG::3xFLAG::EFA-6(ΔC)/ pCZGY2729 | EFA-6 lacking the C-terminal PH and CC domains<br>Deletion generated by pcr-mutagenesis.<br>Mutagenesis Primers: SD21602 + SD21961<br>Starting clone-pCZGY3707(mNG::3xFLAG::EFA-6(ΔC)-pCR8)<br>Intermediate clone: pCZGY3736(Pcol-10: mNG::3xFLAG::EFA-6(ΔC))<br>Primers used cloning into pCZGY2729: SD21637 + SD22111 |
| pCZGY3774 | Pcol-10: mNG::3xFLAG::EFA-6(ΔSec7)/ pCZGY2729 | EFA-6 lacking the Sec7 catalytic domain<br>Deletion generated by pcr-mutagenesis.<br>Mutagenesis Primers: SD22378 + SD22379<br>Starting clone-pCZGY3709(mNG::3xFLAG::EFA-6(ΔSec7)-pCR8)<br>Intermediate clone: pCZGY3738(Pcol-10: mNG::3xFLAG::EFA-6(Δsec7))<br>Deletion generated by pcr-mutagenesis.<br>Primers used cloning into pCZGY2729: SD21637 + SD21638 |

| | | |
|---|---|---|
| pCZGY3782 | Pcol-10: mNG::3xFLAG::EFA-6(ΔCC)/ pCZGY2729 | EFA-6 lacking the coiled-coil domain<br>Deletion generated by pcr-mutagenesis.<br>Mutagenesis Primers: SD22832 + SD22833<br>Starting clone-pCZGY3711(mNG::3xFLAG::EFA-6(ΔCC)-pCR8)<br>Intermediate clone: pCZGY3739(Pcol-10: mNG::3xFLAG::EFA-6(ΔCC))<br>Deletion generated by pcr-mutagenesis.<br>Primers used cloning into pCZGY2729: SD21637 + SD21638 |
| pCZGY3777 | Pcol-10: mNG::3xFLAG::IDR/ pCZGY2729 | Full-length EFA-6 intrinsically disordered region<br>Starting clone-pCZGY3713(mNG::3xFLAG::IDR(G59-L342)-pCR8)<br>Intermediate clone: pCZGY3742(Pcol-10:mNG::3xFLAG::IDR(G59-L342))<br>Primers used cloning into pCZGY2729: SD21637 + SD22834 |
| pCZGY3775 | Pcol-10: mNG::3xFLAG::EFA-6(ΔPH)/ pCZGY2729 | EFA-6 lacking the pleckstrin homology domain<br>Deletion generated by pcr-mutagenesis.<br>Mutagenesis Primers: SD22830 + SD22831<br>Starting clone-pCZGY3710(mNG::3xFLAG::EFA-6(ΔPH)-pCR8)<br>Intermediate clone: pCZGY3740(Pcol-10: mNG::3xFLAG::EFA-6(ΔPH))<br>Deletion generated by pcr-mutagenesis.<br>Primers used cloning into pCZGY2729: SD21637 + SD21638 |
| pCZGY3770 | Pcol-10: mNG::3xFLAG::EFA-6(PrLD)/ pCZGY2729 | EFA-6 IDR replaced with TIAR2 PrLD sequence<br>Starting construct generated by Gibson assemby; primers used: SD23138 + SD23139; SD23137+ SD23140<br>Starting clone-pCZGY3708(mNG::3xFLAG::EFA-6(PrLD)-pCR8)<br>Intermediate clone: pCZGY3733(Pcol-10:mNG::3xFLAG::EFA-6(PrLD))<br>Primers used cloning into pCZGY2729: SD21637 + SD21638 |
| pCZGY3778 | Pcol-10: mNG::3xFLAG::IDR1.2(G59-E136)/ pCZGY2729 | EFA-6 IDR N-terminus flanking sequence with ENIRM 14aa motif<br>Starting clone generated by pcr-mutagenesis<br>Mutagenesis primers: SD22842 and SD22840<br>Starting clone-pCZGY3716 (mNG::3xFLAG::IDR1.2(G59-E136)-pCR8)<br>Intermediate clone: pCZGY3743 Pcol-10: mNG::3xFLAG::IDR1.2(G59-E136)<br>Primers used cloning into pCZGY2729: SD21637 and SD22957 |
| pCZGY3779 | Pcol-10: mNG::3xFLAG::ENIRM(A123-E136)/ pCZGY2729 | EFA-6 IDR-derived 14-amino-acid ENIRM sequence<br>Starting clone generated by pcr-mutagenesis.<br>Mutagenesis primers: SD22953 and SD22954 |

| | | |
|---|---|---|
| | | Starting clone-pCZGY3720 (mNG::3xFLAG:: ENIRM(A123-E136)-pCR8)<br>Intermediate clone: pCZGY3745 (Pcol-10: mNG::3xFLAG:: ENIRM(A123-E136))<br>Primers used cloning into pCZGY2729: SD21637 and SD22957 |
| pCZGY3780 | Pcol-10: mNG::3xFLAG::IDR(A123-L342)/ pCZGY2729 | ENIRM motif with IDR C-terminal flanking region<br>Starting clone generated by pcr-mutagenesis<br>Mutagenesis primers: SD22844 and SD22949<br>Starting clone-pCZGY3728 (mNG::3xFLAG::IDR(A123-L342)-pCR8)<br>Intermediate clone: pCZGY3748 (Pcol-10: mNG::3xFLAG::IDR(A123-L342)<br>Primers used cloning into pCZGY2729: SD21637 + SD22834 |
| pCZGY3781 | Pcol-10: mNG::3xFLAG::IDRΔENIRM(A123-E136)/ pCZGY2729 | EFA-6 IDR lacking the ENIRM motif<br>Starting clone generated by pcr-mutagenesis<br>Mutagenesis primers: SD23142 and SD23143<br>Starting clone-pCZGY3731 (mNG::3xFLAG::IDRΔENIRM(A123-E136)-pCR8)<br>Intermediate clone: pCZGY3732 (Pcol-10: mNG::3xFLAG::IDRΔENIRM)<br>Primers used cloning into pCZGY2729: SD21637 + SD22834 |

Note: All constructs encode N-terminally mNG-tagged proteins driven by the epidermis-specific Pcol-10 promoter. Intermediate plasmids were generated by Gateway LR recombination between the respective entry clones and the Pcol-10 destination vector pCZGY3639. The resulting plasmids were used as templates to amplify the desired expression cassettes, which were subsequently cloned into the CasSCI vector pCZGY2729 by Gibson assembly.

**Table S6**: CRISPR/Cas9 genome editing information

| allele | crRNA (including **PAM** site) | Flanking Sequence |
|---|---|---|
| *ju2231* | TAAGTGGTAATCCTTTGATC**GGG** | GTTCACTGATACCCGATCTAAGGACTACCACTTGGAAGATGAAGTTCTCAATACAATTC |
| *ju2232* | | |
| *ju2233* | GGTGTTCCCGAAACCACACA**TGG** | TGCAATTCCAGGTGTCGTTTCGGGAACCCCACCACCA |
| *ju2235* | CCCAACTGATGATTATCATG**AGG** | Exact alignment unknown, ~300bp deletion in exon 57-59 |
| *ju2193* | TCAGAAGTTGCATGGAGTGA**CGG** | CAATAGTTCAGCATCAGCCAGTCCGTCAGGAGCCGGAGCCCCTAAAGATCCAGCCAAACCTCCGGCCAAGGCACAAGTTGTGGGATGGCCACCGGTGAGATCATACCGGAAGAACGTGATGGTTTCCTGCCAAAAATCAAGCGGTGGCCCGGAGGCGGCGGCGTTCGTGAAGCTCCATGCAACTTCTGAAA |

Note: Synonymous mutations are highlighted in GREEN, Mutants SNP/inserted sequence are marked in RED, Flanking sequence around edited region is underlined)

**Table S7:** Summary of phenotypes associated with different *him-4* alleles

| Allele | Effect on EFA-6 patterning | Neurite Ensheathment | Vulva bursting | Random male generation |
|---|---|---|---|---|
| *mec-1(ju1688, T1498STOP)* | Ectopic foci formation in the proximal neurite | Completely lost | <1% animals | Absent |
| *mec-1(e1066)* | Ectopic foci formation in the proximal neurite | Completely lost | <1% animals | Absent |
| *him-4*(rh319, Q875STOP) | Ectopic foci formation in the proximal neurite | Completely lost | >90% animals | Present |
| *him-4*(ju1687, A140T) | Ectopic foci formation in the proximal neurite | Severe loss | <1% animals | Present |
| *him-4(ju1768,* C1022G) | Ectopic foci formation in the proximal neurite | Partial loss | <1% animals | Present |
| *him-4*(ju1689, Y4619STOP) | Ectopic foci formation in the proximal neurite | Completely lost | <1% animals | Present |
